# Quantitative dissection of transcription in development yields evidence for transcription factor-driven chromatin accessibility

**DOI:** 10.1101/2020.01.27.922054

**Authors:** Elizabeth Eck, Jonathan Liu, Maryam Kazemzadeh-Atoufi, Sydney Ghoreishi, Shelby Blythe, Hernan G. Garcia

**Author notes:** These authors contributed equally (Elizabeth Eck, Jonathan Liu).

## Abstract

Thermodynamic models of gene regulation can predict transcriptional regulation in bacteria, but in eukaryotes chromatin accessibility and energy expenditure may call for a different framework. Here we systematically tested the predictive power of models of DNA accessibility based on the Monod-Wyman-Changeux (MWC) model of allostery, which posits that chromatin fluctuates between accessible and inaccessible states. We dissected the regulatory dynamics of *hunchback* by the activator Bicoid and the pioneer-like transcription factor Zelda in living *Drosophila* embryos and showed that no thermodynamic or non-equilibrium MWC model can recapitulate *hunchback* transcription. Therefore, we explored a model where DNA accessibility is not the result of thermal fluctuations but is catalyzed by Bicoid and Zelda, possibly through histone acetylation, and found that this model can predict *hunchback* dynamics. Thus, our theory-experiment dialogue uncovered potential molecular mechanisms of transcriptional regulatory dynamics, a key step toward reaching a predictive understanding of developmental decision-making.

## 1. Introduction

Over the last decade, hopeful analogies between genetic and electronic circuits have posed the challenge of predicting the output gene expression of a DNA regulatory sequence in much the same way that the output current of an electronic circuit can be predicted from its wiring diagram (Endy, 2005). This challenge has been met with a plethora of theoretical works, including thermodynamic models, which use equilibrium statistical mechanics to calculate the probability of finding transcription factors bound to DNA and to relate this probability to the output rate of mRNA production (Ackers et al., 1982; Buchler et al., 2003; Vilar and Leibler, 2003; Bolouri and Davidson, 2003; Bintu et al., 2005b,a; Sherman and Cohen, 2012). Thermodynamic models of bacterial transcription launched a dialogue between theory and experiments that has largely confirmed their predictive power for several operons (Ackers et al., 1982; Bakk et al., 2004; Zeng et al., 2010; He et al., 2010; Garcia and Phillips, 2011; Brewster et al., 2012; Cui et al., 2013; Brewster et al., 2014; Sepulveda et al., 2016; Razo-Mejia et al., 2018) with a few potential exceptions (Garcia et al., 2012; Hammar et al., 2014).

Following these successes, thermodynamic models have been widely applied to eukaryotes to describe transcriptional regulation in yeast (Segal et al., 2006; Gertz et al., 2009; Sharon et al., 2012; Zeigler and Cohen, 2014), human cells (Giorgetti et al., 2010), and the fruit fly *Drosophila melanogaster* (Jaeger et al., 2004a; Zinzen et al., 2006; Segal et al., 2008; Fakhouri et al., 2010; Parker et al., 2011; Kanodia et al., 2012; White et al., 2012; Samee et al., 2015; Sayal et al., 2016). However, two key differences between bacteria and eukaryotes cast doubt on the applicability of thermodynamic models to predict transcriptional regulation in the latter. First, in eukaryotes, DNA is tightly packed in nucleosomes and must become accessible in order for transcription factor binding and transcription to occur (Polach and Widom, 1995; Levine, 2010; Schulze and Wallrath, 2007; Lam et al., 2008; Li et al., 2011; Fussner et al., 2011; Bai et al., 2011; Li et al., 2014a; Hansen and O’Shea, 2015). Second, recent reports have speculated that, unlike in bacteria, the equilibrium framework may be insufficient to account for the energy-expending steps involved in eukaryotic transcriptional regulation, such as histone modifications and nucleosome remodeling, calling for non-equilibrium models of transcriptional regulation (Kim and O’Shea, 2008; Estrada et al., 2016; Li et al., 2018; Park et al., 2019).

Recently, various theoretical models have incorporated chromatin accessibility and energy expenditure in theoretical descriptions of eukaryotic transcriptional regulation. First, models by Mirny (2010), Narula and Igoshin (2010), and Marzen et al. (2013) accounted for chromatin occluding transcription-factor binding by extending thermodynamic models to incorporate the MonodWyman-Changeux (MWC) model of allostery (Fig. 1A; Monod et al., 1965). This thermodynamic MWC model assumes that chromatin rapidly transitions between accessible and inaccessible states via thermal fluctuations, and that the binding of transcription factors to accessible DNA shifts this equilibrium toward the accessible state. Like all thermodynamic models, this model relies on the “occupancy hypothesis” (Hammar et al., 2014; Garcia et al., 2012; Phillips et al., 2019): the probability *p_bound_* of finding RNA polymerase (RNAP) bound to the promoter, a quantity that can be easily computed, is linearly related to the rate of mRNA production 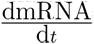, a quantity that can be experimentally measured, such that

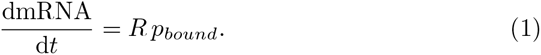

**Figure 1:**
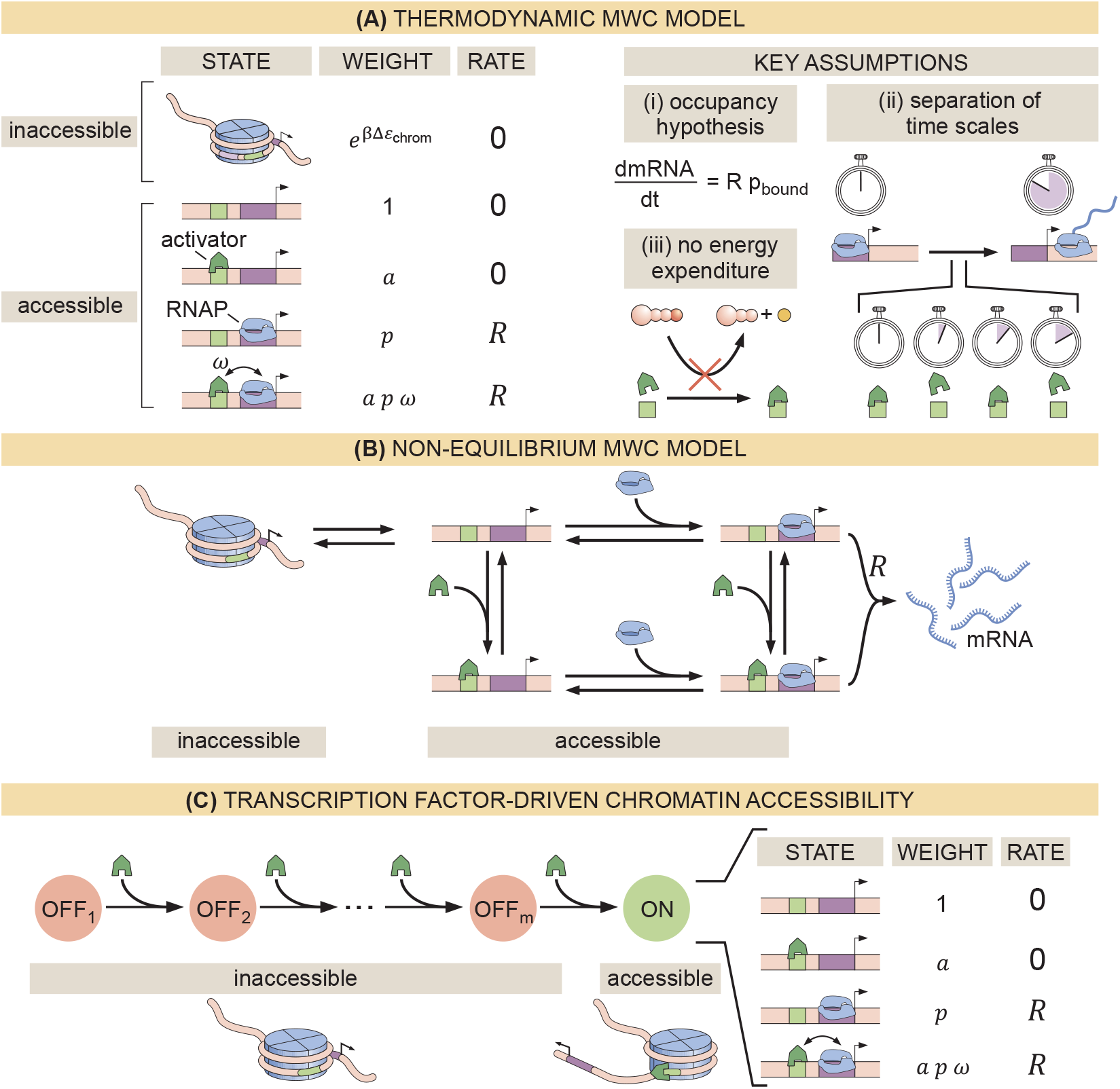
Three models of chromatin accessibility and transcriptional regulation. (A) Thermodynamic MWC model where chromatin can be inaccessible or accessible to transcription factor binding. Each state is associated with a statistical weight given by the Boltzmann distribution and with a rate of transcriptional initiation. Δ*ε*_chrom_ is the energy cost associated with making the DNA accessible and *ω* is an interaction energy between the activator and RNAP. *a* = [activator]*/K_a_* and *p* = [RNAP]*/K_p_* with *K_a_* and *K_p_* being the dissociation constants of the activator and RNAP, respectively. This model assumes the occupancy hypothesis, separation of time scales, and lack of energy expenditure described in the text. (B) Non-equilibrium MWC model where no assumptions about separation of time scales or energy expenditure are made. Transition rates that depend on the concentration of the activator or RNAP are indicated by an arrow incorporating the respective protein. (C) Transcription factor-driven chromatin accessibility model where the activator catalyzes irreversible transitions of the DNA through *m* silent states before it becomes accessible. Once this accessible state is reached, the system is in equilibrium.

Here, *R* is the rate of mRNA production when the system is in an RNAP-bound state (see Section S1.1 for a more detailed overview). Additionally, in all thermodynamic models, the transitions between states are assumed to be much faster than both the rate of transcriptional initiation and changes in transcription factor concentrations. This separation of time scales, combined with a lack of energy dissipation in the process of regulation, makes it possible to consider the states to be in equilibrium such that the probability of each state can be computed using its Boltzmann weight (Garcia et al., 2007).

Despite the predictive power of thermodynamic models, eukaryotic transcription may not adhere to the requirements imposed by the thermodynamic framework. Indeed, Narula and Igoshin (2010), Hammar et al. (2014), Estrada et al. (2016), Scholes et al. (2017), and Li et al. (2018) have proposed theoretical treatments of transcriptional regulation that maintain the occupancy hypothesis, but make no assumptions about separation of time scales or energy expenditure in the process of regulation. When combined with the MWC mechanism of DNA allostery, these models result in a non-equilibrium MWC model (Fig. 1B). Here, no constraints are imposed on the relative values of the transition rates between states and energy can be dissipated over time. To our knowledge, neither the thermodynamic MWC model nor the non-equilibrium MWC model have been tested experimentally in eukaryotic transcriptional regulation.

Here, we performed a systematic dissection of the predictive power of these MWC models of DNA allostery in the embryonic development of the fruit fly *Drosophila melanogaster* in the context of the step-like activation of the *hunch-back* gene by the Bicoid activator and the pioneer-like transcription factor Zelda (Driever et al., 1989; Nien et al., 2011; Xu et al., 2014). Specifically, we compared the predictions from these MWC models against dynamical measurements of input Bicoid and Zelda concentrations and output *hunchback* transcriptional activity. Using this approach, we discovered that no thermodynamic or nonequilibrium MWC model featuring the regulation of *hunchback* by Bicoid and Zelda could describe the transcriptional dynamics of this gene. We proposed a model in which Bicoid and Zelda, rather than passively biasing thermal fluctuations of chromatin toward the accessible state, actively assist the overcoming of an energetic barrier to make chromatin accessible through the recruitment of energy-consuming histone modifiers or chromatin remodelers. This model (Fig. 1C) recapitulated all of our experimental observations. This interplay between theory and experiment establishes a clear path to identify the molecular steps that make DNA accessible, to systematically test our model of transcription factor-driven chromatin accessibility, and to make progress toward a predictive understanding of transcriptional regulation in development.

## 2. Results

### 2.1 A thermodynamic MWC model of activation and chromatin accessibility by Bicoid and Zelda

During the first two hours of embryonic development, the *hunchback* P2 minimal enhancer (Margolis et al., 1995; Driever et al., 1989; Perry et al., 2012; Park et al., 2019) is believed to be devoid of significant input signals other than activation by Bicoid and regulation of chromatin accessibility by both Bicoid and Zelda (Perry et al., 2012; Xu et al., 2014; Hannon et al., 2017). As a result, the early regulation of *hunchback* provides an ideal scaffold for a stringent test of simple theoretical models of eukaryotic transcriptional regulation.

Our implementation of the thermodynamic MWC model (Fig. 1A) in the context of *hunchback* states that in the inaccessible state, neither Bicoid nor Zelda can bind DNA. In the accessible state, DNA is unwrapped and the binding sites become accessible to these transcription factors. Due to the energetic cost of opening the chromatin (Δ*ε*_chrom_), the accessible state is less likely to occur than the inaccessible one. However, the binding of Bicoid or Zelda can shift the equilibrium toward the accessible state (Adams and Workman, 1995; Miller and Widom, 2003; Mirny, 2010; Narula and Igoshin, 2010; Marzen et al., 2013).

In our model, we assume that all binding sites for a given molecular species have the same binding affinity. Relaxing this assumption does not affect any of our conclusions (as we will see below in Sections 2.3 and 2.4). Bicoid upregulates transcription by recruiting RNAP through a protein-protein interaction characterized by the parameter *ω_bp_*. We allow cooperative protein-protein interactions between Bicoid molecules, described by *ω_b_*. However, since to our knowledge there is no evidence of direct interaction between Zelda and any other proteins, we assume no interaction between Zelda and Bicoid, or between Zelda and RNAP.

In Fig. 2A, we illustrate the simplified case of two Bicoid binding sites and one Zelda binding site, plus the corresponding statistical weights of each state given by their Boltzmann factors. Note that the actual model utilized throughout this work accounts for at least six Bicoid binding sites and ten Zelda binding sites that have been identified within the *hunchback* P2 enhancer (Section 4.1; Driever and Nusslein-Volhard, 1988b; Driever et al., 1989; Park et al., 2019). This general model is described in detail in Section S1.2.

**Figure 2:**
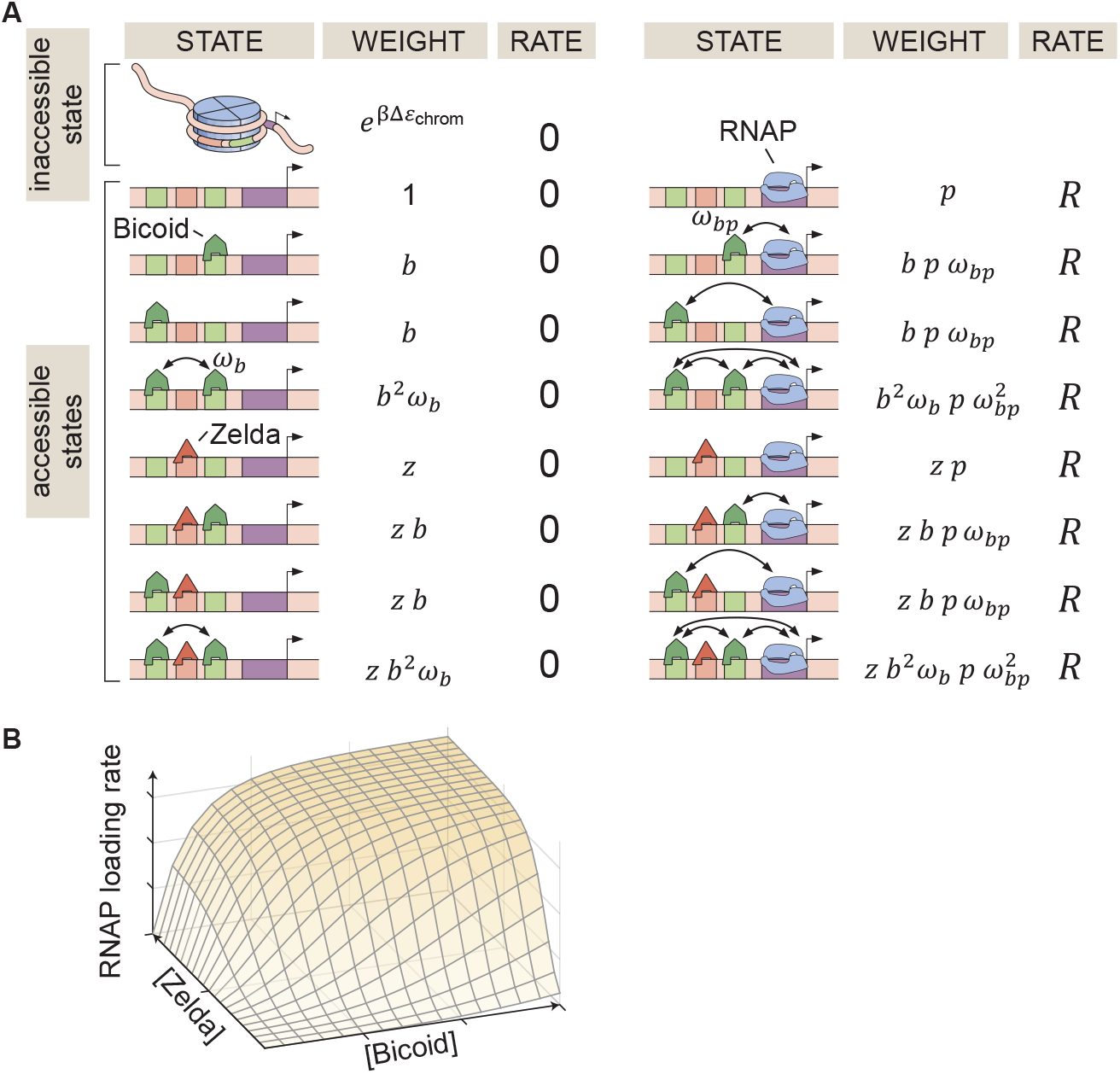
Thermodynamic MWC model of transcriptional regulation by Bicoid and Zelda. (A) States and statistical weights for a simplified version of the *hunchback* P2 enhancer. In this model, we assume that chromatin occluded by nucleosomes is not accessible to transcription factors or RNAP. Parameters are defined in the text. (B) 3D input-output function predicting the rate of RNAP loading (and of transcriptional initiation) as a function of Bicoid and Zelda concentrations for a given set of model parameters.

The probability of finding RNAP bound to the promoter is calculated by dividing the sum of all statistical weights featuring RNAP by the sum of the weights corresponding to all possible system states. This leads to

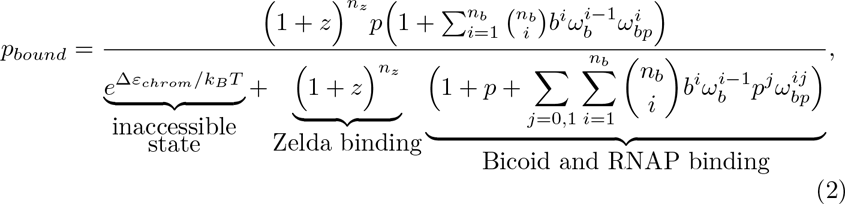

where *b* = [*Bicoid*]*/K_b_*, *z* = [*Zelda*]*/K_z_*, and *p* = [*RNAP*]*/K_p_*, with [*Bicoid*], [*Zelda*], and [*RNAP*] being the concentrations of Bicoid, Zelda, and RNAP, respectively, and *K_b_*, *K_z_*, and *K_p_* their dissociation constants (see Sections S1.1 and S1.2 for a detailed derivation). Given a set of model parameters, plugging *p_bound_* into Equation 1 predicts the rate of RNAP loading as a function of Bicoid and Zelda concentrations as shown in Fig. 2B. Note that in this work, we treat the rate of transcriptional initiation and the rate of RNAP loading interchangeably.

### 2.2. Dynamical prediction and measurement of input-output functions in development

In order to experimentally test the theoretical model in Fig. 2, it is necessary to measure both the inputs – the concentrations of Bicoid and Zelda – as well as the output rate of RNAP loading. Typically, when testing models of transcriptional regulation in bacteria and eukaryotes, input transcription-factor concentrations are assumed to not be modulated in time: regulation is in steady state (Ackers et al., 1982; Bakk et al., 2004; Segal et al., 2008; Garcia and Phillips, 2011; Sherman and Cohen, 2012; Cui et al., 2013; Little et al., 2013; Raveh-Sadka et al., 2009; Sharon et al., 2012; Zeigler and Cohen, 2014; Xu et al., 2015; Sepulveda et al., 2016; Estrada et al., 2016; Razo-Mejia et al., 2018; Zoller et al., 2018; Park et al., 2019). However, embryonic development is a highly dynamic process in which the concentrations of transcription factors are constantly changing due to their nuclear import and export dynamics, and due to protein production, diffusion, and degradation (Edgar and Schubiger, 1986; Edgar et al., 1987; Jaeger et al., 2004b; Gregor et al., 2007b). As a result, it is necessary to go beyond steady-state assumptions and to predict and measure how the *instantaneous*, time-varying concentrations of Bicoid and Zelda at each point in space dictate *hunchback* output transcriptional dynamics.

In order to quantify the concentration dynamics of Bicoid, we utilized an established Bicoid-eGFP line (Sections 4.2, 4.4, and 4.5; Figs. 3A and S3A; Video S1; Gregor et al., 2007b; Liu et al., 2013). As expected, this line displayed the exponential Bicoid gradient across the length of the embryo (Section S2.1; Fig. S3B). We measured mean Bicoid nuclear concentration dynamics along the anterior-posterior axis of the embryo, as exemplified for two positions in Fig. 3A. As previously reported (Gregor et al., 2007b), after anaphase and nuclear envelope formation, the Bicoid nuclear concentration quickly increases as a result of nuclear import. These measurements were used as inputs into the theoretical model in Fig. 2.

**Figure 3:**
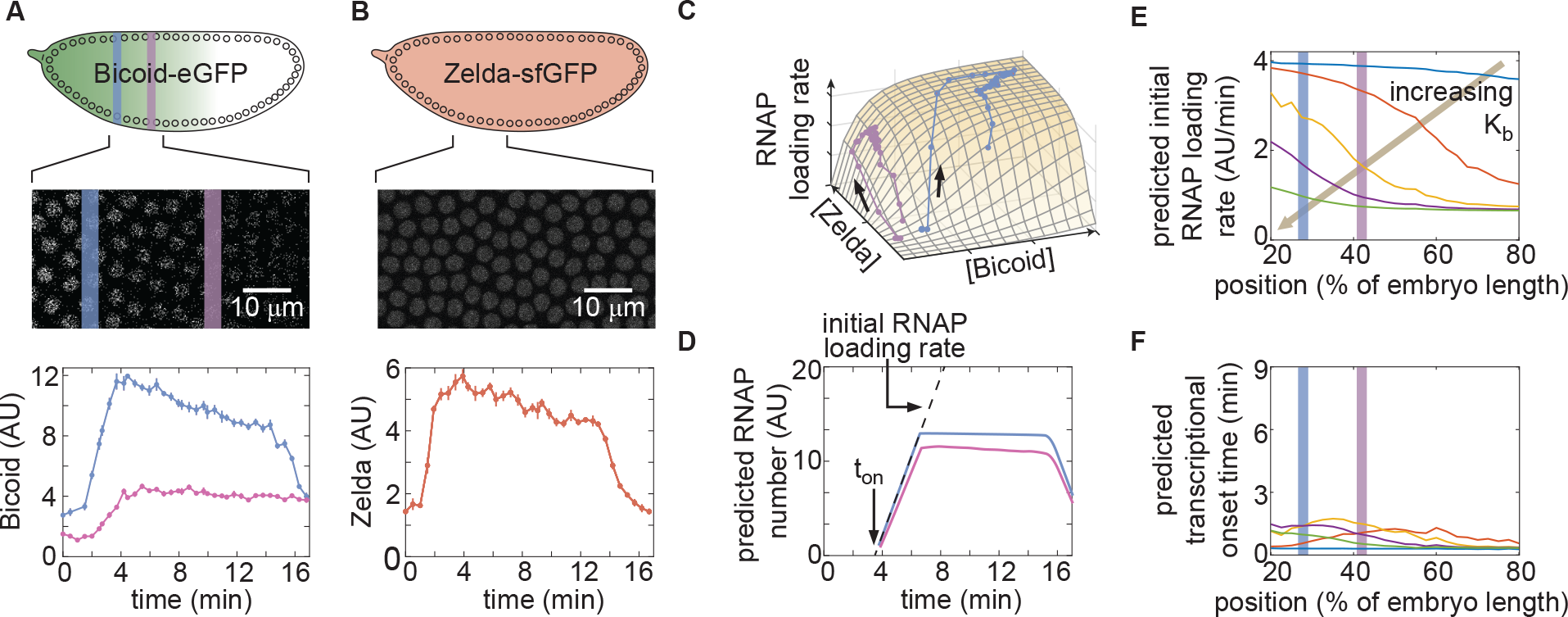
Prediction and measurement of dynamical input-output functions. (A) Measurement of Bicoid concentration dynamics in nuclear cycle 13. Color denotes different positions along the embryo and time is defined with respect to anaphase. (B) Zelda concentration dynamics. These dynamics are uniform throughout the embryo. (C) Trajectories defined by the input concentration dynamics of Bicoid and Zelda along the predicted input-output surface. Each trajectory corresponds to the RNAP loading-rate dynamics experienced by nuclei at the positions indicated in (A). (D) Predicted number of RNAP molecules actively transcribing the gene as a function of time and position along the embryo, and calculation of the corresponding initial rate of RNAP loading and the time of transcriptional onset, *t_on_*. (E,F) Predicted *hunchback* (E) initial rate of RNAP loading and (F) *t_on_* as a function of position along the embryo for varying values of the Bicoid dissociation constant *K_b_*. (A, B, error bars are standard error of the mean nuclear fluorescence in an individual embryo, averaged across all nuclei at a given position; D, the standard error of the mean predicted RNAP number in a single embryo, propagated from the errors in A and B, is thinner than the curve itself; E, F, only mean predictions are shown so as to not obscure differences between them; we imaged n=6 Bicoid-GFP and n=3 Zelda-GFP embryos.)

Zelda concentration dynamics were measured in a Zelda-sfGFP line (Sections 4.2, 4.4, and 4.5; Video S2; Hamm et al., 2017). Consistent with previous results (Staudt et al., 2006; Liang et al., 2008; Dufourt et al., 2018), the Zelda concentration was spatially uniform along the embryo (Fig. S3). Contrasting Figs. 3A and B reveals that the overall concentration dynamics of both Bicoid and Zelda are qualitatively comparable. As a result of Zelda’s spatial uniformity, we used mean Zelda nuclear concentration dynamics averaged across all nuclei within the field of view to test our model (Section S2.1; Fig. 3B).

Given the high reproducibility of the concentration dynamics of Bicoid and Zelda (Fig. S3), we combined measurements from multiple embryos by synchronizing their anaphase in order to create an “averaged embryo” (Section S2.1), an approach that has been repeatedly used to describe protein and transcriptional dynamics in the early fly embryo (Garcia et al., 2013; Bothma et al., 2014, 2015; Berrocal et al., 2018; Lammers et al., 2019).

Our model assumes that *hunchback* output depends on the instantaneous concentration of input transcription factors. As a result, at each position along the anterior-posterior axis of the embryo, the combined Bicoid and Zelda concentration dynamics define a trajectory over time along the predicted input-output function surface (Fig. 3C). The resulting trajectory predicts the rate of RNAP loading as a function of time. However, instead of focusing on calculating RNAP loading rate, we used it to compute the number of RNAP molecules actively transcribing *hunchback* at each point in space and time, a more experimentally accessible quantity (Section 2.3). This quantity can be obtained by accounting for the RNAP elongation rate and the cleavage of nascent RNA upon termination (Section S2.2; Fig. S4; Bothma et al., 2014; Lammers et al., 2019) yielding the predictions shown in Fig. 3D.

Instead of examining the full time-dependent nature of our data, we analyzed two main dynamical features stemming from our prediction of the number of RNAP molecules actively transcribing *hunchback*: the initial rate of RNAP loading and the transcriptional onset time, *t_on_*, defined by the slope of the initial rise in the predicted number of RNAP molecules, and the time after anaphase at which transcription starts as determined by the x-intercept of the linear fit to the initial rise, respectively (Fig. 3D).

Examples of the predictions generated by our theoretical model are shown in Fig. 3E and F, where we calculate the initial rate of RNAP loading and *t_on_* for different values of the Bicoid dissociation constant *K_b_*. This framework for quantitatively investigating dynamic input-output functions in living embryos is a necessary step toward testing the predictions of theoretical models of transcriptional regulation in development.

### 2.3. The thermodynamic MWC model fails to predict activation of hunchback in the absence of Zelda

In order to test the predictions of the thermodynamic MWC model (Fig. 3E and F), we used the MS2 system (Bertrand et al., 1998; Garcia et al., 2013; Lucas et al., 2013). Here, 24 repeats of the MS2 loop are inserted in the 5’ untranslated region of the *hunchback* P2 reporter (Garcia et al., 2013), resulting in the fluorescent labeling of sites of nascent transcript formation (Fig. 4A; Video S3). This fluorescence is proportional to the number of RNAP molecules actively transcribing the gene (Garcia et al., 2013). The experimental mean fluorescence as a function of time measured in a narrow window (2.5% of the total embryo length, averaged across nuclei in the window) along the length of the embryo (Fig. 4B) is in qualitative agreement with the theoretical prediction (Fig. 3D).

**Figure 4:**
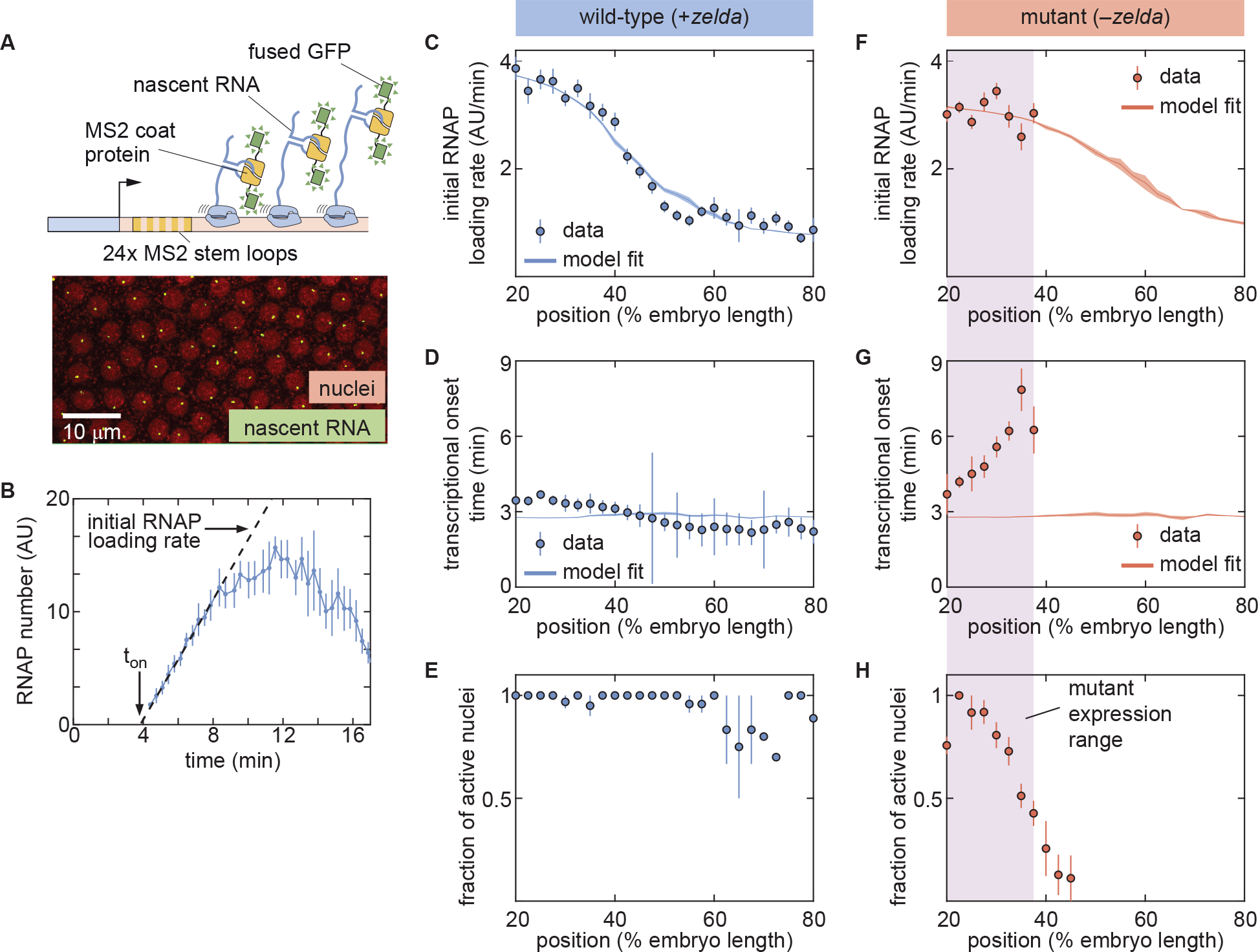
The thermodynamic MWC model can explain *hunchback* transcriptional dynamics in wild-type, but not *zelda^−^*, embryos. (A) The MS2 system measures the number of RNAP molecules actively transcribing the *hunchback* reporter gene in live embryos. (B) Representative MS2 trace featuring the quantification of the initial rate of RNAP loading and *t_on_*. (C) Initial RNAP loading rate and (D) *t_on_*for wild-type embryos (points), compared with best fit to the thermodynamic MWC model (lines). (E) Fraction of transcriptionally active nuclei for wild-type embryos. Active nuclei are defined as nuclei that exhibited an MS2 spot at any time during the nuclear cycle. (F) Initial RNAP loading rate and (G) *t_on_* for *zelda^−^*embryos (points), compared with best fit to the thermodynamic MWC model (lines). (H) Fraction of transcriptionally active nuclei for *zelda^−^*embryos. Purple shading indicates the spatial range over which at least 50% of nuclei display transcription. (B, error bars are standard error of the mean observed RNAP number, averaged across nuclei in a single embryo; C,D,F,G, solid lines indicate mean predictions of the model, shading represents standard error of the mean; C,D,E, error bars in data points represent standard error of the mean over 8 embryos; F,G,H, error bars in data points represent standard error of the mean over 14 embryos.)

To compare theory and experiment, we next obtained the initial RNAP loading rates (Fig. 4C, points) and *t_on_* (Fig. 4D, points) from the experimental data (Section S2.3; Fig. S5B). The step-like shape of the RNAP loading rate (Fig. 4C) agrees with previous measurements performed on this same reporter construct (Garcia et al., 2013). The plateaus at the extreme anterior and posterior positions were used to constrain the maximum and minimum theoretically allowed values in the model (Section S1.3). With these constraints in place, we attempted to simultaneously fit the thermodynamic MWC model to both the initial rate of RNAP loading and *t_on_*. For a given set of model parameters, the measurements of Bicoid and Zelda concentration dynamics predicted a corresponding initial rate of RNAP loading and *t_on_* (Fig. 3E and F). The model parameters were then iterated using standard curve-fitting techniques (Section 4.6) until the best fit to the experimental data was achieved (Fig. 4C and D, lines).

Although the model accounted for the initial rate of RNAP loading (Fig. 4C, line), it produced transcriptional onset times that were much lower than those that we experimentally observed (Fig. S6B, purple line). We hypothesized that this disagreement was due to our model not accounting for mitotic repression, when the transcriptional machinery appears to be silent immediately after cell division (Shermoen and O’Farrell, 1991; Gottesfeld and Forbes, 1997; Parsons and Tg, 1997; Garcia et al., 2013). Thus, we modified the thermodynamic MWC model to include a mitotic repression window term, implemented as a time window at the start of the nuclear cycle during which no transcription could occur; the rate of mRNA production is thus given by

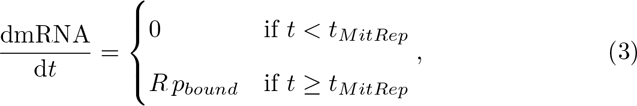

where *R* and *p_bound_* are as defined in Eqns. 1 and 2, respectively, and *t_MitRep_*is the mitotic repression time window over which no transcription can take place after anaphase (Sections S1.2 and S3.1). After incorporating mitotic repression, the thermodynamic MWC model successfully fit both the rates of RNAP loading and *t_on_* (Fig. 4C and D, lines, Fig. S6A and B).

Given this success, we next challenged the model to perform the simpler task of explaining Bicoid-mediated regulation in the absence of Zelda. This scenario corresponds to setting the concentration of Zelda to zero in the models in Section S1.2 and Fig. 2. In order to test this seemingly simpler model, we repeated our measurements in embryos devoid of Zelda protein (Video S4). These *zelda^−^*embryos were created by inducing clones of non-functional *zelda* mutant (*zelda*^294^) germ cells in female adults (Sections 4.2, 4.3; Liang et al., 2008). All embryos from these mothers lack maternally deposited Zelda; female embryos still have a functional copy of *zelda* from their father, but this copy is not transcribed until after the maternal-to-zygotic transition, during nuclear cycle 14 (Liang et al., 2008). We confirmed that the absence of Zelda did not have a substantial effect on the spatiotemporal pattern of Bicoid (Section S4.1; Xu et al., 2014).

While close to 100% of nuclei in wild-type embryos exhibited transcription along the length of the embryo (Fig. 4E; Video S5), measurements in the *zelda^−^* background revealed that some nuclei never displayed any transcription during the entire nuclear cycle (Video S6). Specifically, transcription occurred only in the anterior part of the embryo, with transcription disappearing completely in positions posterior to about 40% of the embryo length (Fig. 4H). From those positions in the mutant embryos that did exhibit transcription in at least 50% of observed nuclei, we extracted the initial rate of RNAP loading and *t_on_* as a function of position. Interestingly, these RNAP loading rates were comparable to the corresponding rates in wild-type embryos (Fig. 4F, points). However, unlike in the wild-type case (Fig. 4D, points), *t_on_* was not constant in the *zelda ^−^* background. Instead, *t_on_* became increasingly delayed in more posterior positions until transcription ceased posterior to 40% of the embryo length (Fig. 4G, points). Together, these observations indicated that removing Zelda primarily results in a delay of transcription with only negligible effects on the underlying rates of RNAP loading, consistent with previous fixed-embryo experiments (Nien et al., 2011; Foo et al., 2014) and with recent live-imaging measurements in which Zelda binding was reduced at specific enhancers (Dufourt et al., 2018; Yamada et al., 2019). We speculate that the loss of transcriptionally active nuclei posterior to 40% of the embryo length is a direct result of this delay in *t_on_*: by the time that onset would occur in those nuclei, the processes leading to the next mitosis have already started and repressed transcriptional activity.

Next, we attempted to simultaneously fit the model to the initial rates of RNAP loading and *t_on_* in the *zelda^−^* mutant background. Although the model recapitulated the observed initial RNAP loading rates, we noticed a discrepancy between the observed and fitted transcriptional onset times of up to *∼*5 min (Fig. 4F and G). While the mutant data exhibited a substantial delay in more posterior nuclei, the model did not produce significant delays (Fig. 4G, line). Further, our model could not account for the lack of transcriptional activity posterior to 40% of the embryo length in the *zelda^−^* mutant (Fig. 4H).

These discrepancies suggest that the thermodynamic MWC model cannot fully describe the transcriptional regulation of the *hunchback* promoter by Bicoid and Zelda. However, the attempted fits in Fig. 4F and G correspond to a particular set of model parameters and therefore do not completely rule out the possibility that there exists some parameter set of the thermodynamic MWC model capable of recapitulating the *zelda^−^* data.

In order to determine whether this model is *at all* capable of accounting for the *zelda^−^* transcriptional behavior, we systematically explored how its parameters dictate its predictions. To characterize and visualize the limits of our model, we examined a mathematically convenient metric: the average transcriptional onset delay along the anterior-posterior axis (Fig. 5A). This quantity is defined as the area under the curve of *t_on_* versus embryo position, from 20% to 37.5% along the embryo (the positions where the *zelda^−^* data display transcription), divided by the corresponding distance along the embryo

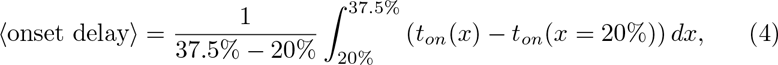

where *x* is the position along the embryo and the value of *t_on_* at 20% along the embryo was chosen as the offset with respect to which to define the zero of this integral (Section S5.1). The average *t_on_* delay corresponding to the wildtype data is close to zero, and is substantially different from the larger value obtained from measurements in the *zelda^−^* background within experimental error (Fig. 5C, points).

**Figure 5:**
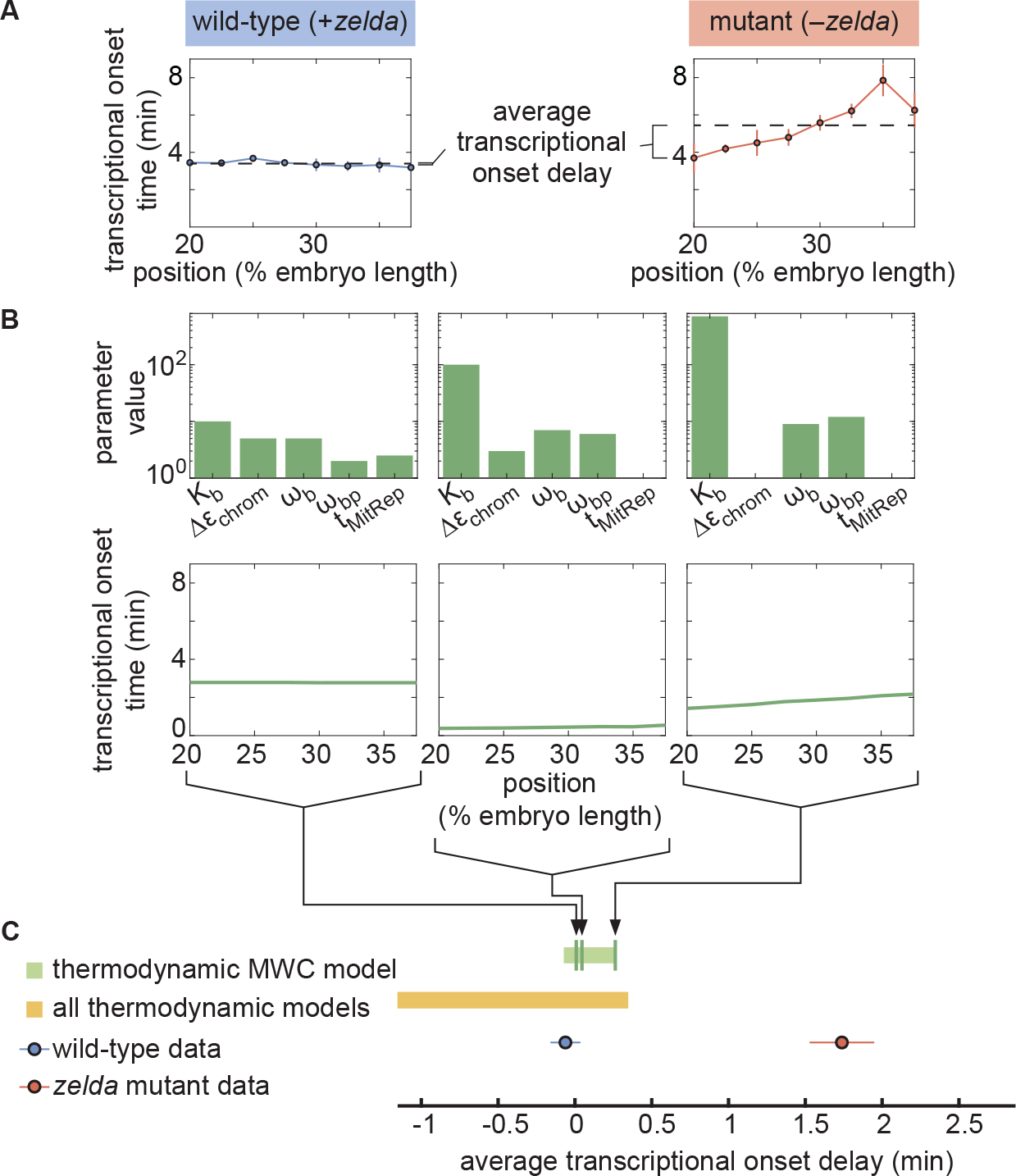
Failure of thermodynamic models to describe Bicoid-dependent activation of *hunchback*. (A) Experimentally determined *t_on_*. Horizontal dashed lines indicate the average *t_on_* delay with respect to *t_on_*(*x* = 20%) for wild-type and *zelda^−^* data sets. (B) Exploration of average *t_on_* delay from the thermodynamic MWC model. Each choice of model parameters predicts a distinct *t_on_* profile of along the embryo. (C) Predicted range of average *t_on_* delay for all possible parameter choices of the thermodynamic MWC model (green), as well as for all thermodynamic models considering 12 Bicoid binding sites (yellow), compared with experimental data (red, blue). (A,C, error bars represent standard error of the mean over 8 and 14 embryos for the wild-type and *zelda^−^*datasets, respectively; B, solid lines indicate mean predictions of the model)

Based on Estrada et al. (2016) and as detailed in Section S5.1, we used an algorithm to efficiently sample the parameter space of the thermodynamic MWC model (dissociation constants *K_b_* and *K_z_*, protein-protein interaction terms *ω_b_* and *ω_bp_*, energy to make the DNA accessible Δ*ε*_chrom_, and length of the mitotic repression window *t_MitRep_*), and to calculate the corresponding average *t_on_* delay for each parameter set. Fig. 5B features three specific realizations of this parameter search; for each combination of parameters considered, the predicted *t_on_* is calculated and the corresponding average *t_on_* delay computed. Although the wild-type data were contained within the thermodynamic MWC model region, the range of the average *t_on_* delay predicted by the model (Fig. 5C, green rectangle) did not overlap with the *zelda^−^* data. We concluded that our thermodynamic MWC model is not sufficient to explain the regulation of *hunchback* by Bicoid and Zelda.

### 2.4. No thermodynamic model can recapitulate the activation of hunchback by Bicoid alone

Since the failure of the thermodynamic MWC model to predict the *zelda^−^* data does not necessarily rule out the existence of another thermodynamic model that can account for our experimental measurements, we considered other possible thermodynamic models. Conveniently, an arbitrary thermodynamic model featuring *n_b_* Bicoid binding sites can be generalized using the mathematical expression

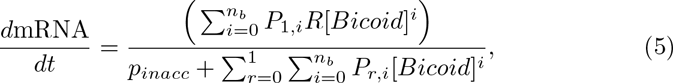

where *p_inacc_* and *P_r,i_* are *arbitrary* weights describing the states in our generalized thermodynamic model, *R* is a rate constant that relates promoter occupancy to transcription rate, and the *r* and *i* summations refer to the numbers of RNAP and Bicoid molecules bound to the enhancer, respectively (Section S6.1; Bintu et al., 2005a; Estrada et al., 2016; Scholes et al., 2017).

Although this generalized thermodynamic model contains many more parameters than the thermodynamic MWC model previously considered, we could still systematically explore these parameters and the resulting average *t_on_* delays. For added generality, and to account for recent reports suggesting the presence of more than six Bicoid binding sites in the *hunchback* minimal enhancer (Park et al., 2019), we expanded this model to include up to 12 Bicoid binding sites. Even though the generalized thermodynamic model occupied a larger region of the average *t_on_* delay space than the thermodynamic MWC model, it still failed to explain the *zelda^−^* data (Section S6.2; Fig. 5C, yellow rectangle). Thus, our results strongly suggest that no thermodynamic model of Bicoid-activated *hunchback* transcription can predict transcriptional onset in the absence of Zelda, casting doubt on the general applicability of these models to transcriptional regulation in development.

Qualitatively, the reason for the failure of thermodynamic models to predict *hunchback* transcriptional is revealed by comparing Bicoid and Zelda concentration dynamics to those of the MS2 output signal (Fig. S10). The thermodynamic models investigated in this work have assumed that the system responds *instantaneously* to any changes in input transcription factor concentration. As a result, since Bicoid and Zelda are imported into the nucleus by around 3 min into the nuclear cycle (Fig. 3A and B), these models always predict that transcription will ensue at approximately that time. Thus, thermodynamic models cannot accommodate delays in the *t_on_* such as those revealed by the *zelda^−^* data (see Section S6.3 for a more detailed explanation). Rather than further complicating our thermodynamic models with additional molecular players to attempt to describe the data, we instead decided to examine the broader purview of non-equilibrium models to attempt to reach an agreement between theory and experiment.

### 2.5. A non-equilibrium MWC model also fails to describe the zelda^−^ data

Thermodynamic models based on equilibrium statistical mechanics can be seen as limiting cases of more general kinetic models that lie out of equilibrium (Section S6.4; Fig. 1B). Following recent reports (Estrada et al., 2016; Li et al., 2018; Park et al., 2019) that the theoretical description of transcriptional regulation in eukaryotes may call for models rooted in non-equilibrium processes – where the assumptions of separation of time scales and no energy expenditure may break down – we extended our earlier models to produce a non-equilibrium MWC model (Sections S6.4 and S7.1; Kim and O’Shea, 2008; Narula and Igoshin, 2010). This model, shown for the case of two Bicoid binding sites in Fig. 6A, accounts for the dynamics of the MWC mechanism by positing transition rates between the inaccessible and accessible chromatin states, but makes no assumptions about the relative magnitudes of these rates, or about the rates of Bicoid and RNAP binding and unbinding.

**Figure 6:**
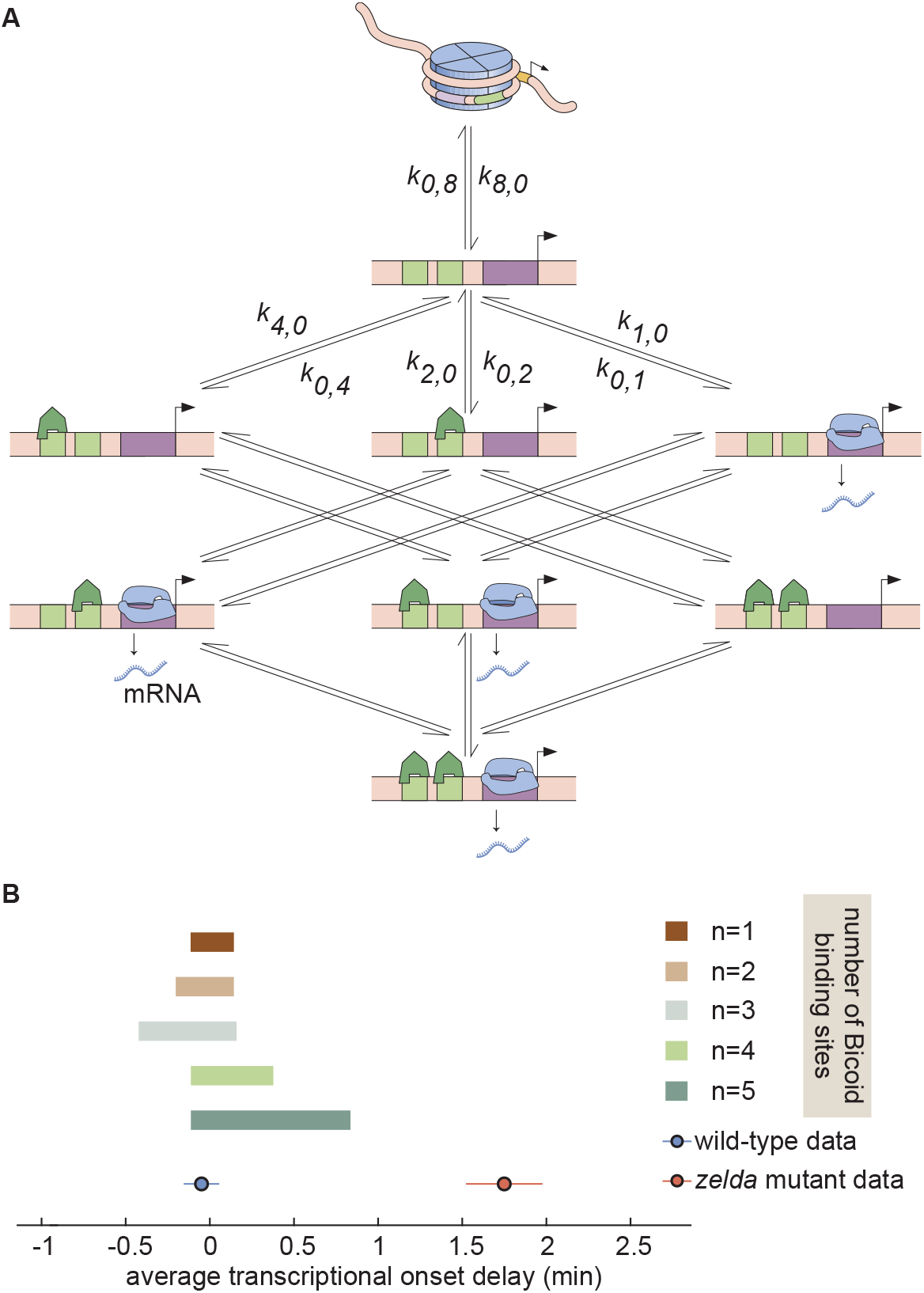
Non-equilibrium MWC model of transcriptional regulation cannot predict the observed *t_on_* delay. (A) Model that makes no assumptions about the relative transition rates between states or about energy expenditure. Each transition rate *i, j* represents the rate of switching from state *i* to state *j*. See Section S7.1 for details on how the individual states are labeled. (B) Exploration of average *t_on_* delay attainable by the non-equilibrium MWC models as a function of the number of Bicoid binding sites compared to the experimentally obtained values corresponding to the wild-type and *zelda^−^* mutant backgrounds. While the non-equilibrium MWC model can explain the wild-type data, the exploration reveals that it fails to explain the*zelda^−^* data, for up to five Bicoid binding sites.

Since this model can operate out of steady state, we calculate the probabilities of each state as a function of time by solving the system of coupled ordinary differential equations (ODEs) associated with the system shown in Fig. 6A. Consistent with prior measurements (Blythe and Wieschaus, 2016), we assume that chromatin is inaccessible at the start of the nuclear cycle. Over time, the system evolves such that the probability of it occupying each state becomes nonzero, making it possible to calculate the fraction of time RNAP is bound to the promoter and, through the occupancy hypothesis, the rate of RNAP loading.

We systematically varied the magnitudes of the transition rates and solved the system of ODEs in order to calculate the corresponding average *t_on_* delay. Due to the combinatorial increase of free parameters as more Bicoid binding sites are included in the model, we could only explore the parameter space for models containing up to five Bicoid binding sites (Section S7.2; Figs. 6B and S9). Interestingly, while the upper bound of the average *t_on_* delay monotonically increased with Bicoid binding site number, the lower bound of this delay did not vary in a systematic fashion (Fig. 6B). This phenomenon where the parameter space of a model does not strictly increase with binding site number has been previously observed (Estrada et al., 2016) and the reason for this effect remains uncertain. Regardless, none of the non-equilibrium MWC models with up to five Bicoid binding sites came close to reaching the mutant average *t_on_* delay (Fig. 6B). We conjecture that the observed behavior extends to the biologically relevant case of six or more binding sites. Thus, we conclude that the more comprehensive non-equilibrium MWC model still cannot account for the experimental data, motivating an additional reexamination of our assumptions.

### 2.6. Transcription factor-driven chromatin accessibility can capture all aspects of the data

Since even non-equilibrium MWC models incorporating energy expenditure and non-steady behavior could not explain the *zelda^−^* data, we further revised the assumptions of our model in an effort to quantitatively predict the regulation of *t_on_* along the embryo. In accordance with the MWC model of allostery, all of our theoretical treatments so far have posited that the DNA is an allosteric molecule that transitions between open and closed states as a result of thermal fluctuations (Narula and Igoshin, 2010; Mirny, 2010; Marzen et al., 2013; Phillips et al., 2013).

In the MWC models considered here, the presence of Zelda and Bicoid does not affect the microscopic rates of DNA opening and closing; rather, their binding to open DNA shifts the equilibrium of the DNA conformation toward the accessible state. However, recent biochemical work has suggested that Zelda and Bicoid play a more direct role in making chromatin accessible. Specifically, Zelda has been implicated in the acetylation of chromatin, a histone modification that renders nucleosomes unstable and increases DNA accessibility (Li et al., 2014b; Li and Eisen, 2018). Further, Bicoid has been shown to interact with the co-activator dCBP, which possesses histone acetyltransferase activity (Fu et al., 2004). Additionally, recent studies by Desponds et al. (2016) in *hunchback* and by Dufourt et al. (2018) in *snail* have proposed the existence of multiple transcriptionally silent steps that the promoter needs to transition through before transcriptional onset. These steps could correspond to, for example, the recruitment of histone modifiers, nucleosome remodelers, and the transcriptional machinery (Li et al., 2014b; Park et al., 2019), or to the step-wise unraveling of discrete histone-DNA contacts (Culkin et al., 2017). Further, Dufourt et al. (2018) proposed that Zelda plays a role in modulating the number of these steps and their transition rates.

We therefore proposed a model of transcription factor-driven chromatin accessibility in which, in order for the DNA to become accessible and transcription to ensue, the system slowly and irreversibly transitions through *m* transcriptionally silent states (Section S8.1; Fig. 7A). We assume that the transitions between these states are all governed by the same rate constant *π*. Finally, in a stark deviation from the MWC framework, we posit that these transitions can be catalyzed by the presence of Bicoid and Zelda such that

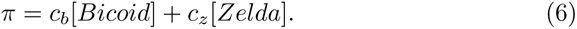

**Figure 7:**
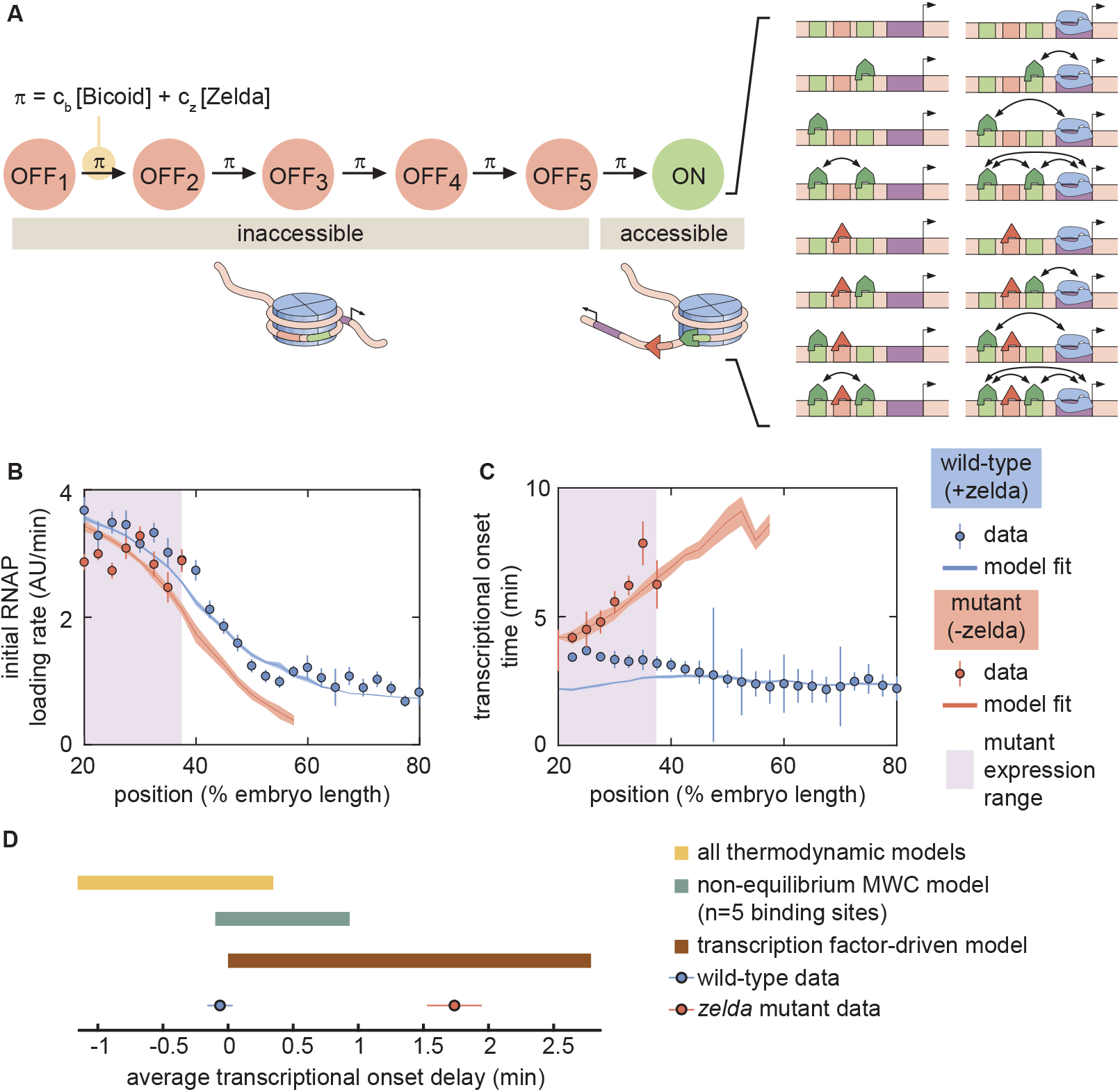
A model of transcription factor-driven chromatin accessibility is sufficient to recapitulate *hunchback* transcriptional dynamics. (A) Overview of the proposed model, with five (*m* = 5) effectively irreversible Zelda and/or Bicoid-mediated intermediate transitions from the inaccessible to the accessible states. (B,C) Experimentally fitted (B) initial RNAP loading rates and (C) *t_on_* for wild-type and *zelda^−^* embryos using a single set of parameters and assuming six Bicoid binding sites. (D) The domain of average *t_on_* delay covered by this transcription factor-driven chromatin accessibility model (brown rectangle) is much larger than those of the generalized thermodynamic model (yellow rectangle) and the non-equilibrium MWC models (green rectangle), and easily encompasses both experimental datasets (points). (B-D, error bars represent standard error of the mean over multiple embryos).

Here, *π* describes the rate (in units of inverse time) of each irreversible step, expressed as a sum of rates that depend separately on the concentrations of Bicoid and Zelda, and *c_b_* and *c_z_* are rate constants that scale the relative contribution of each transcription factor to the overall rate.

In this model of transcription factor-driven chromatin accessibility, once the DNA becomes irreversibly accessible after transitioning through the *m* nonproductive states, we assume that, for the rest of the nuclear cycle, the system equilibrates rapidly such that the probability of it occupying any of its possible states is still described by equilibrium statistical mechanics. Like in our previous models, transcription only occurs in the RNAP-bound states, obeying the occupancy hypothesis. Further, our model assumes that if the transcriptional onset time of a given nucleus exceeds that of the next mitosis, this nucleus will not engage in transcription.

Unlike the thermodynamic and non-equilibrium MWC models, this model of transcription factor-driven chromatin accessibility quantitatively recapitulated the observation that posterior nuclei do not engage in transcription, the initial rate of RNAP loading, and *t_on_* for both the wild-type and *zelda^−^* mutant data (Fig. 7B and C). Additionally, we found that a minimum of *m* = 5 steps was required to sufficiently explain the data (Section S8.2; Fig. S13). Interestingly, unlike all previously considered models, the model of transcription factor-driven chromatin accessibility did not require mitotic repression to explain *t_on_* (Sections S3.1 and S8.1). Instead, the timing of transcriptional output arose directly from the model’s initial irreversible transitions (Fig. S13), obviating the need for an arbitrary suppression window in the beginning of the nuclear cycle. The only substantive disagreement between our theoretical model and the experimental data was that the model predicted that no nuclei should transcribe posterior to 60% of the embryo length, whereas no transcription posterior to 40% was experimentally observed in the embryo (Fig. 7B and C). Finally, note that this model encompasses a much larger region of parameter space than the equilibrium and non-equilibrium MWC models and, as expected from the agreement between model and experiment described above, contained both the wild-type and *zelda^−^* data points within its domain (Fig. 7D).

## 3. Discussion

For four decades, thermodynamic models rooted in equilibrium statistical mechanics have constituted the null theoretical model for calculating how the number, placement and affinity of transcription factor binding sites on regulatory DNA dictates gene expression (Bintu et al., 2005a,b). Further, the MWC mechanism of allostery has been proposed as an extra layer that allows thermodynamic and more general non-equilibrium models to account for the regulation of chromatin accessibility (Mirny, 2010; Narula and Igoshin, 2010; Marzen et al., 2013).

In this investigation, we tested thermodynamic and non-equilibrium MWC models of chromatin accessibility and transcriptional regulation in the context of *hunchback* activation in the early embryo of the fruit fly *D. melanogaster* (Driever et al., 1989; Nien et al., 2011; Xu et al., 2014). While chromatin state (accessibility, post-translational modifications) is highly likely to influence transcriptional dynamics of associated promoters, specifically measuring the influence of chromatin state on transcriptional dynamics is challenging because of the sequential relationship between changes in chromatin state and transcriptional regulation. However, the *hunchback* P2 minimal enhancer provides a unique opportunity to dissect the relative contribution of chromatin regulation on transcriptional dynamics because, in the early embryo, chromatin accessibility at *hunchback* is granted by both Bicoid and Zelda (Hannon et al., 2017). The degree of *hunchback* transcriptional activity, however, is regulated directly by Bicoid (Driever and Nusslein-Volhard, 1989; Driever et al., 1989; Struhl et al., 1989). Therefore, while genetic elimination of Zelda function interferes with acquisition of full chromatin accessibility, the *hunchback* locus retains a measurable degree of accessibility and transcriptional activity stemming from Bicoid function, allowing for a quantitative determination of the contribution of Zeldadependent chromatin accessibility on the transcriptional dynamics of the locus.

With these attributes in mind, we constructed a thermodynamic MWC model which, given a set of parameters, predicted an output rate of *hunch-back* transcription as a function of the input Bicoid and Zelda concentrations (Fig. 2B). In order to test this model, it was necessary to acknowledge that development is not in steady-state, and that both Bicoid and Zelda concentrations change dramatically in space and time (Fig. 3A and B). As a result, we went beyond widespread steady-state descriptions of development and introduced a novel approach that incorporated transient dynamics of input transcription-factor concentrations in order to predict the instantaneous output transcriptional dynamics of *hunchback* (Fig. 3C). Given input dynamics quantified with fluorescent protein fusions to Bicoid and Zelda, we both predicted output transcriptional activity and measured it with an MS2 reporter (Figs. 3D and 4B).

This approach revealed that the thermodynamic MWC model sufficiently predicts the timing of the onset of transcription and the subsequent initial rate of RNAP loading as a function of Bicoid and Zelda concentration. However, when confronted with the much simpler case of Bicoid-only regulation in a *zelda* mutant, the thermodynamic MWC model failed to account for the observations that only a fraction of nuclei along the embryo engaged in transcription, and that the transcriptional onset time of those nuclei that do transcribe was significantly delayed with respect to the wild-type setting (Fig. 4D, E, G, and H). Our systematic exploration of all thermodynamic models (over a reasonable parameter range) showed that that no thermodynamic model featuring regulation by Bicoid alone could quantitatively recapitulate the measurements performed in the *zelda* mutant background (Fig. 5C, yellow rectangle).

This disagreement could be resolved by invoking an unknown transcription factor that regulates the *hunchback* reporter in addition to Bicoid. However, at the early stages of development analyzed here, such a factor would need to be both maternally provided and patterned in a spatial gradient to produce the observed position-dependent transcriptional onset times. To our knowledge, none of the known maternal genes regulate the expression of this *hunchback* reporter in such a fashion (Chen et al., 2012; Perry et al., 2012; Xu et al., 2014). We conclude that the MWC thermodynamic model cannot accurately predict *hunchback* transcriptional dynamics.

To explore non-equilibrium models, we retained the MWC mechanism of chromatin accessibility, but did not demand that the accessible and inaccessible states be in thermal equilibrium. Further, we allowed for the process of Bicoid and RNAP binding, as well as their interactions, to consume energy. For up to five Bicoid binding sites, no set of model parameters could quantitatively account for the transcriptional onset delays in the *zelda* mutant background (Fig. 6B). While we were unable to investigate models with more than five Bicoid binding sites due to computational complexity (Estrada et al., 2016), the substantial distance in parameter space between the mutant data and the investigated models (Fig. 6B) suggested that a successful model with more than five Bicoid binding sites would probably operate near the limits of its explanatory power, similar to the conclusions from studies that explored *hunchback* regulation under the steady-state assumption (Park et al., 2019). Thus, despite the simplicity and success of the MWC model in predicting the effects of protein allostery in a wide range of biological contexts (Keymer et al., 2006; Swem et al., 2008; Martins and Swain, 2011; Marzen et al., 2013; Rapp and Yifrach, 2017; Razo-Mejia et al., 2018; Chure et al., 2019; Rapp and Yifrach, 2019), the observed transcriptional onset times could not be described by any previously proposed thermodynamic MWC mechanism of chromatin accessibility, or even by a more generic non-equilibrium MWC model in which energy is continuously dissipated (Tu, 2008; Kim and O’Shea, 2008; Narula and Igoshin, 2010; Estrada et al., 2016; Wang et al., 2017).

Since Zelda is associated with histone acetylation, which is correlated with increased chromatin accessibility (Li et al., 2014b; Li and Eisen, 2018), and Bicoid interacts with the co-activator dCBP, which has histone acetyltransferase activity (Fu et al., 2004; Fu and Ma, 2005; Park et al., 2019), we suspect that both Bicoid and Zelda actively drive DNA accessibility. A molecular pathway shared by Bicoid and Zelda to render chromatin accessible is consistent with our results, and with recent genome-wide experiments showing that Bicoid can rescue the function of Zelda-dependent enhancers at high enough concentrations (Hannon et al., 2017). Thus, the binding of Bicoid and Zelda, rather than just biasing the equilibrium of the open chromatin state as in the MWC mechanism, may trigger a set of molecular events that locks DNA into an accessible state. In addition, the promoters of *hunchback* (Desponds et al., 2016) and *snail* (Dufourt et al., 2018) may transition through a set of intermediate, non-productive states before transcription begins.

We therefore explored a model in which Bicoid and Zelda catalyze the transition of chromatin into the accessible state via a series of slow, effectively irreversible steps. These steps may be interpreted as energy barriers that are overcome through the action of Bicoid and Zelda, consistent with the coupling of these transcription factors to histone modifiers, nucleosome remodelers (Fu et al., 2004; Li et al., 2014b; Li and Eisen, 2018; Park et al., 2019), and with the step-wise breaking of discrete histone-DNA contacts to unwrap nucleosomal DNA (Culkin et al., 2017). In this model, once accessible, the chromatin remains in that state and the subsequent activation of *hunchback* by Bicoid is described by a thermodynamic model.

Crucially, this transcription factor-driven chromatin accessibility model successfully replicated all of our experimental observations. A minimum of five intermediate transcriptionally silent states were necessary to explain our data (Figs. 7D and S13C). Interestingly, recent work dissecting the transcriptional onset time distribution of *snail* suggested the existence of three such intermediate steps in the context of that gene (Dufourt et al., 2018). These intermediate steps may reveal fundamental constraints for transcriptional regulation in the fruit fly. Intriguingly, accounting for the intermediate states obviated the need for the *ad hoc* imposition of a mitotic repression window (Sections S3.1 and S8.1), which was required in the thermodynamic MWC model (Fig. S6). Our results suggest a mechanistic interpretation of the phenomenon of mitotic repression after anaphase, where the promoter must traverse through intermediary states before transcriptional onset can occur. Finally, given that, as in *hunchback*, the removal and addition of Zelda modulates the timing of transcriptional onset of *sog* and *snail* (Dufourt et al., 2018; Yamada et al., 2019), we speculate that transcription factor-driven chromatin accessibility may also be at play in these pathways.

These clues into the molecular mechanisms of action of Bicoid, Zelda, and their associated modifications to the chromatin landscape pertain to a time scale of a few minutes, a temporal scale that is inaccessible with widespread genome-wide and fixed-tissue approaches. Here, we revealed the regulatory action of Bicoid and Zelda by utilizing the dynamic information provided by live imaging to analyze the transient nature of the transcriptional onset time, highlighting the need for descriptions of development that go beyond steady state and acknowledge the highly dynamic changes in transcription-factor concentrations that drive developmental programs.

While we showed that one model incorporating transcription factor-driven chromatin accessibility could recapitulate *hunchback* transcriptional regulation by Bicoid and Zelda, and is consistent with molecular evidence on the modes of action of these transcription factors, other models may have comparable explanatory power. In the future, a systematic exploration of different classes of models and their unique predictions will identify measurements that determine *which* specific model is the most appropriate description of transcriptional regulation in development and *how* it is implemented at the molecular level. While all the analyses in this work relied on mean levels of input concentrations and output transcription levels, detailed studies of single-cell features of transcriptional dynamics such as the distribution of transcriptional onset times (Narula and Igoshin, 2010; Dufourt et al., 2018) could shed light on these chromatin-regulating mechanisms. Simultaneous measurement of local transcription-factor concentrations at sites of transcription and of transcriptional initiation with high spatiotemporal resolution, such as afforded by lattice light-sheet microscopy (Mir et al., 2018), could provide further information about chromatin accessibility dynamics. Finally, different theoretical models may make distinct predictions about the effect of modulating the number, placement, and affinity of Bicoid and Zelda sites (and even of nucleosomes) in the *hunchback* enhancer. These models could be tested with future experiments that implement these modulations in reporter constructs.

In sum, here we engaged in a theory-experiment dialogue to respond to the theoretical challenges of proposing a passive MWC mechanism for chromatin accessibility in eukaryotes (Mirny, 2010; Narula and Igoshin, 2010; Marzen et al., 2013); we also questioned the suitability of thermodynamic models in the context of development (Estrada et al., 2016). At least regarding the activation of *hunchback*, and likely similar developmental genes such as *snail* and *sog* (Dufourt et al., 2018; Yamada et al., 2019), we speculate that Bicoid and Zelda actively drive chromatin accessibility, possibly through histone acetylation. Once chromatin becomes accessible, thermodynamic models can predict *hunchback* transcription without the need to invoke energy expenditure and non-equilibrium models. Regardless of whether we have identified the only possible model of chromatin accessibility and regulation, we have demonstrated that this dialogue between theoretical models and the experimental testing of their predictions at high spatiotemporal resolution is a powerful tool for biological discovery. The new insights afforded by this dialogue will undoubtedly refine theoretical descriptions of transcriptional regulation as a further step toward a predictive understanding of cellular decision-making in development.

## Acknowledgments

We are grateful to Jack Bateman, Jacques Bothma, Mike Eisen, Jeremy Gunawardena, Jane Kondev, Oleg Igoshin, Rob Phillips, Christine Rushlow and Peter Whitney for their guidance and comments on our manuscript. We thank Kenneth Irvine and Yuanwang Pan for providing the *his-irfp* fly line. This work was supported by the Burroughs Wellcome Fund Career Award at the Scientific Interface, the Sloan Research Foundation, the Human Frontiers Science Program, the Searle Scholars Program, the Shurl and Kay Curci Foundation, the Hellman Foundation, the NIH Directors New Innovator Award (DP2 OD024541-01), and an NSF CAREER Award (1652236) (HGG), an NSF GRFP (DGE 1752814) and UC Berkeley Chancellor’s Fellowship (EE), and the DoD NDSEG graduate fellowship (JL).

## 4. Materials and Methods

### 4.1. Predicting Zelda binding sites

Zelda binding sites in the *hunchback* promoter were identified as heptamers scoring 3 or higher using a Zelda alignment matrix (Harrison et al., 2011) and the Advanced PASTER entry form online (http://stormo.wustl. edu/consensus/cgi-bin/Server/Interface/patser.cgi) (Hertz et al., 1990; Hertz and Stormo, 1999). PATSER was run with setting “Seq. Alphabet and Normalization” as “a:t 3 g:c 2” to provide the approximate background frequencies as annotated in the Berkeley Drosophila Genome Project (BDGP)/Celera Release 1. Reverse complementary sequences were also scored.

### 4.2. Fly Strains

Bicoid nuclear concentration was imaged in embryos from line *yw; his2avmrfp1;bicoidE1,egfp-bicoid* (Gregor et al., 2007b). Similarly, Zelda nuclear concentration was determined by imaging embryos from line *sfgfp-zelda;+;his-irfp*. The *sfgfp-zelda* transgene was obtained from Hamm et al. (2017) and the *hisiRFP* transgene is courtesy of Kenneth Irvine and Yuanwang Pan.

Transcription from the *hunchback* promoter was measured by imaging embryos resulting from crossing female virgins *yw;HistoneRFP;MCP-NoNLS(2)* with male *yw;P2P-MS2-LacZ/cyo;+* (Garcia et al., 2013).

In order to image transcription in embryos lacking maternally deposited Zelda protein, we crossed mother flies whose germline was *w, his2av-mrfp1,zelda(294),FRT19A;+;MCP-egfp(4F)/+* obtained through germline clones (see below) with fathers carrying the *yw;P2P-MS2-LacZ/cyo;+* reporter. The zelda(294) transgene is courtesy of Christine Rushlow (Liang et al., 2008). The *MCP-egfp(4F)* transgene expresses approximately double the amount of MCP than the *MCP-egfp(2)* (Garcia et al., 2013), ensuring similar levels of MCP in the embryo in all experiments.

Imaging Bicoid nuclear concentration in embryos lacking maternally deposited Zelda protein was accomplished by replacing the *MCP-egfp(4F)* transgene described in the previous paragraph with the *bicoidE1,egfp-bicoid* transgene used for imaging nuclear Bicoid in a wildtype background. We crossed mother flies whose germline was *w, his2av-mrfp1,zelda(294),FRT19A;+;bicoidE1,egfp-bicoid/+* obtained through germline clones (see below) with *yw* fathers.

### 4.3. Zelda germline clones

In order to generate mother flies containing a germline homozygous null for *zelda*, we first crossed virgin females of *w,his2av-mrfp1,zelda(294),FRT19A/FM7,y,B;+;MCP-egfp(4F)/TM3,ser* (or *w, his2av-mrfp1,zelda(294),FRT19A;+;bicoidE1,egfp-bicoid/+* to image nuclear Bicoid) with males of *ovoD,hs-FLP,FRT19A;+;+* (Liang et al., 2008). The resulting heterozygotic offspring were heat-shocked in order to create maternal germline clones as described in Liang et al. (2008). The resulting female virgins were crossed with male *yw;P2P-MS2-LacZ/cyo;+* (Garcia et al., 2013) to image transcription or male *yw* to image nuclear Bicoid concentration.

Male offspring are null for zygotic *zelda*. Female offspring are heterozygotic for functional *zelda*, but zygotic *zelda* is not transcribed until nuclear cycle 14 (Liang et al., 2008), which occurs after the analysis in this work. All embryos lacking maternally deposited Zelda showed aberrant morphology in nuclear size and shape (data not shown), as previously reported (Liang et al., 2008; Staudt et al., 2006).

### 4.4. Sample preparation and data collection

Sample preparation followed procedures described in Bothma et al. (2014), Garcia and Gregor (2018), and Lammers et al. (2019).

Embryos were collected and mounted in halocarbon oil 27 between a semipermeable membrane (Lumox film, Starstedt, Germany) and a coverslip. Data collection was performed using a Leica SP8 scanning confocal microscope (Leica Microsystems, Biberach, Germany). Imaging settings for the MS2 experiments were the same as in Lammers et al. (2019), except the Hybrid Detector (HyD) for the His-RFP signal used a spectral window of 556-715 nm. The settings for the Bicoid-GFP measurements were the same, except for the following. The power setting for the 488 nm line was 10 *µ*W. The confocal stack was only 10 slices in this case, rather than 21, resulting in a spacing of 1.11 *µ*m between planes. The images were acquired at a time resolution of 30 s, using an image resolution of 512 x 128 pixels.

The settings for the Zelda-sfGFP measurements were the same as the Bicoid-GFP measurements, except different laser lines were used for the different fluorophores. The sf-GFP excitation line was set at 485 nm, using a power setting of 10 *µ*W. The His-iRFP excitation line was set at 670 nm. The HyD for the His-iRFP signal was set at a 680-800 nm spectral window. All specimens were imaged over the duration of nuclear cycle 13.

### 4.5. Image analysis

Images were analyzed using custom-written software following the protocol in Garcia et al. (2013). Briefly, this procedure involved segmenting individual nuclei using the histone signal as a nulear mask, segmenting each transcription spot based on its fluorescence, and calculating the intensity of each MCP-GFP transcriptional spot inside a nucleus as a function of time.

Additionally, the nuclear protein fluorescences of the Bicoid-GFP and Zelda-sfGFP fly lines were calculated as follows. Using the histone-labeled nuclear mask for each individual nucleus, the fluorescence signal within the mask was extracted in xyz, as well as through time. For each timepoint, the xy signal was averaged to give an average nuclear fluorescence as a function of z and time. This signal was then maximum projected in z, resulting in an average nuclear concentration as a function of time, per single nucleus. These single nucleus concentrations were then averaged over anterior-posterior position to create the protein concentrations reported in the main text.

### 4.6. Data Analysis

All fits in the main text were performed by minimizing the least-squares error between the data and the model predictions. Unless stated otherwise, error bars reflect standard error of the mean over multiple embryo measurements. See Section S2.1 for more details on how this was carried out for model predictions.

## Supplementary Information

### S1. Equilibrium Models of Transcription

#### S1.1. An overview of equilibrium thermodynamics models of transcription

In this section we give a brief overview of the theoretical concepts behind equilibrium thermodynamics models of transcription. For a more detailed overview, we refer the reader to Bintu et al. (2005b) and Bintu et al. (2005a). These models invoke statistical mechanics in order to to calculate bulk properties of a system by enumerating the probability of each possible microstate of the system. The probability of a given microstate is proportional to its Boltzmann weight e^−βε^, where ε is the energy of the microstate and β = (k_B_T)^−1^ with k_B_ being the Boltzmann constant and T the absolute temperature of the system (Garcia et al., 2007).

Specific examples of these microstates in the context of simple activation are featured in Fig. S1. As reviewed in Garcia et al. (2007), the Boltzmann weight of each of these microstates can also be written in a thermodynamic language that accounts for the concentration of the molecular species, their dissociation constant to DNA, and a cooperativity term ω that accounts for the protein-protein interactions between the activator and RNAP. To calculate the probability of finding RNAP bound to the promoter p_bound_, we divide the sum of the weights of the RNAP-bound states by the sum of all possible states

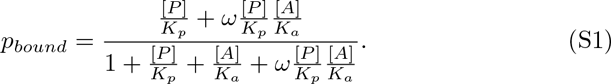

Here, [P] and [A] are the concentrations of RNAP and activator, respectively. K_p_ and K_a_ are their corresponding dissociation constants, and ω indicates an interaction between activator and RNAP: ω > 1 corresponds to cooperativity, whereas 0 < ω < 1 corresponds to anti-cooperativity.

Using p_bound_, we write the subsequent rate of mRNA production by assuming the occupancy hypothesis, which states that

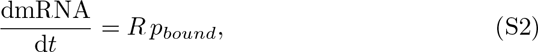

where R is an underlying rate of transcriptional initiation (usually interpreted as the rate of loading RNAP from the promoter-bound state). In the case of simple activation illustrated in Fig. S1, the overall transcriptional initiation rate is then given by

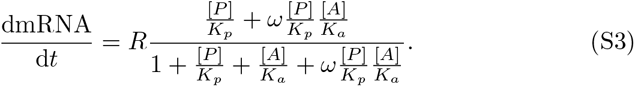

**Figure S1:**
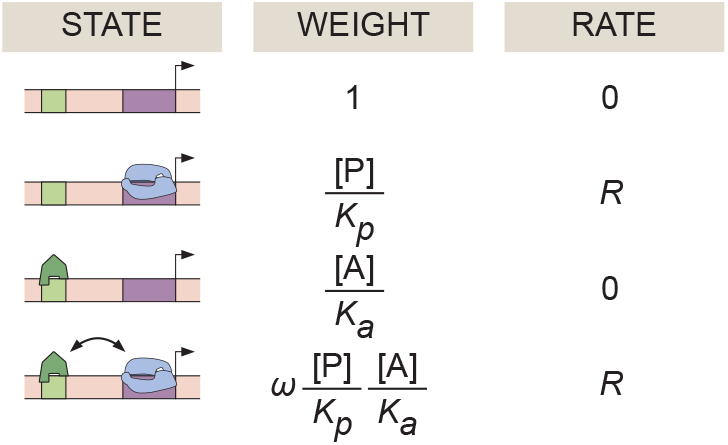
Equilibrium thermodynamic model of simple activation. A promoter region with one binding site for an activator molecule has four possible microstates, each with its corresponding statistical weight and rate of RNAP loading.

From Eq. S1, one can derive the Hill equation that is frequently used to model biophysical binding. In the limit of high cooperativity, 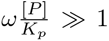 and 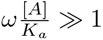 such that

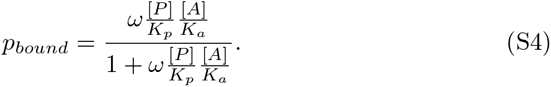

If we then define a new binding constant 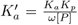, we get the familiar Hill equation of order 1 with a binding constant *K*^2^_a_

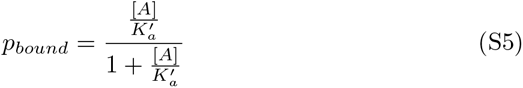

In general, any Hill equation of order n can be derived from a more fundamental equilibrium thermodynamic model of simple activation possessing n activator binding sites in the appropriate limits of high cooperativity. Thus, any time a Hill equation is invoked, equilibrium thermodynamics is implicitly used, bringing with it all of the underlying assumptions described in Section S6.4. This highlights the importance of rigorously grounding the assumptions made in any model of transcription, to better discriminate between the effects of equilibrium and non-equilibrium processes.

#### S1.2. Thermodynamic MWC model

In the thermodynamic MWC model, we consider a system with six Bicoid binding sites and ten Zelda binding sites. In addition, we allow for RNAP binding to the promoter.

In our model, the DNA can be in either an accessible or an inaccessible state. The difference in free energy between the two states is given by −Δε_chrom_, where Δε_chrom_ is defined as

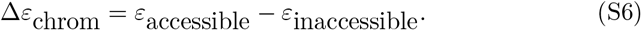

Here, ε_accessible_ and ε_accessible_ are the energies of the accessible and inaccessible states, respectively. A positive Δε_chrom_ signifies that the inaccessible state is at a lower energy level, and therefore more probable, than the accessible state. We assume that all binding sites for a given molecular species have the same binding affinity, and that all accessible states exist at the same energy level compared to the inaccessible state. Thus, the total number of states is determined by the combinations of occupancy states of the three types of binding sites as well as the presence of the inaccessible, unbound state. We choose to not allow any transcription factor or RNAP binding when the DNA is inaccessible.

In this equilibrium model, the statistical weight of each accessible microstate is given by the thermodynamic dissociation constants K_b_, K_z_, and K_p_ of Bicoid, Zelda, and RNAP respectively. The statistical weight for the inaccessible state is e^Δε^chrom. We allow for a protein-protein interaction term ω_b_ between nearest-neighbor Bicoid molecules, as well as a pairwise cooperativity ω_bp_ between Bicoid and RNAP. However, we posit that Zelda does not interact directly with either Bicoid or RNAP. For notational convenience, we express the statistical weights in terms of the non-dimensionalized concentrations of Bicoid, Zelda, and RNAP, given by b, z and p, respectively, such that, for example, 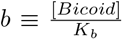. Fig. S2 shows the states and statistical weights for this thermodynamic MWC model, with all the associated parameters.

**Figure S2:**
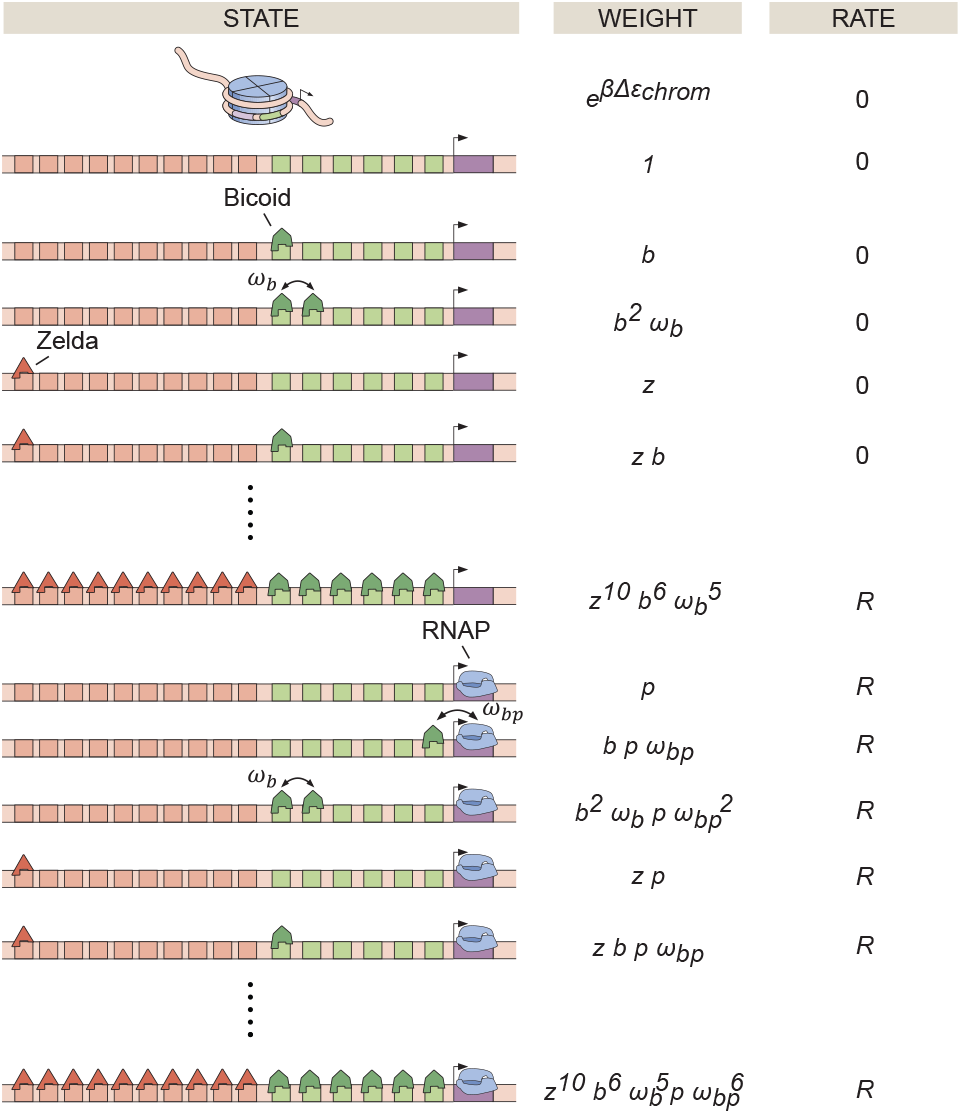
States, weights, and rate of RNAP loading diagram for the thermodynamic MWC model, containing six Bicoid binding sites, ten Zelda binding sites, and a promoter.

Incorporating all the microstates, we can calculate a statistical mechanical partition function, the sum of all possible weights, which is given by

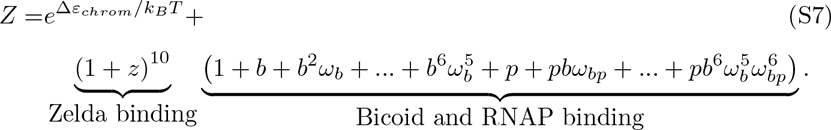

Using the binomial theorem

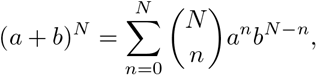

Eq. S7 can be expressed more compactly as

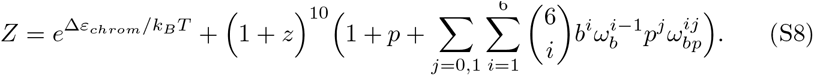

From this partition function, we can calculate p_bound_, the probability of being in an RNAP-bound state. This term is given by the sum of the statistical weights of the RNAP-bound states divided by the partition function

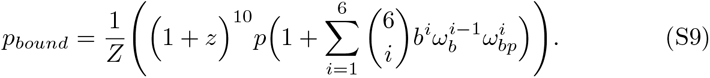

In this model, we once again assume that the transcription associated with each microstate is zero unless RNAP is bound, in which case the associated rate is R. Then, the overall transcriptional initiation rate is given by the product of p_bound_ and R

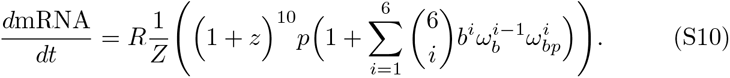

Note that since the MS2 technology only measures nascent transcripts, we can ignore the effects of mRNA degradation and focus on transcriptional initiation.

#### S1.3. Constraining model parameters

The transcription rate R of the RNAP-bound states can be experimentally constrained by making use of the fact that the hunchback minimal reporter used in this work produces a step-like pattern of transcription across the length of the fly embryo (Fig. 4C). Since in the anterior end of the embryo, the observed transcription appears to level out to a maximum value, we assume that Bicoid binding is saturated in this anterior end of the embryo such that

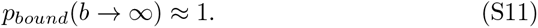

In this limit, Eq. S10 can be written as

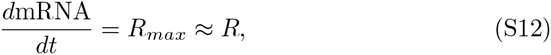

where R_max_ is the maximum possible transcription rate. Importantly, R_max_ is an experimentally observed quantity rather than a free parameter. As a result, the model parameter R is determined by experimentally measurable quantity Rmax.

The value of p can also be constrained by measuring the transcription rate in the embryo’s posterior, where we assume Bicoid concentration to be negligible. Here, the observed transcription bottoms out to a minimum level R_min_ (Fig. 4C), which we can connect with the model’s theoretical minimum rate. Specifically, in this limit, b approaches zero in Eq. S10 such that all Bicoid-dependent terms drop out, resulting in

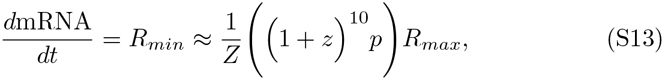

where we have replaced R with R_max_ as described above. Next, we can express p in terms of the other parameters such that

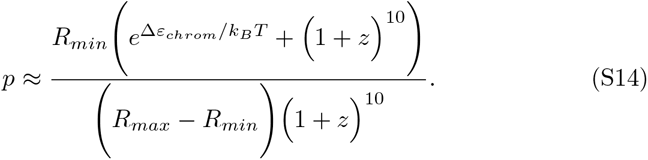

Thus, p is no longer a free parameter, but is instead constrained by the experimentally observed maximum and minimum rates of transcription R_max_ and R_min_, as well as our choices of K_z_ and Δε_chrom_. In our analysis, R_max_ and R_min_ are calculated by taking the mean RNAP loading rate across all embryos from the anterior and posterior of the embryo respectively, extrapolated using the trapezoidal fitting scheme described in Section S2.3.

Finally, we expand this thermodynamic MWC model to also account for suppression of transcription in the beginning of the nuclear cycle via mechanisms such as mitotic repression (Section S3.1). To make this possible, we include a trigger time term t_MitRep_, before which we posit that no readout of Bicoid or Zelda by hunchback is possible and the rate of RNAP loading is fixed at 0. For times t > t_MitRep_, the system behaves according to Eq. S10. Thus, given the constraints stemming from direct measurements of R_max_ and R_min_, the model has six free parameters: Δε_chrom_, ω_b_, ω_bp_, K_b_, K_z_, and t_MitRep_. The final calculated transcription rate is then integrated in time to produce a predicted MS2 fluorescence as a function of time (Section S2.2).

For subsequent parameter exploration of this model (Section S5.1), constraints were placed on the parameters to ensure sensible results. Each parameter was constrained to be strictly positive such that:

- Δε_chrom_ > 0
- K_b_ > 0
- K_z_ > 0
- ω_b_ > 0
- ω_bp_ > 0
- tMitRep > 0.

### S2. Input-Output measurements, predictions, and characterization

#### S2.1. Input measurement methodology

Input transcription-factor measurements were carried out separately in individual embryos containing a eGFP-Bicoid transgene in a bicoid null mutant background (Gregor et al., 2007b) or a Zelda-sfGFP CRISPR-mediated homologous recombination at the endogenous zelda locus (Hamm et al., 2017). Over the course of nuclear cycle 13, the fluorescence inside each nucleus was extracted (details given in Section 4.5), resulting in a measurement of the nuclear concentration of each transcription factor over time. Six eGFP-Bicoid and three Zelda-sfGFP embryos were imaged.

Representative fluorescence traces of eGFP-Bicoid for a single embryo indicate that the magnitude of eGFP-Bicoid fluorescence decreases for nuclei located toward the posterior of the embryo (Fig. S3A). Further, the nuclear fluorescence of eGFP-Bicoid at 8 min into nuclear cycle 13 (Fig. S3B) exhibited the known exponential decay of Bicoid, with a mean decay length of 23.5% ± 0.6% of the total embryo length, consistent with but slightly different than previous measurements that suggested a mean decay length of 19.1%±0.8% (Liu et al., 2013).

This discrepancy could stem, for example, from minor differences in acquisition from the laser-scanning two-photon microscope used in Liu et al. (2013) versus the laser-scanning confocal microscope used here, such as differences in axial resolution (due both to different choices of objectives and the inherent differences in axial resolution of one-photon and two-photon fluorescence excitation processes). Nevertheless, the difference was minute enough that we felt confident in our eGFP-Bicoid measurements.

Intra-embryo variability in eGFP-Bicoid nuclear fluorescence, defined by the standard deviation across nuclei within a single embryo divided by the mean, was in the range of 10-30%, as was the inter-embryo variability, defined by the standard deviation of the mean amongst nuclei, across different embryos (Fig. S3C, blue and black, respectively). Six separate eGFP-Bicoid embryos were measured.

Similarly, representative fluorescence time traces of Zelda-sfGFP for a single embryo are shown in Fig. S3D. Unlike the eGFP-Bicoid profile, the Zelda-sfGFP nuclear fluorescence was approximately uniform across embryo position (Fig. S3E), consistent with previous fixed-tissue measurements (Staudt et al., 2006; Liang et al., 2008). Intra-embryo variability in Zelda-sfGFP nuclear fluorescence was very low (less than 10%), whereas inter-embryo variability was relatively higher, up to 20% (Fig. S3F, red and black, respectively). Three separate Zelda-sfGFP embryos were measured.

Due to the consistency of Zelda-sfGFP nuclear fluorescence, we assumed the Zelda profile to be spatially uniform in our analysis, and thus created a mean Zelda-sfGFP measurement for each individual embryo by averaging all mean nuclear fluorescence traces in space across the anterior-posterior axis of the embryo (Fig. S3D, inset). This mean measurement was used as an input in the theoretical models. However, we still retained inter-embryo variability in Zelda, as described below.

To combine multiple embryo datasets as inputs to the models explored throughout this work, the fluorescence traces corresponding to each dataset were aligned at the start of nuclear cycle 13, defined as the start of anaphase. Because each embryo may have possessed slightly different nuclear cycle lengths and/or experimental sampling rates (due to the manual realignment of the z-stack to keep nuclei in focus), the individual datasets were not combined in order to create average Bicoid and Zelda profiles across embryos. Instead, a simulation and model prediction were performed for each combination of measured input Bicoid and Zelda datasets, essentially an *in silico* experiment covering a portion of the full embryo length. In all, outputs at each embryo position were predicted in at least three separate simulations. Subsequent analyses used the mean and standard error of the mean of these amalgamated simulations. With six GFP-Bicoid datasets and three Zelda-GFP datasets, there were 18 unique combinations of input embryo datasets; for a single set of parameters used in a particular model, each derived metric (e.g. *t_on_*) was calculated using predicted outputs from each of the 18 possible input combinations. This procedure provided full embryo coverage and resulted in a distribution of the derived metric for that particular set of parameters. From this distribution, the mean and standard error of the mean were calculated, leading to the error bars in plots such as Fig. S6.

#### S2.2. MS2 fluorescence simulation protocol

To calculate a predicted MS2 fluorescence trace from measured Bicoid and Zelda inputs for a given theoretical model, we utilized a simple model of transcription initiation, elongation, and termination. First, the dynamic transcription-factor concentrations were used as inputs to each of the theoretical models outlined throughout the paper. These models generated a rate of RNAP loading as a function of time and space across the embryo over the course of nuclear cycle 13.

For each position along the anterior-posterior axis, the predicted rate of RNAP loading was integrated over time to generate a predicted MS2 fluorescence trace. Given the known reporter construct length L of 5.2 kb (Garcia et al., 2013), we assume that RNAP molecules are loaded onto the start of the gene at a rate R(t) predicted by the particular model under consideration (Fig. S4; see Sections S1.2, S6.1, S7.1, and S8.1 for model details). Each RNAP molecule traverses the gene at a constant velocity v of 1.54 kb/min, as measured experimentally by Garcia et al. (2013). With these numbers, we calculate an elongation time

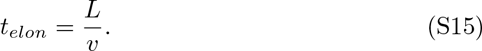

**Figure S3:**
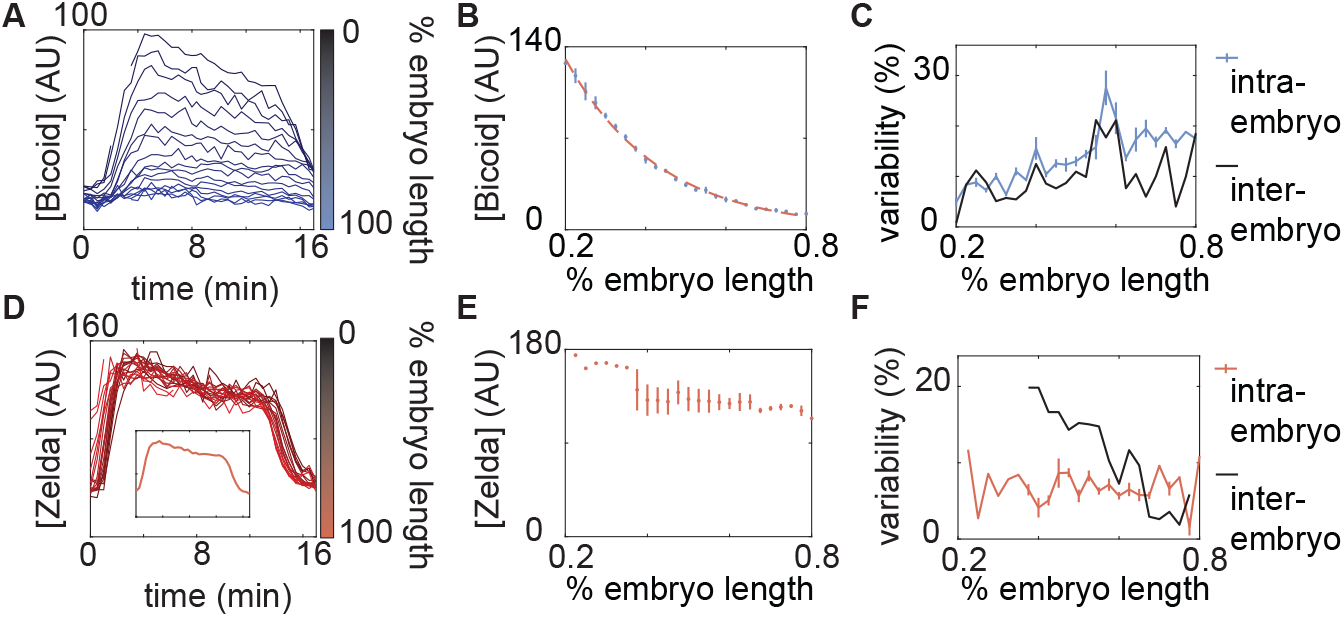
Measurements of input transcription-factor concentration dynamics. (A) Nuclear eGFP-Bicoid concentration as a function of time into nuclear cycle 13 across various positions along the anterior-posterior axis of a single embryo. (B) eGFP-Bicoid concentration at 8 min into nuclear cycle 13 as a function of position along the embryo averaged over all measured embryos (n=6). The fit of the concentration profile to an exponential function results in a decay length of 23% *±* 0.6% embryo length. (C) Intra- and inter-embryo variability in eGFP-Bicoid nuclear fluorescence along the anterior-posterior axis. (D) Zelda-sfGFP concentration as a function of time into nuclear cycle 13 across various anterior-posterior positions of a single embryo. (D, inset) Zelda-sfGFP concentration averaged over the data shown in D. (E) Zelda-sfGFP concentration at 8 min into nuclear cycle 13 as a function of position along the anterior-posterior axis of the embryo averaged over all measured embryos (n=3). Note that anterior of 40% and posterior of 77.5% only a single embryo was measured; no error bars were calculated. (F) Intra- and inter-embryo variability in Zelda-sfGFP nuclear fluorescence along the anterior-posterior axis. (B,E, error bars represent standard error of the mean nuclear fluorescence, measured across embryos; C,F, error bars represent standard error of the mean intra-embryo variability, measured across embryos.)

Finally, we assume that upon reaching the end of the reporter gene, the RNAP molecules terminate and disappear instantly such that they no longer contribute to spot fluorescence.

The MS2 fluorescence signal reports on the number of RNAP molecules actively occupying the gene at any given time and, under the assumptions outlined above, is given by the integral

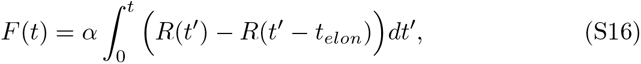

where F (t) is the predicted fluorescence value, R(t) is the RNAP loading rate predicted by each specific model, R(t − t_elon_) is the time-shifted loading rate that accounts for RNAP molecules finishing transcription at the end of the gene, and α is an arbitrary scaling factor to convert from absolute numbers of RNAP molecules to arbitrary fluorescence units. The predicted value F (t) was scaled by α to match the experimental data.

The final predicted MS2 signal was modified in a few additional ways. First, any RNAP molecule that had not yet reached the position of the MS2 stem loops had its fluorescence value set to zero (Fig. S4, i), since only RNAP molecules downstream of the MS2 stem loop sequence exhibit a fluorescent signal. Second, RNAP molecules that were only partially done elongating the MS2 stem loops contributed a partial fluorescence intensity, given by the ratio of the distance traversed through the stem loops to the total length of the stem loops

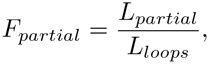

where F_partial_ is the partial fluorescence contributed by an RNAP molecule within the stem loop sequence region, L_partial_ is the distance within the stem loop sequence traversed, and L_loops_ is the length of the stem loop sequence (Fig. S4, ii). For this reporter construct, the length of the stem loops was approximately L_loops_ = 1.28 kb. RNAP molecules that had finished transcribing the MS2 stem loops contributed the full amount of fluorescence (Fig. S4, iii). Finally, to make this simulation compatible with the trapezoidal fitting scheme in Section S2.3, we included a falling signal at the end of the nuclear cycle, achieved by setting R(t) = 0 after 17 min into the nuclear cycle and thus preventing new transcription initiation events.

**Figure S4:**
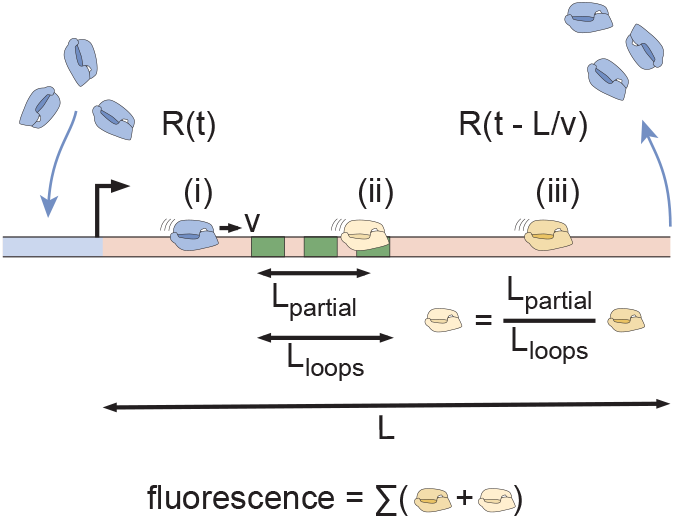
MS2 fluorescence calculation protocol. RNAP molecules load onto the reporter gene at a time-dependent rate *R*(*t*), after which they elongate at a constant velocity *v*. Upon reaching the end of the gene after a length *L* has been transcribed, they are assumed to terminate and disappear instantly, given by the time-shifted rate 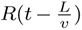. The time-dependent MS2 fluorescence is calculated by summing the contributions of RNAP molecules that are located before, within, or after the MS2 stem loop sequence (i, ii, and iii, respectively).

Given the predicted MS2 fluorescence trace, the rate of RNAP loading and t_on_ were extracted with the fitting procedure used on the experimental data (Section S2.3).

#### S2.3. Extracting initial RNAP loading rate and transcriptional onset time

To extract the initial rate of RNAP loading and the transcriptional onset time t_on_ used in the data analysis, we fit both the experimental and calculated MS2 signals to a constant loading rate model, the trapezoidal model (Garcia et al., 2013).

The trapezoidal model provides a heuristic fit of the main features of the MS2 signal by assuming that the RNAP loading rate is either zero or some constant value r (Fig. S5A). At time t_on_, the loading rate switches from zero to this constant value r, producing a linear rise in the MS2 signal. After the elongation time t_elong_, the loading of new RNAP molecules onto the gene is balanced by the loss of RNAP molecules at the end of the gene, producing a plateau in the MS2 signal. Finally, at the end of the nuclear cycle, transcription ceases at t_off_ and the RNAP loading rate switches back to zero, producing the falling edge of the MS2 signal and completing the trapezoidal shape. Because we only consider the initial dynamics of transcription in the nuclear cycle in this investigation, we do not explore the behavior of t_off_.

Fig. S5B shows the results of fitting the mean MS2 fluorescence from a narrow window within a single embryo to the trapezoidal model (Section 2.3). With this fit, we can extract the initial rate of RNAP loading (given by the initial slope) as well as *t_on_* (given by the intercept of the fit onto the x-axis).

As a consistency check, the *t_on_* values extrapolated from the trapezoidal fit of the data were compared with the experimental time points at which the first MS2 spots were observed for both the wild-type and *zelda^−^* mutant experiments (Fig. S5C). Due to the detection limit of the microscope, this latter method reports on the time at which a few RNAP molecules have already begun transcribing the reporter gene, rather than a “true” transcriptional onset time. Using the first frame of spot detection yields similar trends to the trapezoidal fits, except that the measured first frame times are systematically larger by 3-5 min. Additionally, utilizing the first frame of detection to measure *t_on_* appears to be a noisier method, likely because the actual MS2 spots cannot be observed below a finite signal-detection limit, whereas the extrapolated *t_on_* from the trapezoidal fit corresponds to a “true” onset time below the signal-detection limit.

**Figure S5:**
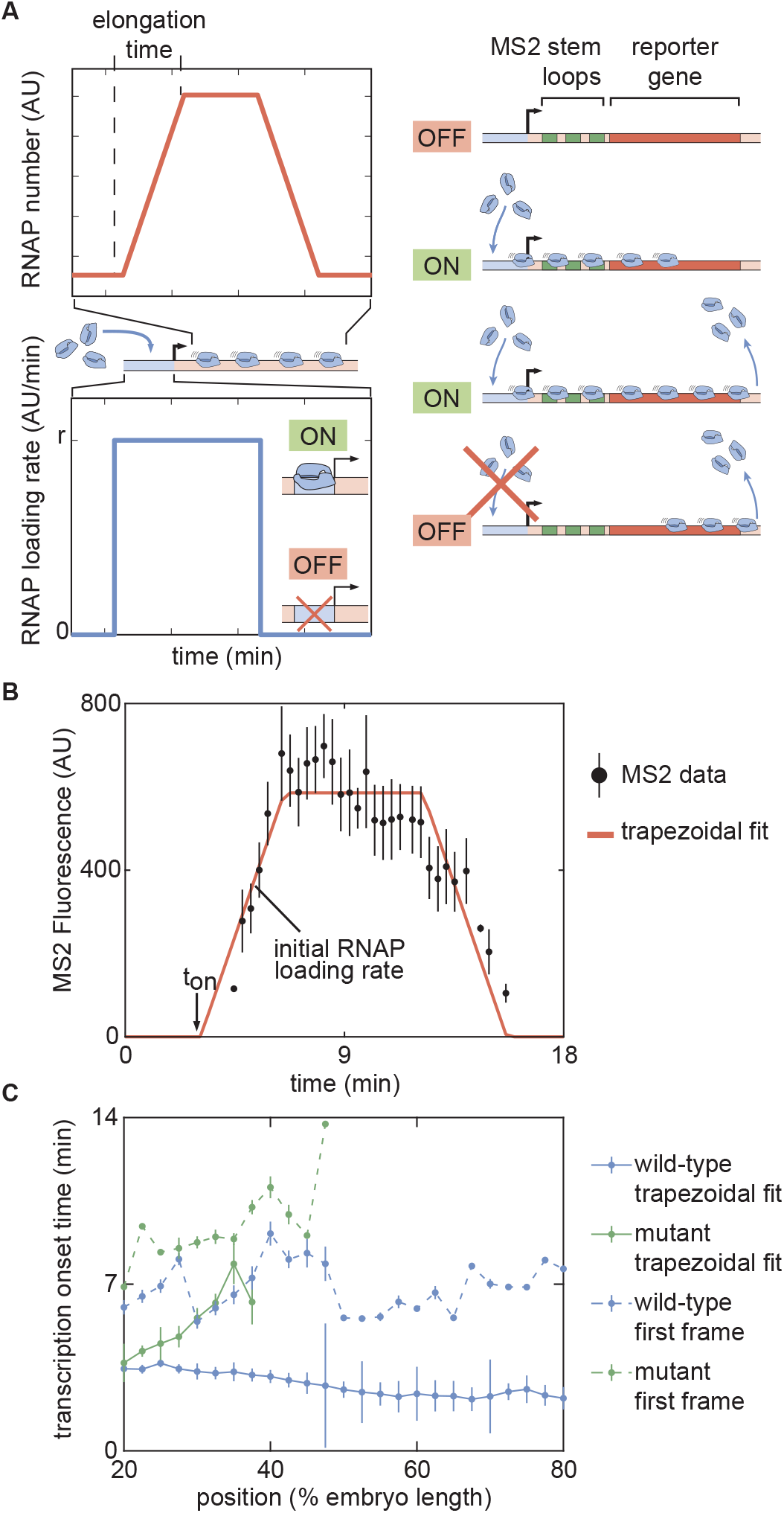
Outline of fitting to the trapezoidal model of transcription. (A) The trapezoidal model of transcription, where transcription begins at an onset time *t_on_* and loads RNAP molecules with a constant rate *r*. (B) Results of fitting the MS2 fluorescence data from a single embryo to the trapezoidal model to extract *t_on_* and the initial rate of RNAP loading. (C) Comparison of inferred *t_on_* values between the trapezoidal model (solid lines) and using the time of first detection of signal in a fluorescence spot (dashed lines) for both wild-type and *zelda^−^* backgrounds. (B, error bars are standard error of the mean averaged over multiple nuclei within the embryo, for data in a wild-type background at 50% along the embryo length; C, error bars are standard error of the mean, averaged across embryos).

### S3. Mitotic Repression

#### S3.1. Mitotic repression is necessary to recapitulate Bicoid- and Zelda-mediated regulation of hunchback using the thermodynamic MWC model

As described in Section 2.3 of the main text, a mitotic repression window was incorporated into the thermodynamic MWC model (Section S1.2) in order to explain the observed transcriptional onset times of *hunchback*. Here, we justify and explain this theoretical modification in greater detail.

Fig. S6A and B depicts the experimentally observed initial rates of RNAP loading and *t_on_* across the length of the embryo (blue points) for the wild-type background. After constraining the maximum and minimum theoretically allowed rates of RNAP loading (Section S1.3), we attempted to simultaneously fit the thermodynamic MWC model to both the rate of RNAP loading and *t_on_*.

The fit results demonstrate that while the thermodynamic MWC model can recapitulate the measured step-like rate of RNAP loading at *hunchback* (Fig. S6A, purple line), it fails to predict the *t_on_* throughout the embryo (Fig. S6B, purple line; see Sections S2.2 and S2.3 for details about experimental and theoretical calculations). This model yields values of *t_on_* that are much smaller than those experimentally observed, a trend that holds throughout the length of the embryo. This disagreement becomes more evident when comparing the output transcriptional activity reported by the measured MS2 fluorescence with the input concentrations of Bicoid and Zelda. Specifically, the Bicoid and Zelda concentration measurements at 45% along the embryo, shown for a single embryo in Fig. S6C, are used in conjunction with the previously mentioned best-fit model parameters to predict the output MS2 signal at the same position. This prediction can then be directly compared with experimental data (Fig. S6D, purple line vs. black points, respectively). Whereas the model predicts that transcription will commence around 1 min after anaphase due to the concurrent increase in the Bicoid and Zelda concentrations, the observed MS2 signal begins to increase around 3 min after anaphase (Fig. S6D). As a result, the predicted transcriptional dynamics in Fig. S6D are systematically shifted in time with respect to the observed data.

**Figure S6:**
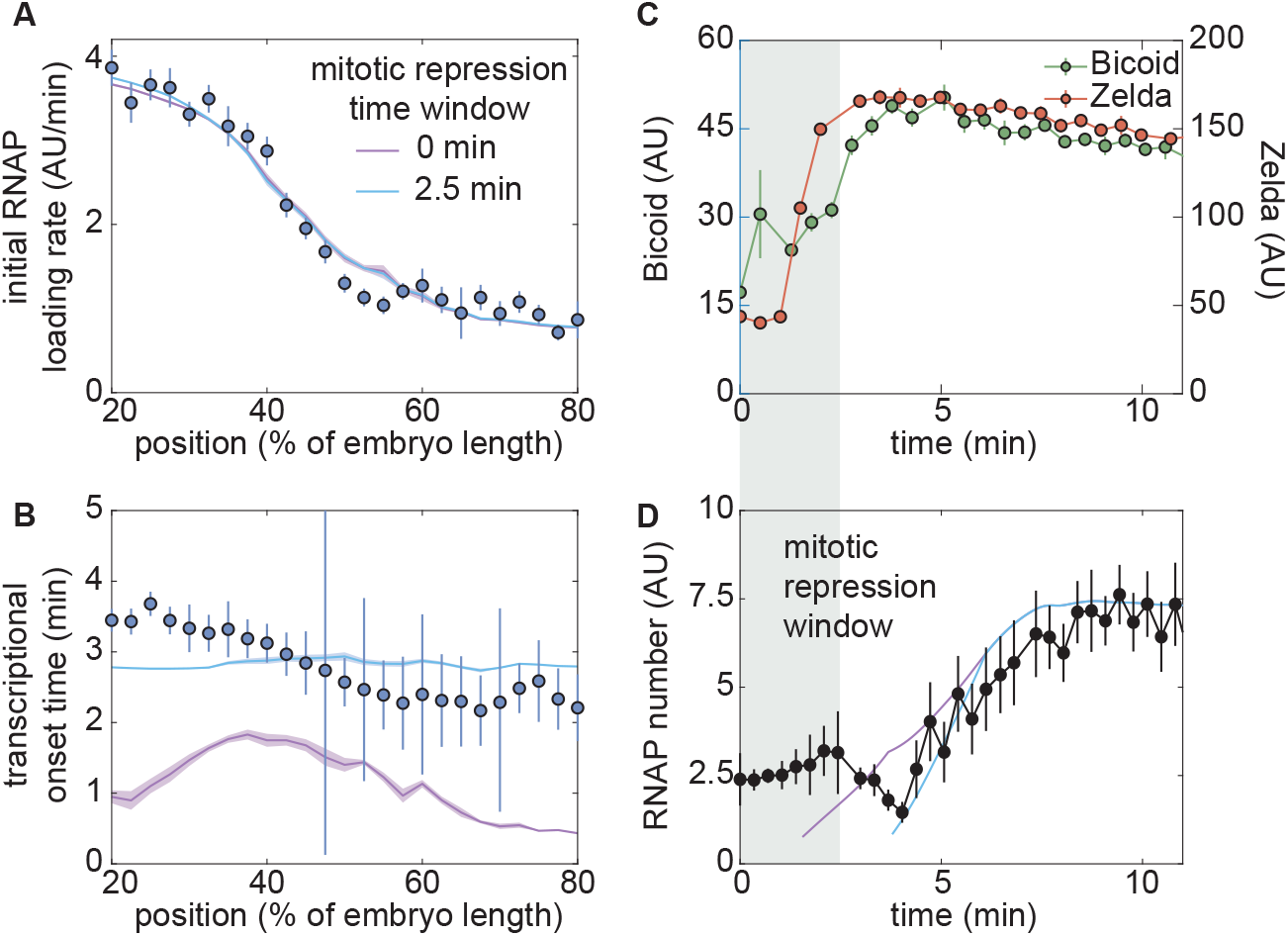
A thermodynamic MWC model including mitotic repression can recapitulate *hunchback* regulation by Bicoid and Zelda. (A) Measured initial rates of RNAP loading and (B) *t_on_* (blue points) across the length of the embryo, compared to fits to the thermodynamic MWC model with and without accounting for mitotic repression (blue and purple curves, respectively). (C) Nuclear concentration dynamics of Bicoid and Zelda with proposed mitotic repression window (gray shading). (D) Predicted MS2 dynamics with no mitotic repression term or a 2.5 min mitotic repression window compared to experimental measurements. (A,B, solid lines indicate mean predictions of the model and shading represents standard error of the mean, while points indicate data and error bars represent the standard error of the mean, across 8 embryos; C, D, data from single embryos at 45% of the embryo length with error bars representing the standard error of the mean across nuclei, errors in model predictions in D were negligible and are obscured by the prediction curve; fitted parameter values for a 2.5 min mitotic repression window were Δ*E_chrom_*= 8.9 *k_B_T*, *K_b_*= 152.3 *AU*, *Kz*= 416.3 *AU*, with different arbitrary fluorescent units for Bicoid and Zelda, *ω_b_* = 6.4, *ω_bp_* = 2.5, for a model assuming six Bicoid binding sites.)

The observed disagreement in *t_on_* suggests that in this model, transcription is prevented from starting at the time dictated solely by the increase of Bicoid and Zelda concentrations. While we speculate that this effect could stem from processes such as RNAP escape from the promoter, DNA replication at the start of the cell cycle, and post-mitotic nucleosome clearance from the promoter, we choose not to commit to a detailed molecular picture and instead ascribe this transcriptional refractory period at the beginning of the nuclear cycle to mitotic repression, the observation that the transcriptional machinery cannot operate during mitosis (Shermoen and O’Farrell, 1991; Gottesfeld and Forbes, 1997; Parsons and Tg, 1997; Garcia et al., 2013). To account for this phenomenon, we revised our thermodynamic MWC model by stating that *hunchback* can only read out the inputs and begin transcription after a specified mitotic repression time window after the previous anaphase (Section S1.3).

Since we expect mitotic repression to operate independently of position along the length of the embryo (Shermoen and O’Farrell, 1991), we assumed that the duration of mitotic repression was uniform throughout the embryo. After incorporating a uniform 2.5 min mitotic repression window into the thermodynamic MWC model (Fig. S6C and D, grey shaded region), the model successfully recapitulates *t_on_* throughout the embryo (Fig. S6B and D, blue curves), while still explaining the observed rates of RNAP loading (Fig. S6A, blue curve). Thus, once mitotic repression is accounted for, the thermodynamic MWC model based on statistical mechanics can quantitatively recapitulate the regulation of *hunch-back* transcription by Bicoid and Zelda.

### S4. *zelda^−^* mutant embryos

#### S4.1. The effect of the zelda^−^ background on the Bicoid concentration profile

Our models rest on the assumption that the Bicoid gradient remains unaltered regardless of whether these measurements are made in the wild-type or *zelda^−^* backgrounds. To confirm this assumption, we measured eGFP-Bicoid concentrations in a *zelda^−^* background. These flies were heterozygous for eGFP-labeled Bicoid and for wild-type Bicoid, resulting in roughly 50% of total Bicoid being labeled with eGFP. As shown in Fig. S7, the resultant eGFP-Bicoid nuclear fluorescence levels at 8 min into nuclear cycle 13 in the *zelda^−^* background (red) were roughly half the magnitude of the equivalent measurements in the wild-type background (blue). After doubling the heterozygote eGFP-Bicoid nuclear fluorescence measurements to rescale them (black), the two eGFP-Bicoid measurement curves were similar, although the *zelda^−^* eGFP-Bicoid values were systematically lower than in the wild-type background. The normalized difference, defined as the absolute value of the difference between the wild-type and *zelda^−^* profiles at each position in the embryo divided by the value of the wild-type profile at the position, averaged across all measured positions, was 15.46% *±* 1.64%. This value is within the range of the inter-embryo variability of eGFP Bicoid in wild-type background embryos (Fig. S3C). Measuring the decay length of the eGFP-Bicoid profile in the *zelda^−^* background also yielded a slightly different result: 20.8% +/- 1.4% of the total embryo length, as opposed to 23.5% +/- 0.6% in the wild-type background (dashed curves).

Nevertheless, these differences would have a negligible effect on our overall conclusions. In the context of our models, an overall rescaling in the magnitude of the Bicoid gradient between the wild-type and *zelda^−^* backgrounds can be compensated by a corresponding rescaling in the dissociation constant of Bicoid, *K_b_*. Because our systematic exploration of theoretical models considers many possible parameter values (Section S5.1), this rescaling has no effect on our conclusion that the equilibrium models are insufficient to explain the *zelda^−^* data. As a result and given that our statistics for the wild-type eGFP-Bicoid data consisted of more embryos than the data for the *zelda^−^* background, we used this wild-type data in our analyses as an input to both the wild-type and *zelda^−^* model calculations.

**Figure S7:**
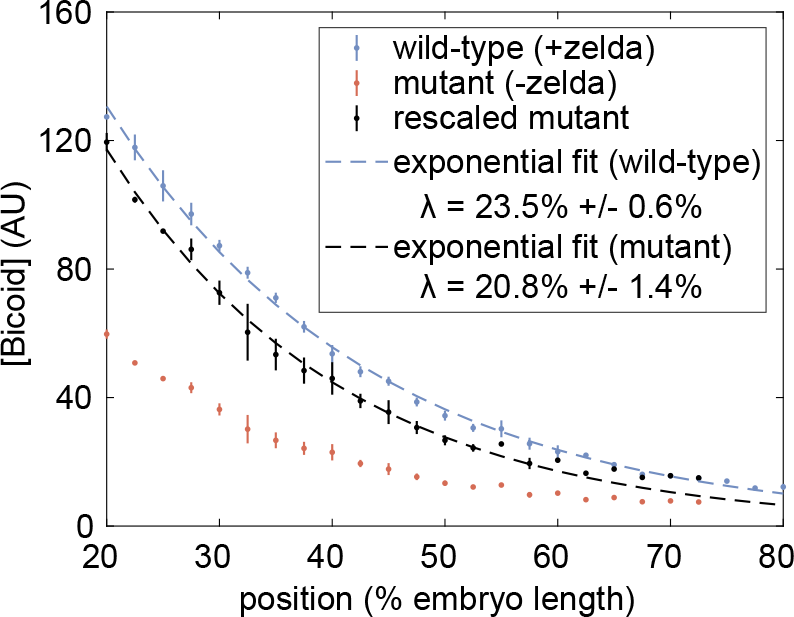
eGFP-Bicoid measurements in wild-type (blue) and *zelda^−^* mutant embryos (red), along with rescaled mutant profiles (black). Fits to an exponentially decaying function yield decay lengths in each background (blue and black dashed curves). A total of n=3 embryos were measured. All error bars are standard error of mean across embryos.

### S5. State-space exploration of theoretical models

#### S5.1. General methodology

To help visualize the limits of our models, we collapsed our observations onto a two-dimensional state space, following a method similar to that described in Estrada et al. (2016). In this space, the x-axis is the average t_on_ delay. This magnitude was computed by integrating the t_on_ across 20% to 37.5% of the embryo length, corresponding to the range in which both wild-type and zelda^−^ experiments exhibited transcription in at least 50% of observed nuclei (Figs. S8A and 5A; Eq. 4). We first subtracted all values of t_on_ by the value at 20% of the embryo length, removing the “baseline” from the calculation of the integral. After computing the integral, the resulting value was then normalized by dividing by the distance along the embryo integrated over (see Section 2.3 for more detail).

The y-axis in our state space exploration plane is given by the position along the embryo at which the rate of RNAP loading reaches its midpoint (Fig. S8B) when fitted to a Hill function of Hill-coefficient six, the number of known Bicoid binding sites on the hunchback P2 enhancer (Driever and Nusslein-Volhard, 1988a; Park et al., 2019). The maximum and minimum values of the RNAP loading rate are fixed to R_max_ and R_min_ (Section S1.3), motivated by the observation that the wild-type RNAP loading rates exhibit maximum and minimum plateaus at either end of the embryo (Fig. S6A). Note that due to the large space of possible model behaviors, many predicted rate profiles did not fit well to this Hill function (e.g. Fig. S8B). Instead, this process should be interpreted as a way to extract, for a particular model realization, the best-fit midpoint position of the observed wild-type RNAP loading rate profile. If a particular model is capable of recapitulating the experimental data, then the best-fit midpoint position under this protocol should agree with that of the data. Hereafter we refer to this quantity as the rate midpoint position.

Combined, the average t_on_ delay and the rate midpoint position provide a simplified description of our data as well as of our theoretical predictions. Each theoretical model inhabits a finite region in this two-dimensional state space, which we can calculate by systematically varying model parameters. Fig. S8A and B show an example of how the average t_on_ delay and the rate midpoint position are calculated using the wild-type background data presented in Fig. 4C and D (points) in the main text.

Due to the large number of parameters of each model explored, the corresponding state-space boundaries were generated by efficient sampling of the underlying high-dimensional parameter space (Fig. S8C). The methodology is similar to the one described in Estrada et al. (2016). Briefly, a set of 100 initial points was generated, each with a randomized set of initial parameters, the specifics of which depended on the model being tested (Fig. S8C, i). The state space was sectioned into 10 horizontal and vertical slices (Fig. S8C, ii). The most extremal points in each slice were found, resulting in two extremal anterior-posterior positions for the mid-point of the rate of RNAP loading and two extremal average t_on_ delay points (Fig. S8C, iii). For each of these points, a new set of five points was generated using random parameters within a small neighborhood of the seed points determined by the extremal points of the previous iteration (Fig. S8C, iv). These new points were plotted; some of these points may be more extreme than the previous set of points. Steps ii-iv were iterated, resulting in a growing boundary over time (Fig. S8C, v). Constraints imposed by the data were used to filter unrealistic results. First, if the simulated t_on_ at the most anterior position (20% of the embryo length) was greater than 5.5 min or less than 1.5 min, the point was filtered out. This removal was justified experimentally, since none of the observed transcriptional onset values in this position in either experiment lay outside of these bounds (Fig. 4D and G). Second, if the simulated rate midpoint position was smaller than 16% or larger than 100% of the embryo length, then this point was also excluded. This exclusion was also justified experimentally, since the rate midpoints values of both experiments were within this interval (Fig. 4C and F). Points that fulfilled these constraints were retained for the next iteration of the algorithm. This process was repeated until the resulting space of points no longer grew appreciably, resulting in an estimate of the size and shape of the state space for each of the models presented in Sections S1.2, S6.1, S7.1, and S8.1.

To determine whether the algorithm had indeed converged, the total area of each model’s region in parameter space was tracked with each iteration number. If the algorithm worked well, then this area would approach some saturation value. Fig. S8D shows the area of the state space corresponding to each model (normalized by the final area at the final iteration number) as a function of the iteration number. Each model converged to a finite value, indicating that the parameter space occupied by the models had been thoroughly explored.

### S5.2. State space exploration with the thermodynamic MWC model

Fig. S9A shows the resulting two-dimensional state space for the thermodynamic MWC model (green shading), as well as all of the theoretical models considered here. We plotted the wild-type and *zelda^−^* data on the same state space. While the wild-type data are represented as a small ellipse of uncertainty, the *zelda^−^* data appear as a large region because we cannot accurately determine the midpoint of the rate of RNAP loading due to the lack of transcriptional activity past about 40% of the embryo length. This shaded region represents a conservative estimate of the possible location of the *zelda^−^* data in this state space. Any successful model must occupy a region that overlaps both the wild-type data point and the *zelda^−^* data region.

The state space corresponding to the thermodynamic MWC model fails to overlap with the *zelda^−^* data. To make the picture more intuitive, this two-dimensional state space was projected onto the x-dimension, the average *t_on_* delay. To do this projection, we reasoned that since both the wild-type and *zelda^−^* data only occupied midpoint values between 40% and 100%, we would only retain points in that range. The resulting subset of state space was collapsed onto the x-axis, resulting in the bars shown in Fig. S9B. Even in this one-dimensional representation, the failure of the thermodynamic MWC model (Fig. S9B, green bar) is evident. This one-dimensional projection is the one presented in the main text.

### S6. Failures and assumptions of thermodynamic models of transcription

#### S6.1. Generalized thermodynamic model

The generalized thermodynamic model is an extension of the thermodynamic MWC model presented in Section S1.2. For extra generality, we assume the presence of twelve Bicoid binding sites and one RNAP binding site, but do not include the action of Zelda since the objective was to attempt to recapitulate the zelda^−^ mutant experimental data. We still allow for an inaccessible DNA state.

In this generalized model, the weight of each microstate can be arbitrary, rather than determined by underlying biophysical parameters. Since p_bound_ only depends on whether RNAP is bound, there is no need to distinguish between different microstates that have the same number of Bicoid molecules bound: the arbitrary coefficients allow separate microstates to effectively be combined together into the same weight. Thus, each microstate corresponds only to the overall number of bound molecules, regardless of binding site ordering. With twelve Bicoid sites, in addition to the inaccessible state, there are 27 total microstates and 26 free parameters describing the weights of each state (with the accessible, unbound microstate normalized to unity). Like with the thermodynamic MWC model, we assume that transcription only occurs when RNAP is bound, with the same constrained maximum rate of RNAP loading R_max_. However, since the weights of each microstate are arbitrary, we no longer have a variable p that can be constrained by R_min_ like in Eq. S14.

**Figure S8:**
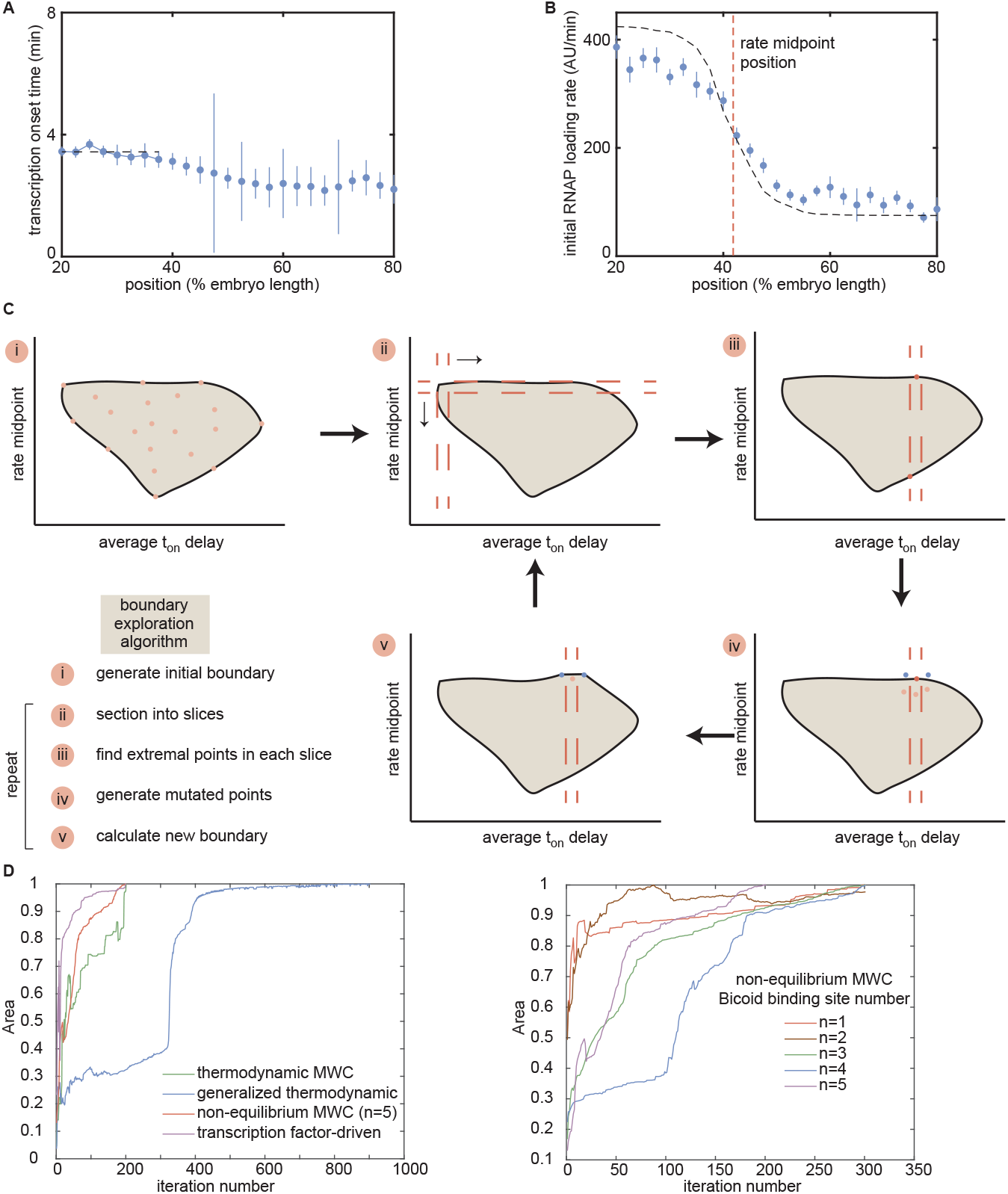
Description of state-space metrics and boundary-exploration algorithm. (A) Representative average *t_on_*delay (black dashed line) for the wild-type background data in Fig. 4D, calculated by integrating the area under the curve from positions 20% to 37.5% along the embryo length and normalizing by dividing by the length over which the integral was taken. (B) Rate midpoint position for the wild-type background data in Fig. 4C, calculated by fitting a Hill equation of order 6 to the initial rates of RNAP loading. (C) Overview of the boundary-exploration algorithm. (i) A set of 100 points with random input parameters generates an initial state space of the investigated model. (ii) The space is sectioned into 10 horizontal and 10 vertical slices. (iii) The extremal points of each slice are found. (iv) For each extremal point, five new points are generated with input parameters in a small neighborhood around the parameters of this extremal point. (v) The new space is plotted with these new points, and steps (ii) - (iv) are repeated. (D) Normalized area of each investigated model’s region in parameter space as a function of algorithm iteration number. All areas approach a steady value, indicating convergence.

**Figure S9:**
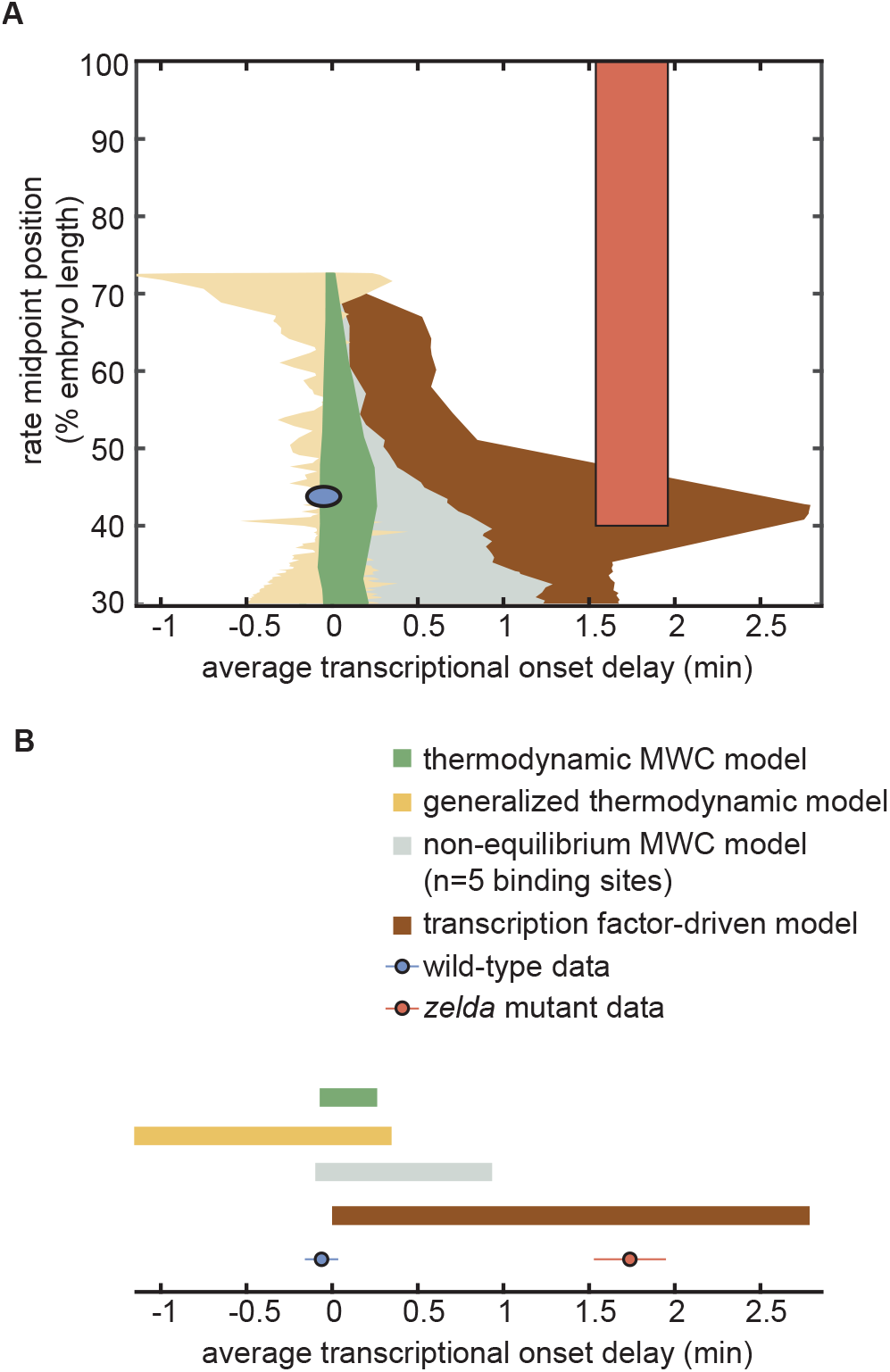
Exploration of state space. (A) Two-dimensional state-space exploration, showing the extents of state space of the wild-type (blue) and *zelda^−^* (red) data as well as of various models explored in the main text. (B) One-dimensional state-space exploration, created by projecting the two-dimensional state space in (A) for rate midpoint values between 40% and 100% onto the x-axis corresponding to the average *t_on_* delay. Areas covered by the experimental data represent the standard error of the mean.

This generalized model is much more powerful than the thermodynamic MWC model due to a lack of coupling between individual microstate weights. Whereas in the previous model the underlying parameters K_b_ and ω_b_ caused similar microstates to be related mathematically, now the statistical weights for each microstate are completely independent. Physically, this scenario can arise due to, for example, higher-order cooperativities or non-identical binding energies between binding sites (Estrada et al., 2016).

The partition function in this generalized thermodynamic model is given by the polynomial

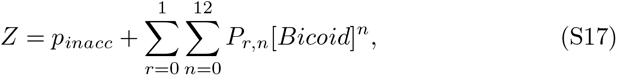

where p_inacc_ is the weight of the inaccessible state and P_r,n_ is the weight of the accessible state with r RNAP molecules bound and n Bicoid molecules bound.

The overall transcriptional initiation rate is now

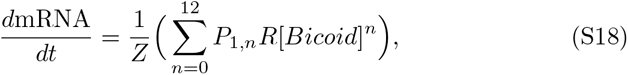

where P_1,n_ is the statistical weight of each RNAP-bound state and R is the corresponding rate of transcriptional initiation. Note that, as described above, R is still equal to R_max_, the constraint described in Section S1.3, but we no longer use the R_min_ constraint.

The resulting rate of transcriptional initiation is integrated over time to produce a simulated MS2 fluorescence trace using the same procedure as for the models presented in Sections S1.2, S7.1, and S8.1 (see Section S2.2 for details).

As with the thermodynamic MWC model, we allow for a mitotic repression time window to account for the lack of transcription early in the nuclear cycle.

#### S6.2. Generalized thermodynamic model state space exploration

Due to the high-dimensional parameter space of the generalized thermodynamic model, constraints were necessary to efficiently explore this parameter space (Section S5.1). These constraints were placed on the values of the individual microstate weights P_r,n_, based on dimensional analysis and heuristic arguments. Specifically, each weight P_r,n_ is derived from a product of binding constants K_d_ for either Bicoid or RNAP, pairwise cooperativity parameters ω, and higher-order cooperativity terms. For the purposes of these parameter constraints, we only consider the K_d_s and ωs, and ignore constraints on higher-order cooperativities. In principle, each Bicoid binding site possesses a unique K_d_ and protein-protein interaction terms ω with other Bicoid molecules and/or with RNAP. However, as described below, these biophysical parameters, once non-dimensionalized, can be constrained to reasonable values by scaling relations through a simple bounding scheme.

For illustrative purposes, consider the microstate with RNAP and one Bicoid molecule bound. Its weight depends on two independent binding constants p, b and a cooperativity term between RNAP and Bicoid ω_bp_. First, we assume that the p, b terms are non-dimensionalized, i.e. they take the form p = [RNAP]/K_p_ and b = [Bicoid]/K_b_. Although the two individual p, b terms are in principle different since RNAP and Bicoid have can different binding energies, we can be generous about the constraints and assume that the non-dimensionalized forms are both bounded below and above by 0 and 1000, respectively. This strategy is justified by assuming that neither RNAP nor Bicoid exist in concentrations three orders of magnitude above their dissociation constants, and do not exist at negative concentrations (Estrada et al., 2016). Similarly, we can be generous about any possible cooperativities and say that ω_bp_ and ω_b_ have a similar bound between 0 and 1000, thus accounting for both positive and negative cooperativities. For this state with RNAP and one Bicoid molecule bound, we can say that and thus provide a bound for the possible values that the weight P_1,1_ can take. In general, this process can be applied to enforce bounds on any microstate weight P_r,n_ through constraining the possible values of p, b, ω_bp_, and ω_b_. As a result, the weight of a microstate with more Bicoid bound (i.e. higher values of n) will have a more generous dynamic range, due to the larger powers of b and ω_b_. In this way, exploration of parameter space can be made more constrained by restricting the possible values of the microstate weights P_r,n_. The generalized thermodynamic model of transcription encompasses a larger area of the explored state space than the thermodynamic MWC model (Fig. S9A). However, as evident by the projection onto the x-axis, this model fails to capture the delays observed in zelda^−^ data (Fig. S9B, yellow rectangle).

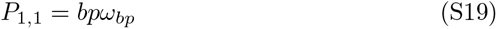

which has bounds

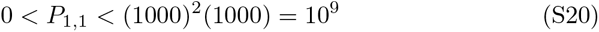

#### S6.3. Investigation of the failure of thermodynamic models

Here, we provide an intuitive explanation for why equilibrium thermodynamic models fail to recapitulate the delay in t_on_ for zelda^−^ embryos. The combination of the occupancy hypothesis and the assumption of separation of times scales described in Section S6.4 imply that the rate of transcriptional initiation at any moment in time is an instantaneous readout of the Bicoid concentration at that time point. Thus, any thermodynamic model is memoryless. Intuitively, this means that a thermodynamic model requires transcription to begin as soon as the Bicoid concentration crosses a certain “threshold” since time delays between input and output require some sense of memory. Examination of the dynamic measurements of MS2 output in zelda^−^ embryos reveals that no matter what “threshold” concentration of Bicoid is assigned for the start of transcription, the model cannot simultaneously describe two values of t_on_ corresponding to different positions along the anterior-posterior axis (Fig. S10A and B).

**Figure S10:**
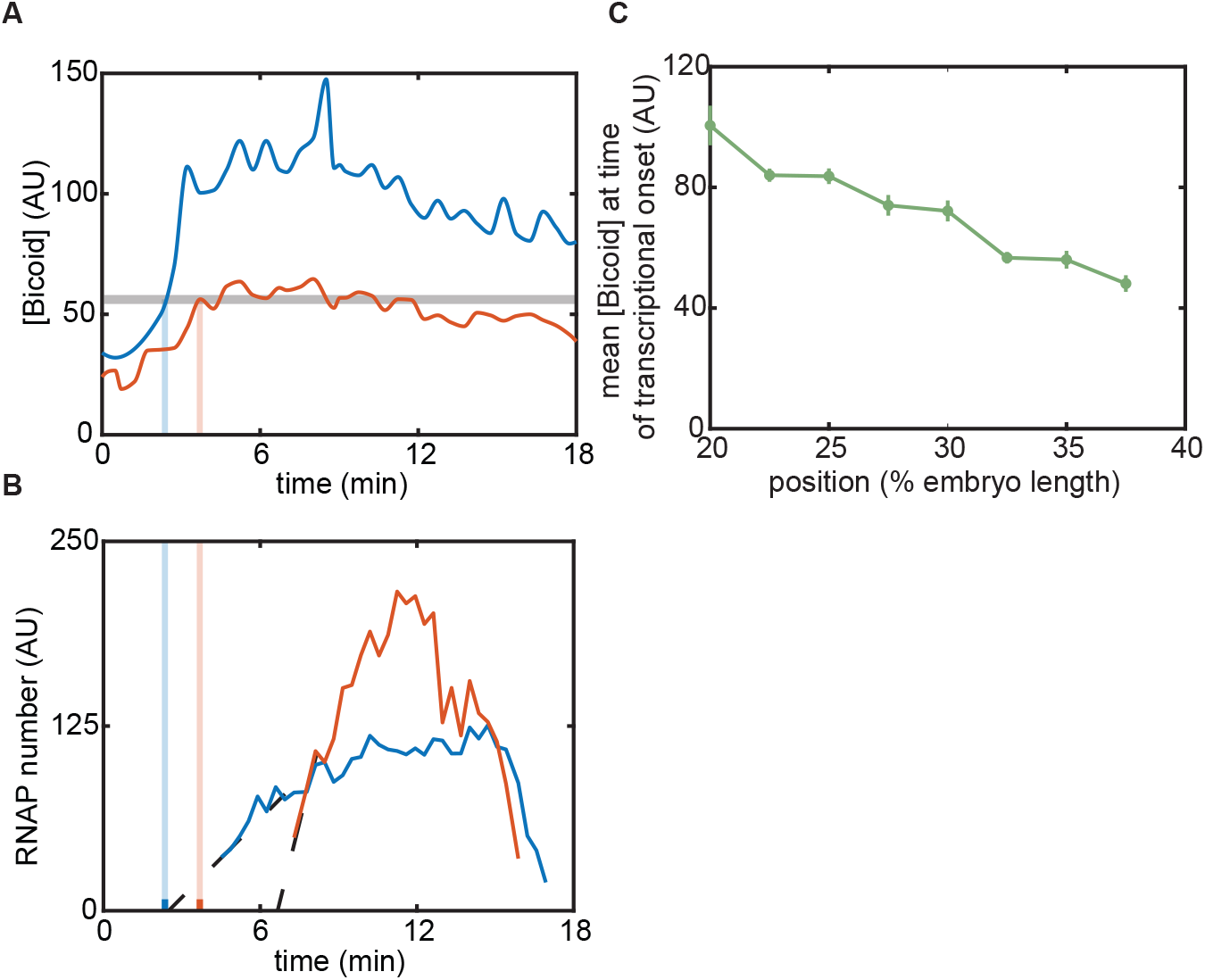
Intuition for failure of equilibrium models. (A) Mean Bicoid concentrations for two positions along the embryo (blue, red), with a “threshold” chosen to to attempt to match the corresponding *t_on_* in (B). (B) MS2 fluorescence signal for the two positions shown in for the *zelda^−^* experiment. Note that no single threshold value of Bicoid can match the timings in (A) with the transcriptional onset times in (B). (C) Mean Bicoid concentration at *t_on_* as a function of position for the *zelda^−^* data.

Another self-consistency check of a thermodynamic model is to examine the concentration of Bicoid at t_on_ for various positions along the embryo. Due to the memoryless nature of thermal equilibrium, a valid thermodynamic model predicts that, at different positions along the embryo, t_on_ will occur when Bicoid reaches the same threshold value. For the zelda^−^ data, however, the level of Bicoid at each anterior-posterior position’s t_on_ value actually decreases with increasing t_on_, suggesting the failure of the thermodynamic model (Fig. S10C). Thus, the strong position-dependent delay in t_on_ for the zelda^−^ data cannot be explained by an instantaneous Bicoid readout mechanism.

#### S6.4. Re-examining thermodynamic models of transcriptional regulation

Thermodynamic models based on equilibrium statistical mechanics can be seen as limiting cases of more general kinetic models. For example, consider simple activation, where an activator whose concentration is modulated in time regulates transcription by binding to a single site (Fig. S11). In this generic model, the presence of activator can modulate the rates of activator and RNAP binding and unbinding through the parameters α, β, γ, and δ.

In order to reduce kinetic models to thermodynamic models where the probabilities of each state are dictated by Boltzmann weights such as those in Fig. 2, four conditions must be fulfilled. First, the rate of mRNA production must be linearly related to the probability of finding RNAP bound to the promoter (Fig. S11i). This occupancy hypothesis is necessary for Eq. S2 to hold. Second, the time scales of binding and unbinding of RNAP and transcription factors must be much faster than the time scales of the concentration dynamics of these proteins (Fig. S11ii). Third, these time scales must also be much faster than the rate of transcriptional initiation and mRNA production (Fig. S11iii). Under these conditions of separation of time scales, the binding and unbinding of proteins quickly reaches steady state while the overall concentrations of these molecular players are modulated (Segel and Slemrod, 1989). Fourth, there must be no energy input into the system (Fig. S11iv). This condition demands “detailed balance” (Vilar and Leibler, 2003; Ahsendorf et al., 2014; Hill, 1985): the product of state transition rates in the clockwise direction over a closed loop is equal to the product going in the counterclockwise direction, a constraint known as the cycle condition (Estrada et al., 2016). In the case of Fig. S11, this requirement implies that

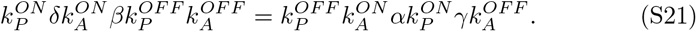

If these four conditions are met, then the system is effectively in equilibrium and the various binding states adopt probabilities that can be calculated using equilibrium statistical mechanics.

**Figure S11:**
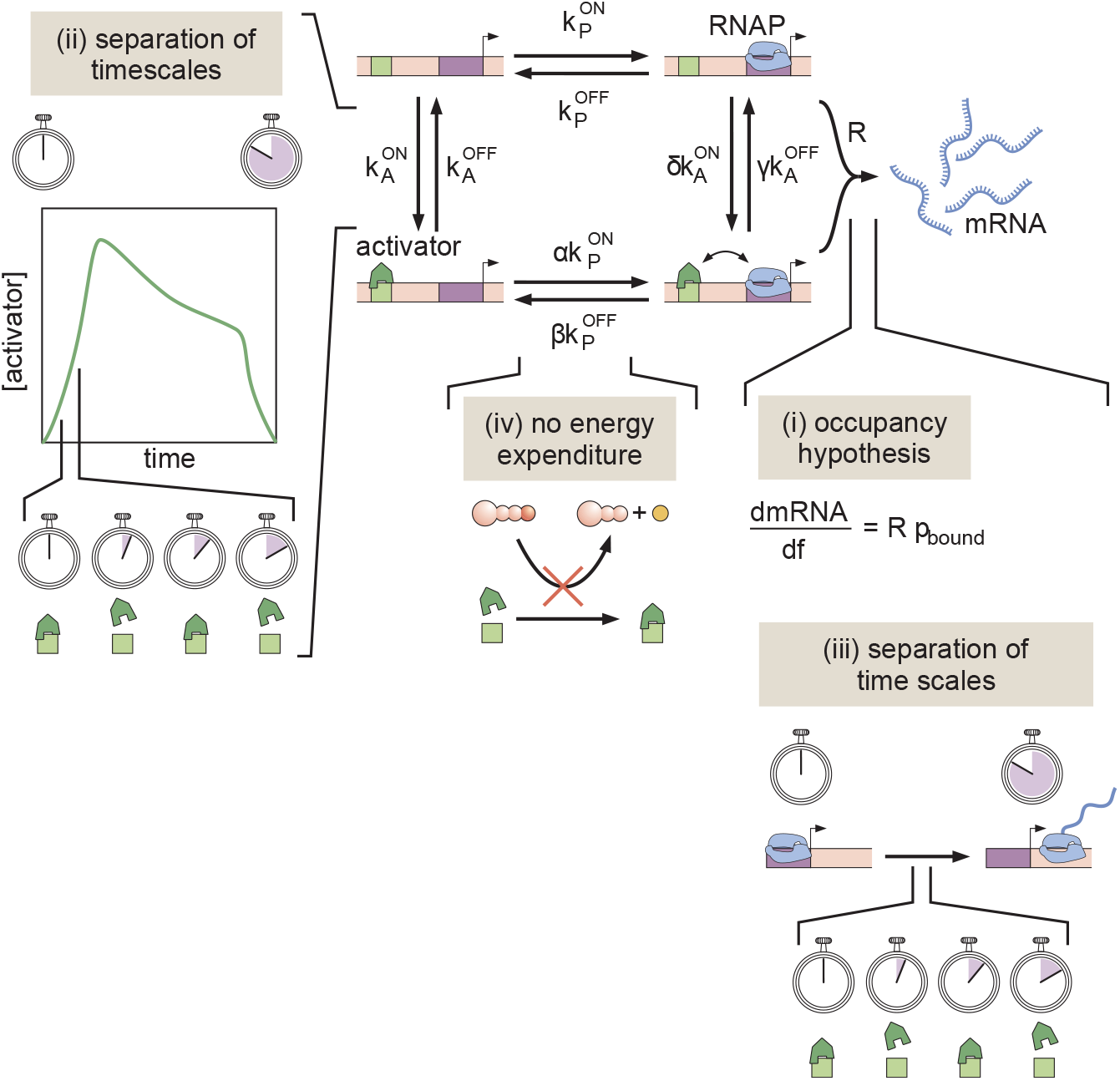
A simple kinetic model of transcriptional activation in which activator molecules influence RNAP binding kinetics. The assumptions that make it possible to turn this kinetic model into a thermodynamic one are (i) the occupancy hypothesis, (ii, iii) a separation of time scales between binding and unbinding rates, and activator and mRNA production dynamics, respectively, and (iv) no energy expenditure (detailed balance).

### S7. Non-equilibrium MWC model

#### S7.1. Non-equilibrium MWC model

The non-equilibrium MWC model is an extension of the thermodynamic MWC model presented in Section S1.2, where we now relax the assumption of separation of time scales (Fig. S11 ii and iii) and make it possible to assume, for example, that the system responds instantaneously to changes in activator concentration. Here, we explicitly simulate the full system of ordinary differential equations (ODEs) that describe the dynamics of the system out of steady state. Additionally, we allow for energy to be expended and thus do not enforce detailed balance through the cycle condition (Fig. S11iv). We still employ a mitotic repression window term, before which no transcription is allowed.

We consider a generic model with n Bicoid binding sites, and again ignore Zelda since we are only interested in recapitulating the zelda^−^ mutant data. As a result, this new model has n + 1 total binding sites which, together with the closed chromatin state, results in a total of 2^n+1^ + 1 = N microstates. In the case of six Bicoid binding sites, this results in N = 129 total microstates. We assign each microstate x_i_ a label i and describe the transition rate from state j to state i using k_ij_, where i, j range from 0 to N − 1, inclusive.

In matrix notation, we write the system of ODEs as

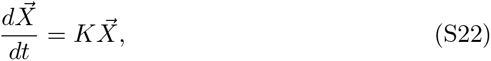

where X⃗ is a vector containing the fractional occupancy of each microstate x_i_ and K is a matrix containing all the transition rates k_ij_. Normalizing such that the sum of all the components in the vector X⃗ is unity, we now have a vector representing the instantaneous probability of being in each microstate.

To relate the occupancies of the different states to the rate of transcriptional initiation, we retain the occupancy hypothesis presented earlier: that p_bound_, the probability of being in a microstate with a bound RNAP molecule, is linearly related to the overall average transcriptional initiation rate that we determine from experimentally measurements.

For this particular system, it is helpful to define an intuitive microstate labeling system. Because the relevant physical processes are the binding and unbinding of Bicoid and RNAP molecules, we can represent any microstate in binary form, where the total number of digits is the total number of binding sites n + 1, and each digit represents an individual binding site. Our convention is to assign the first digit to the promoter, and the subsequent ones to the Bicoid sites. By assigning 0 to an unbound site and 1 to a bound site, we can rewrite each unique microstate’s label i in binary form. For example, for a model with six Bicoid sites, the label for the microstate with no RNAP bound and the first two Bicoid sites occupied is represented with

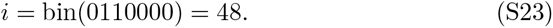

Here, bin() indicates taking the base 2 value of the binary label in the parentheses. The closed chromatin state is added manually and assigned to the last position in our binary label, x_N−1_. This convention allows us to intuitively define each unique label for the system’s microstates and provides a way to map the physical contents of a microstate with its associated label i.

In general, the overall transition matrix K can be very complex. However, we benefit from the fact that the only non-zero transitions k_ij_ are the ones that correspond to physical processes: modifying the open/closed chromatin state, and binding and unbinding of Bicoid or RNAP molecules. In this binary notation, these constraints imply that the only nonzero transitions are the ones that represent individual flips between 0 and 1, as well as between the open and closed states 0 and N − 1. The transition matrix K is then easier to write, since it is clear from the binary representation which transitions must be nonzero. Finally, diagonal elements k_ii_ are entirely constrained because they represent probability loss from a particular state i, and must be equal to the negative of the rest of the column i, such that the sum over each column in K is zero.

Given that the Bicoid concentration changes as a function of time and that we assume first-order binding kinetics, whichever rates k_ij_ correspond to Bicoid binding rates must be multiplied by this time-dependent nuclear concentration. In contrast, all off-rates are independent of Bicoid concentration. To keep subsequent parameter exploration simple, we non-dimensionalized the Bicoid concentration by rescaling it by its approximate scale. This was achieved by dividing all Bicoid concentrations by the average Bicoid concentration, calculated by averaging the mean Bicoid nuclear fluorescence across all datasets, anterior-posterior positions, and time points, yielding approximately 35 arbitrary fluorescence units. Thus, all of the transition rates k_ij_ in the model here are expressed in units of inverse minutes.

To model transcription specifically, we assumed that at the beginning of the nuclear cycle, the system is in the closed chromatin state: x_i_(t = 0) = 0 except for the closed chromatin state x_N−1_(t = 0) = 1. We simulated the full trajectory of all the microstates x_i_ over time by solving the system of ODEs given in Eq. S22. Finally, we calculated p_bound_ by summing the x_i_’s that correspond to RNAP-bound states, and then computed the subsequent transcriptional initiation rate by multiplying p_bound_ with the transcription rate R. Here, R is the same R_max_ as in Sections S1.2 and S6.1 but again we do not constrain the model using R_min_, just as in Section S6.1.

Fig. S12A shows an example of this model for a system with only one Bicoid binding site and no closed chromatin state, for simplicity, resulting in a four-state network. The binary indexing labels (shown beneath each state in light pink) can be converted into the base-10 labels (light teal) ranging from 0 to 3. The connection matrix for this system is

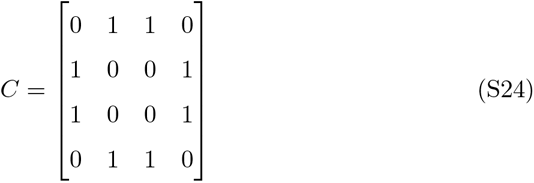

and the corresponding transition rate matrix *K* is

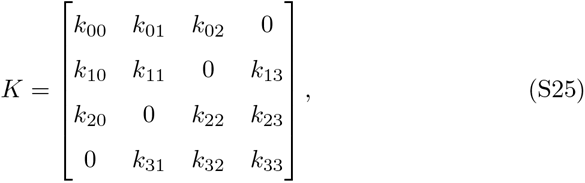

where, in this example, *k*_02_ represents the transition rate from state *j* to state *i*. The diagonal elements *k_ii_* are equal to the negative of the sum of the elements in the rest of the column in order to preserve conservation of probability. For example, *k*_00_ = *−*(*k*_10_ + *k*_20_ + *k*_30_).

With all this information in hand, we solve for the occupancy of each of the four states using the matrix ODE

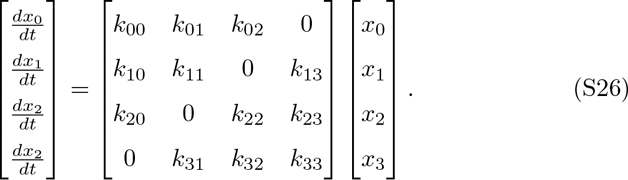

In this case, the occupancy hypothesis relates *p_bound_* to the overall transcription rate, resulting in

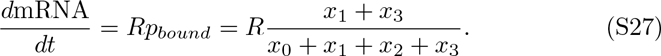

This model can produce time-dependent behavior not found in the thermodynamic models. Fig. S12B contains an example of a hypothetical input Bicoid activator concentration that switches instantaneously from zero to a finite value. In the thermodynamic models, the predicted transcriptional initiation rate also responds instantaneously (Fig. S12B, top). In contrast, for a suitable set of parameters, the non-equilibrium MWC model predicts a slow response over time (Fig. S12B, bottom).

To produce a simulated MS2 fluorescence trace, the resulting rate of mRNA production is integrated over time using the same procedure (Section S2.2) as the models presented in Sections S1.2, S6.1, and S8.1. As with the thermodynamic MWC model, we allow for a time window of mitotic repression to account for the lack of transcription early in the nuclear cycle.

**Figure S12:**
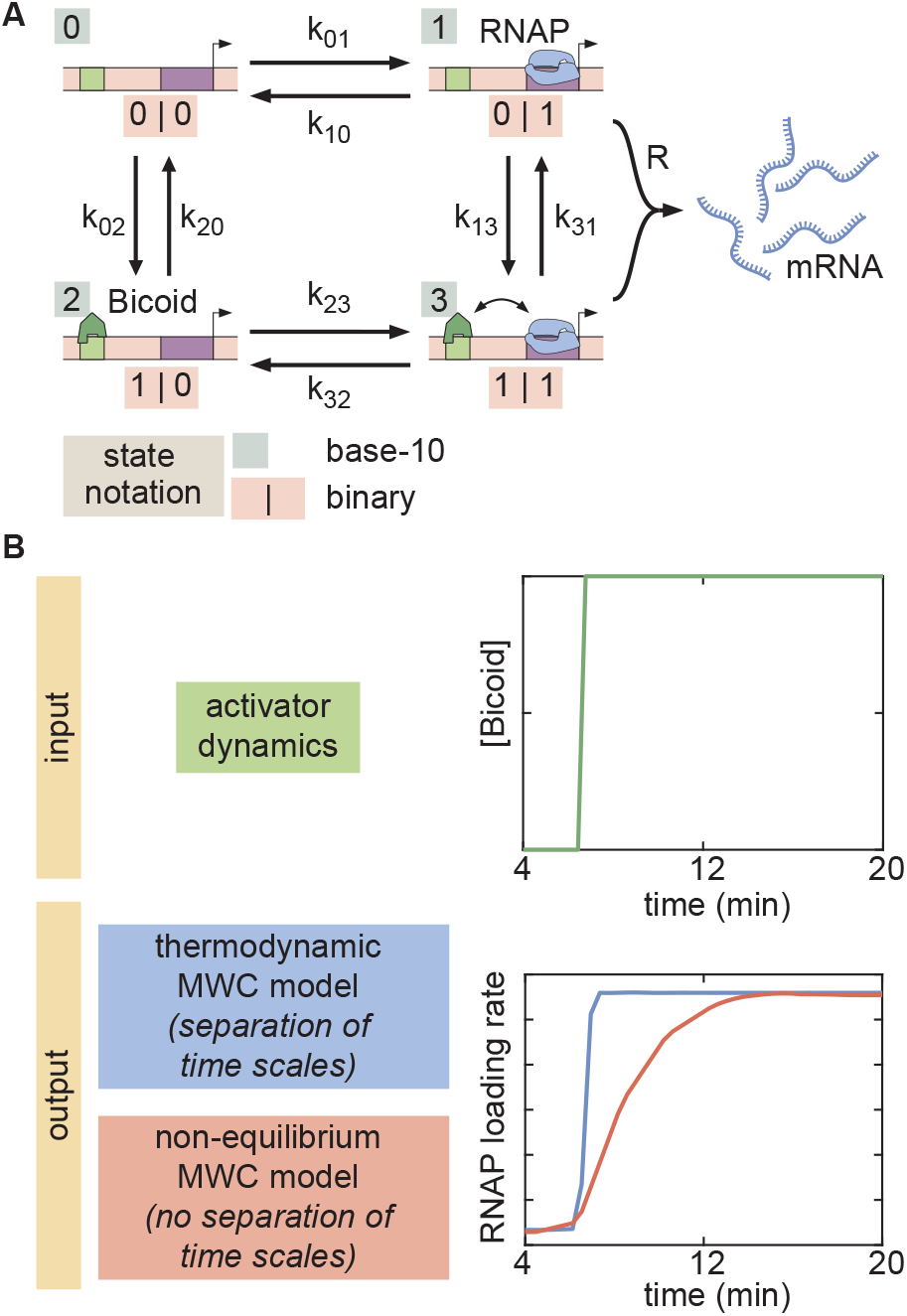
Example of a four-state time-dependent model with one Bicoid binding site and no closed chromatin state. (A) The binary label for each state (light pink) can be converted into a base-10 label for each state (light teal). The transition rates *k_ij_*are defined as the transition rate from state *i* into state *j* using this labeling system. (B) For an example input activator concentration temporal profile that is a step function, the time-dependent response is compared for the cases of separation of time scales and lack thereof. In the former, the transcriptional initiation rate responds instantaneously to the increase in activator input, while the response is slower in the latter.

#### S7.2. Non-equilibrium MWC model state space exploration

In the parameter exploration of this model (Section S5.1), the transition rates *k_ij_* were constrained with minimum and maximum values of *k_min_* = 1 and *k_max_* = 10^5^ respectively, in units of inverse minutes. These bounds were conservatively chosen using the following estimates. First, we estimate the values of the possible unbinding rates *k_off_*. We assume that RNAP and Bicoid obey the same unbinding kinetics. Estimates of *in vivo* single-molecule binding kinetics inferred from Mir et al. (2018) indicate that the lifetime of Bicoid on DNA is on the order of 3 *s^−^*^1^. Second, we estimate the values of the possible on-rates *k_on_* using the classic Berg-Purcell equation for the case of a diffusion-limited binding to a perfectly absorbing spherical receptor (Berg and Purcell, 1977). In this case, the on-rate of molecule binding is given by

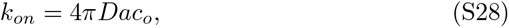

where *D* is the diffusion coefficient of the molecule, *a* is the estimated size of the spherical receptor, and *c*_0_ is the background concentration of the molecular species. Since here we are talking about transcription factor binding to a Bicoid binding site, we assume *a* to be on the order of 5 nm. We assume that RNAP and Bicoid obey the same diffusion characteristics, leading to a diffusion coefficient of approximately 0.3 *µm*^2^*s^−^*^1^ (Gregor et al., 2007b). Finally, Bicoid is is present at concentrations between 10 nM and 55 nM in the nucleus (Gregor et al., 2007a), and we assume that nuclear RNAP concentrations exist within the same range. Plugging these values into Eq. S28 yields estimates for the maximum and minimum on-rates:

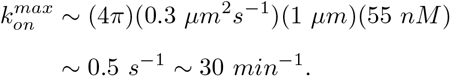

and

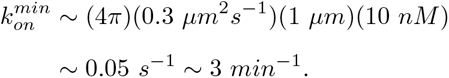

Thus, our maximum and minimum transition rate bounds of *k_min_* = 1 *min^−^*^1^ and *k_max_* = 10^5^ *min^−^*^1^ lie outside these estimated binding and unbinding rates.

One caveat of the state-space exploration approach is that the high dimensionality of the non-equilibrium MWC model prevented us from calculating the full state-space boundary using six Bicoid binding sites. Due to computational costs, we were only able to accurately produce a state-space boundary for this model (Section S7.1) using five Bicoid binding sites. Running the exploration for a model with six Bicoid binding sites took over two weeks on our own server, and the algorithm had not noticeably converged in the end.

The results of the state space exploration for the non-equilibrium MWC model using five Bicoid binding sites resulted in larger average *t_on_* delays than the thermodynamic models (Sections S1.2 and S6.1). However, this model, like those, failed to reproduce the delays observed in *zelda^−^* data (Fig. S9B, gray bar).

### S8. Transcription factor-driven model of chromatin accessibility

#### S8.1. Transcription factor-driven model of chromatin accessibility

The transcription factor-driven model of chromatin accessibility is a slight modification of the thermodynamic MWC model (Section S1.2) that replaces the MWC mechanism of chromatin transitions with a direct driving action due to Bicoid and Zelda. Here, we retain the idea of inaccessible vs. accessible states, but no longer demand that these states be in thermodynamic equilibrium. Instead, the system begins in the inaccessible state and undergoes a series of m identical, slow, and effectively irreversible transitions to the accessible state. Once these transitions into the accessible state occur, the system can rapidly and reversibly transition into all of its accessible microstates such that the probability of the system being in any of these microstates is described by thermodynamic equilibrium. The accessible states are governed by the same rules and parameters as the thermodynamic MWC model (Section S1.2), albeit without the Δε_chrom_ parameter since now the transition from the inaccessible to accessible state is unidirectional.

Additionally, we consider two possible contributions for these irreversible transitions: a Bicoid-dependent pathway and a Zelda-dependent pathway. We assume the transition rates to be first-order in Bicoid and Zelda, respectively, such that

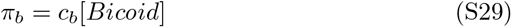

and

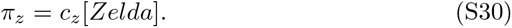

Here, π_b_ is the Bicoid-dependent contribution to the transition rates and π_z_ is the corresponding Zelda-dependent contribution. There are two input parameters c_b_ and c_z_ that give the relative speed of each transition rate contribution. The overall rate π of each irreversible transition is given by the sum

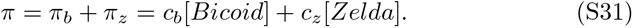

Because the accessible states are in thermodynamic equilibrium with each other, we can effectively treat them as a single state and describe the entire system with m + 1 states, corresponding to the inaccessible, intermediate, and accessible states. We label the inaccessible state with 0, the m − 1 intermediate states with 1 through m − 1, and the final accessible state with m. Thus, we describe the probability p_i_of the system being in the state i with the probability vector P⃗

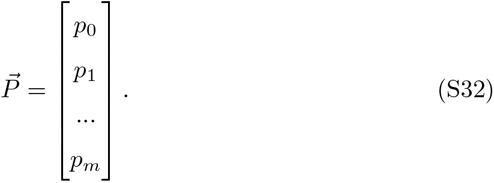

Calculating the overall RNAP loading rate then simply corresponds to rescaling *p_bound_* with the overall probability *p_m_*(*t*) of being in the accessible state: dmRNA

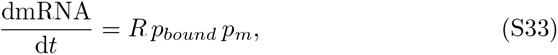

where *R* is the same maximum rate used in Section S1.2. Note that *p_m_*(*t*) is a time-dependent quantity that changes over time. To calculate *p_m_*(*t*), we solve the corresponding system of ODEs that describes the time evolution of *P⃗*

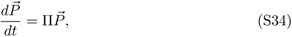

where Π is the transition rate matrix describing the time evolution of the system. Π, by definition, is a square matrix with dimension *m* + 1. Given the initial condition that the system begins in the inaccessible state

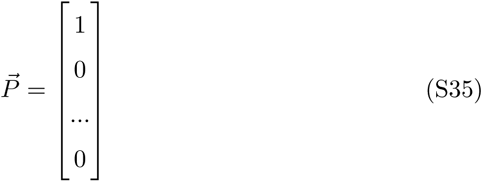

the system of ODEs can be solved to find the probability of being in the accessible state *p_m_*(*t*). For example, for *m* = 3 irreversible steps, Π takes the form

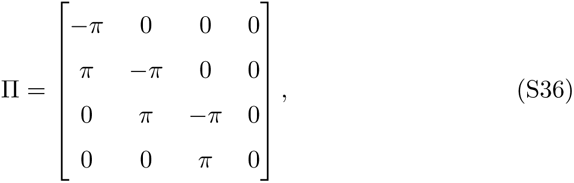

where *π* is given by Eq. S31.

For simplicity, the time evolution of *P⃗* was solved using MATLAB’s ode15s solver.

With the probability *p_m_*(*t*) of the system being in the accessible state calculated, we now calculate the probability *p_bound_* of RNAP bound to the promoter in the accessible states, which lie in thermodynamic equilibrium with each other. Because we now only have accessible states, the partition function is

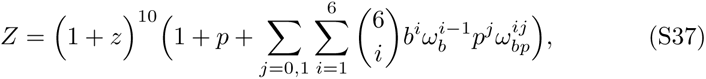

where *z*, *p*, and *b* correspond to the non-dimensionalized concentrations of Zelda, RNAP, and Bicoid, respectively, and *ω_b_* and *ω_bp_* are the cooperativities between

Bicoid molecules and between Bicoid and RNAP, respectively. Thus, the overall transcriptional initiation rate is given by

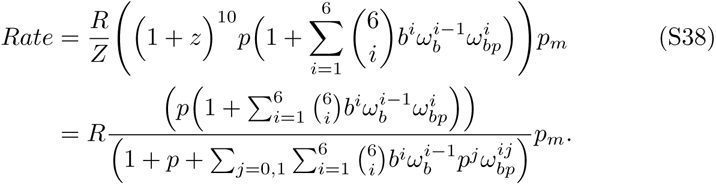

Due to the lack of the inaccessible state in the partition function and because we assume that Zelda does not directly interact with Bicoid or RNAP, now the presence of Zelda mathematically separates out so that only Bicoid influences transcription. The calculation above is a standard equilibrium statistical mechanical calculation, except that we have weighted the final result with *p_m_*(*t*), the probability of being in the accessible states. The resulting rate is integrated to produce a simulated MS2 fluorescence trace using the same procedure (Section S2.2) as the models presented in Sections S1.2, S6.1, and S7.1.

Interestingly, we found that a mitotic repression term was not necessary to recapitulate the data, since the presence of intermediary states produced the necessary delay to explain the experimentally observed *t_on_* values in the data (Fig. 4D and G, points).

In order to sufficiently explain the data, we found that a minimum of *m* = 5 irreversible steps was necessary. Fig. S13A and B show the results of fitting this model to the observed rates of RNAP loading and *t_on_* for the wild-type and *zelda^−^* data, for increasing values of *m* (wild-type results not shown, since all values of *m* easily explained the wild-type data). We see that while lower values of *m* do a poor job of recapitulating the data, once we reach *m* = 5 the model sufficiently predicts the experimental data within experimental error. For values of *m* higher than 5, explanatory power increases marginally. Considering the parameter exploration of this model (Section S8.2) highlights the necessity of having at least *m* = steps.

**Figure S13:**
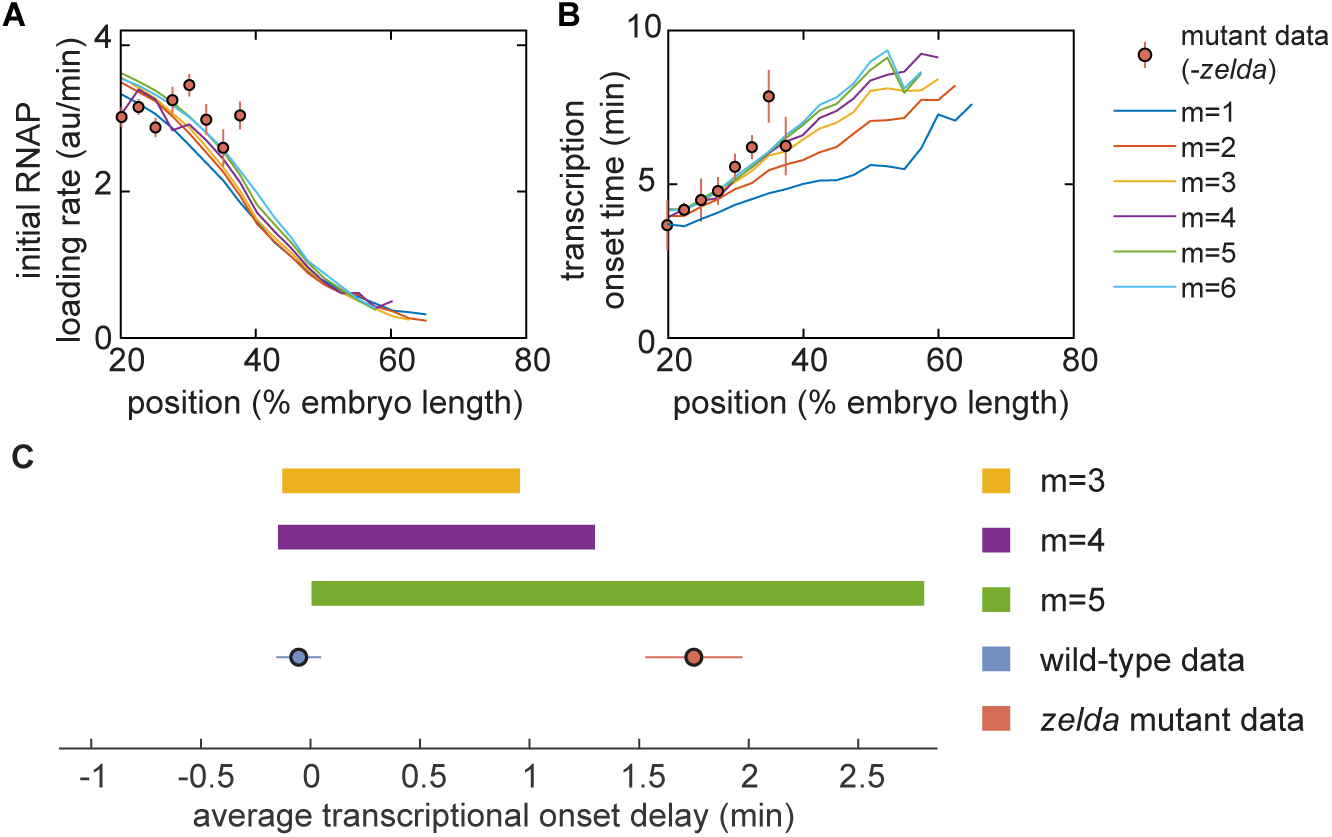
Testing the transcription factor-driven model of chromatin accessibility. (A,B) Best-fit results of the non-equilibrium MWC model to the mutant *zelda^−^* data. (A) initial RNAP loading rates, and (B) *t_on_*, for varying numbers *m* of irreversible steps. (C) Parameter exploration in average *t_on_* delay space for increasing values of *m*.

#### S8.2. Transcription factor-driven model of chromatin accessibility state space exploration

In the parameter exploration of this model (Section S5.1), the parameters were constrained as

- c_b_ > 0
- c_z_ > 0.

The parameters shared with the thermodynamic MWC model retained the constraints described in Section S1.3.

Fig. S13C shows the parameter space explorations (see Section S5.1) of this transcription factor-driven model for increasing numbers of intermediate steps *m*. Not until *m* = 5 does the model explain the average *t_on_* delay for both the wild-type and *zelda^−^* data, indicating that *m* = 5 is the minimum number of irreversible steps necessary. In the state space exploration shown in Fig. S9, the number of irreversible steps was fixed at *m* = 5.

Unlike the other models investigated (Sections S1.2, S6.1, and S7.1), the transcription factor-driven model of chromatin accessibility occupied a region in state space that encompassed both the wild-type and *zelda^−^* data (Fig. S9, brown rectangle).

### S9. Supplementary Videos

S1. **Video 1. Measurement of eGFP-Bicoid.** Movie of eGFP-Bicoid fusion in an embryo in nuclear cycle 13. Time is defined with respect to the previous anaphase.

S2. **Video 2. Measurement of Zelda-sfGFP.** Movie of Zelda-sfGFP fusion in an embryo in nuclear cycle 13. Time is defined with respect to the previous anaphase.

S3. **Video 3. Measurement of MS2 fluorescence in a wild-type background.** Movie of MS2 fluorescent spots in a wild-type background embryo in nuclear cycle 13. Time is defined with respect to the previous anaphase.

S4. **Video 4. Measurement of MS2 fluorescence in a** *zelda^−^* **background.** Movie of MS2 fluorescent spots in a *zelda^−^* background embryo in nuclear cycle 13. Time is defined with respect to the previous anaphase.

S5. **Video 5. Transcriptionally active nuclei in a wild-type background.** Movie of MS2 fluorescent spots in a wild-type background embryo in nuclear cycle 13, with transcriptionally active nuclei labeled with an overlay. Time is defined with respect to the previous anaphase.

S6. **Video 6. Transcriptionally active nuclei in a** *zelda^−^* **background.** Movie of MS2 fluorescent spots in a *zelda^−^* background embryo in nuclear cycle 13, with transcriptionally active nuclei labeled with an overlay. Time is defined with respect to the previous anaphase.

## References

Ackers, G.K., Johnson, A.D., Shea, M.A., 1982. Quantitative model for gene regulation by lambda phage repressor. Proc Natl Acad Sci U S A 79, 1129–33.

Adams, C.C., Workman, J.L., 1995. Binding of disparate transcriptional activators to nucleosomal DNA is inherently cooperative. Molecular and Cellular Biology 15, 1405–1421. doi:10.1128/mcb.15.3.1405.

Ahsendorf, T., Wong, F., Eils, R., Gunawardena, J., 2014. A framework for modelling gene regulation which accommodates non-equilibrium mechanisms. BMC Biol 12, 102. doi:10.1186/s12915-014-0102-4.

Bai, L., Ondracka, A., Cross, F.R., 2011. Multiple sequence-specific factors generate the nucleosome-depleted region on cln2 promoter. Mol Cell 42, 465–76. doi:10.1016/j.molcel.2011.03.028.

Bakk, A., Metzler, R., Sneppen, K., 2004. Sensitivity of or in phage lambda. Biophys J 86, 58–66.

Berg, H.C., Purcell, E.M., 1977. Physics of chemoreception. Biophys J 20, 193–219. doi:S0006-3495(77)85544-6[pii]10.1016/S0006-3495(77)85544-6.

Berrocal, A., Lammers, N.C., Garcia, H.G., Eisen, M.B., 2018. Kinetic sculpting of the seven stripes of the drosophila even-skipped gene. bioRxiv doi:10.1101/335901.

Bertrand, E., Chartrand, P., Schaefer, M., Shenoy, S.M., Singer, R.H., Long, R.M., 1998. Localization of ash1 mrna particles in living yeast. Mol Cell 2, 437–45. doi:S1097-2765(00)80143-4[pii].

Bintu, L., Buchler, N.E., Garcia, H.G., Gerland, U., Hwa, T., Kondev, J., Kuhlman, T., Phillips, R., 2005a. Transcriptional regulation by the numbers: applications. Curr Opin Genet Dev 15, 125–35.

Bintu, L., Buchler, N.E., Garcia, H.G., Gerland, U., Hwa, T., Kondev, J., Phillips, R., 2005b. Transcriptional regulation by the numbers: models. Curr Opin Genet Dev 15, 116–24.

Blythe, S.A., Wieschaus, E.F., 2016. Establishment and maintenance of heritable chromatin structure during early drosophila embryogenesis. Elife 5. doi:10.7554/eLife.20148.

Bolouri, H., Davidson, E.H., 2003. Transcriptional regulatory cascades in development: initial rates, not steady state, determine network kinetics. Proc Natl Acad Sci U S A 100, 9371–6. doi:10.1073/pnas.1533293100.

Bothma, J.P., Garcia, H.G., Esposito, E., Schlissel, G., Gregor, T., Levine, M., 2014. Dynamic regulation of eve stripe 2 expression reveals transcriptional bursts in living drosophila embryos. Proc Natl Acad Sci U S A 111, 1059810603. doi:10.1073/pnas.1410022111.

Bothma, J.P., Garcia, H.G., Ng, S., Perry, M.W., Gregor, T., Levine, M., 2015. Enhancer additivity and non-additivity are determined by enhancer strength in the drosophila embryo. Elife 4. doi:10.7554/eLife.07956.

Brewster, R.C., Jones, D.L., Phillips, R., 2012. Tuning Promoter Strength through RNA Polymerase Binding Site Design in Escherichia coli. PLoS Computational Biology 8. doi:10.1371/journal.pcbi.1002811.

Brewster, R.C., Weinert, F.M., Garcia, H.G., Song, D., Rydenfelt, M., Phillips, R., 2014. The transcription factor titration effect dictates level of gene expression. Cell 156, 1312–23. doi:10.1016/j.cell.2014.02.022.

Buchler, N.E., Gerland, U., Hwa, T., 2003. On schemes of combinatorial transcription logic. Proc Natl Acad Sci U S A 100, 5136–41.

Chen, H., Xu, Z., Mei, C., Yu, D., Small, S., 2012. A system of repressor gradients spatially organizes the boundaries of bicoid-dependent target genes. Cell 149, 618–29. doi:10.1016/j.cell.2012.03.018.

Chure, G., Razo-Mejia, M., Belliveau, N.M., Einav, T., Kaczmarek, Z.A., Barnes, S.L., Lewis, M., Phillips, R., 2019. Predictive shifts in free energy couple mutations to their phenotypic consequences. Proc Natl Acad Sci U S A 116, 18275–18284. doi:10.1073/pnas.1907869116.

Cui, L., Murchland, I., Shearwin, K.E., Dodd, I.B., 2013. Enhancer-like long-range transcriptional activation by lambda ci-mediated dna looping. Proc Natl Acad Sci U S A 110, 2922–7. doi:10.1073/pnas.1221322110.

Culkin, J., de Bruin, L., Tompitak, M., Phillips, R., Schiessel, H., 2017. The role of dna sequence in nucleosome breathing. Eur Phys J E Soft Matter 40, 106. doi:10.1140/epje/i2017-11596-2.

Desponds, J., Tran, H., Ferraro, T., Lucas, T., Perez Romero, C., Guillou, A., Fradin, C., Coppey, M., Dostatni, N., Walczak, A.M., 2016. Precision of readout at the hunchback gene: Analyzing short transcription time traces in living fly embryos. PLoS Comput Biol 12, e1005256. doi:10.1371/journal.pcbi.1005256.

Driever, W., Nusslein-Volhard, C., 1988a. The bicoid protein determines position in the drosophila embryo in a concentration-dependent manner. Cell 54, 95–104.

Driever, W., Nusslein-Volhard, C., 1988b. A gradient of bicoid protein in drosophila embryos. Cell 54, 83–93.

Driever, W., Nusslein-Volhard, C., 1989. The bicoid protein is a positive regulator of hunchback transcription in the early drosophila embryo. Nature 337, 138–43. doi:10.1038/337138a0.

Driever, W., Thoma, G., Nusslein-Volhard, C., 1989. Determination of spatial domains of zygotic gene expression in the drosophila embryo by the affinity of binding sites for the bicoid morphogen. Nature 340, 363–7. doi:10.1038/340363a0.

Dufourt, J., Trullo, A., Hunter, J., Fernandez, C., Lazaro, J., Dejean, M., Morales, L., Nait-Amer, S., Schulz, K.N., Harrison, M.M., Favard, C., Radulescu, O., Lagha, M., 2018. Temporal control of gene expression by the pioneer factor zelda through transient interactions in hubs. Nat Commun 9, 5194. doi:10.1038/s41467-018-07613-z.

Edgar, B.A., Odell, G.M., Schubiger, G., 1987. Cytoarchitecture and the patterning of fushi tarazu expression in the drosophila blastoderm. Genes Dev 1, 1226–37.

Edgar, B.A., Schubiger, G., 1986. Parameters controlling transcriptional activation during early drosophila development. Cell 44, 871–7. doi:0092-8674(86)90009-7[pii].

Endy, D., 2005. Foundations for engineering biology. Nature 438, 449–53. doi:nature04342[pii]10.1038/nature04342.

Estrada, J., Wong, F., DePace, A., Gunawardena, J., 2016. Information integration and energy expenditure in gene regulation. Cell 166, 234–44. doi:10.1016/j.cell.2016.06.012.

Fakhouri, W.D., Ay, A., Sayal, R., Dresch, J., Dayringer, E., Arnosti, D.N., 2010. Deciphering a transcriptional regulatory code: modeling short-range repression in the drosophila embryo. Mol Syst Biol 6, 341. doi:msb200997[pii] 10.1038/msb.2009.97.

Foo, S.M., Sun, Y., Lim, B., Ziukaite, R., O’Brien, K., Nien, C.Y., Kirov, N., Shvartsman, S.Y., Rushlow, C.A., 2014. Zelda potentiates morphogen activity by increasing chromatin accessibility. Current Biology 24, 1341–1346. doi:10.1016/j.cub.2014.04.032.

Fu, D., Ma, J., 2005. Interplay between positive and negative activities that influence the role of bicoid in transcription. Nucleic Acids Res 33, 3985–93. doi:10.1093/nar/gki691.

Fu, D., Wen, Y., Ma, J., 2004. The co-activator creb-binding protein participates in enhancer-dependent activities of bicoid. J Biol Chem 279, 48725–33. doi:10.1074/jbc.M407066200.

Fussner, E., Ching, R.W., Bazett-Jones, D.P., 2011. Living without 30nm chromatin fibers. Trends in Biochemical Sciences 36, 1–6. doi:10.1016/j.tibs.2010.09.002.

Garcia, H.G., Gregor, T., 2018. Live Imaging of mRNA Synthesis in Drosophila. Springer New York, New York, NY. pp. 349–357.

Garcia, H.G., Kondev, J., Orme, N., Theriot, J.A., Phillips, R., 2007. A first exposure to statistical mechanics for life scientists. arXiv preprint arXiv:0708.1899.

Garcia, H.G., Phillips, R., 2011. Quantitative dissection of the simple repression input-output function. Proc Natl Acad Sci U S A 108, 12173–8. doi:1015616108[pii]10.1073/pnas.1015616108.

Garcia, H.G., Sanchez, A., Boedicker, J.Q., Osborne, M., Gelles, J., Kondev, J., Phillips, R., 2012. Operator sequence alters gene expression independently of transcription factor occupancy in bacteria. Cell Rep 2, 150–61. doi:10.1016/j.celrep.2012.06.004.

Garcia, H.G., Tikhonov, M., Lin, A., Gregor, T., 2013. Quantitative imaging of transcription in living drosophila embryos links polymerase activity to patterning. Curr Biol 23, 2140–5. doi:10.1016/j.cub.2013.08.054.

Gertz, J., Siggia, E.D., Cohen, B.A., 2009. Analysis of combinatorial cisregulation in synthetic and genomic promoters. Nature 457, 215–8.

Giorgetti, L., Siggers, T., Tiana, G., Caprara, G., Notarbartolo, S., Corona, T., Pasparakis, M., Milani, P., Bulyk, M.L., Natoli, G., 2010. Noncooperative interactions between transcription factors and clustered dna binding sites enable graded transcriptional responses to environmental inputs. Mol Cell 37, 418–28. doi:10.1016/j.molcel.2010.01.016.

Gottesfeld, J.M., Forbes, D.J., 1997. Mitotic repression of the transcriptional machinery. Trends Biochem Sci 22, 197–202.

Gregor, T., Tank, D.W., Wieschaus, E.F., Bialek, W., 2007a. Probing the limits to positional information. Cell 130, 153–64.

Gregor, T., Wieschaus, E.F., McGregor, A.P., Bialek, W., Tank, D.W., 2007b. Stability and nuclear dynamics of the bicoid morphogen gradient. Cell 130, 141–52.

Hamm, D.C., Larson, E.D., Nevil, M., Marshall, K.E., Bondra, E.R., Harrison, M.M., 2017. A conserved maternal-specific repressive domain in Zelda revealed by Cas9-mediated mutagenesis in Drosophila melanogaster. PLOS Genetics 13, e1007120. URL: http://journals.plos.org/plosgenetics/article?id=10.1371/journal.pgen.1007120, doi:10.1371/journal.pgen.1007120.

Hammar, P., Wallden, M., Fange, D., Persson, F., Baltekin, O., Ullman, G., Leroy, P., Elf, J., 2014. Direct measurement of transcription factor dissociation excludes a simple operator occupancy model for gene regulation. Nat Genet 46, 405–8. doi:10.1038/ng.2905.

Hannon, C.E., Blythe, S.A., Wieschaus, E.F., 2017. Concentration dependent chromatin states induced by the bicoid morphogen gradient. Elife 6. doi:10.7554/eLife.28275.

Hansen, A.S., O’Shea, E.K., 2015. Cis Determinants of Promoter Threshold and Activation Timescale. Cell Reports 12, 1226–1233. URL: http://dx.doi.org/10.1016/j.celrep.2015.07.035, doi:10.1016/j.celrep.2015.07.035.

Harrison, M.M., Li, X.Y., Kaplan, T., Botchan, M.R., Eisen, M.B., 2011. Zelda binding in the early drosophila melanogaster embryo marks regions subsequently activated at the maternal-to-zygotic transition. PLoS Genet 7, e1002266. doi:10.1371/journal.pgen.1002266.

He, H.H., Meyer, C.A., Shin, H., Bailey, S.T., Wei, G., Wang, Q., Zhang, Y., Xu, K., Ni, M., Lupien, M., Mieczkowski, P., Lieb, J.D., Zhao, K., Brown, M., Liu, X.S., 2010. Nucleosome dynamics define transcriptional enhancers. Nat Genet 42, 343–7. doi:10.1038/ng.545.

Hertz, G.Z., Hartzell, G.W., Stormo, G.D., 1990. Identification of consensus patterns in unaligned DNA sequences known to be functionally related. Bioinformatics 6, 81–92. doi:10.1093/bioinformatics/6.2.81.

Hertz, G.Z., Stormo, G.D., 1999. Identifying DNA and protein patterns with statistically significant alignments of multiple sequences. Bioinformatics (Oxford, England) 15, 563–577.

Hill, T.L., 1985. Cooperativity theory in biochemistry: steady-state and equilibrium systems. Springer-Verlag, New York.

Jaeger, J., Blagov, M., Kosman, D., Kozlov, K.N., Manu, Myasnikova, E., Surkova, S., Vanario-Alonso, C.E., Samsonova, M., Sharp, D.H., Reinitz, J., 2004a. Dynamical analysis of regulatory interactions in the gap gene system of drosophila melanogaster. Genetics 167, 1721–37. doi:10.1534/genetics.104.027334.

Jaeger, J., Surkova, S., Blagov, M., Janssens, H., Kosman, D., Kozlov, K.N., Manu, Myasnikova, E., Vanario-Alonso, C.E., Samsonova, M., Sharp, D.H., Reinitz, J., 2004b. Dynamic control of positional information in the early drosophila embryo. Nature 430, 368–71. doi:10.1038/nature02678.

Kanodia, J.S., Liang, H.L., Kim, Y., Lim, B., Zhan, M., Lu, H., Rushlow, C.A., Shvartsman, S.Y., 2012. Pattern formation by graded and uniform signals in the early Drosophila embryo. Biophysical Journal 102, 427–433. URL: http://dx.doi.org/10.1016/j.bpj.2011.12.042, doi:10.1016/j.bpj.2011.12.042.

Keymer, J.E., Endres, R.G., Skoge, M., Meir, Y., Wingreen, N.S., 2006. Chemosensing in escherichia coli: two regimes of two-state receptors. Proc Natl Acad Sci U S A 103, 1786–91. doi:0507438103[pii]10.1073/pnas.0507438103.

Kim, H., O’Shea, E., 2008. A quantitative model of transcription factor-activated gene expression. Nat Struct Mol Biol 15, 1192–1198.

Lam, F.H., Steger, D.J., O’Shea, E.K., 2008. Chromatin decouples promoter threshold from dynamic range. Nature 453, 246–50. doi:10.1038/nature06867.

Lammers, N.C., Galstyan, V., Reimer, A., Medin, S.A., Wiggins, C.H., Garcia, H.G., 2019. Multimodal transcriptional control of pattern formation in embryonic development. Proc Natl Acad Sci U S A doi:10.1073/pnas.1912500117.

Levine, M., 2010. Transcriptional enhancers in animal development and evolution. doi:10.1016/j.cub.2010.06.070.

Li, C., Cesbron, F., Oehler, M., Brunner, M., Hofer, T., 2018. Frequency modulation of transcriptional bursting enables sensitive and rapid gene regulation. Cell Syst 6, 409–423 e11. doi:10.1016/j.cels.2018.01.012.

Li, G.W., Burkhardt, D., Gross, C., Weissman, J.S., 2014a. Quantifying absolute protein synthesis rates reveals principles underlying allocation of cellular resources. Cell 157, 624–35. doi:10.1016/j.cell.2014.02.033.

Li, X.Y., Eisen, M.B., 2018. Zelda potentiates transcription factor binding to zygotic enhancers by increasing local chromatin accessibility during early ¡em¿drosophila melanogaster¡/em¿ embryogenesis. bioRxiv, 380857 doi:10.1101/380857.

Li, X.Y., Harrison, M.M., Villalta, J.E., Kaplan, T., Eisen, M.B., 2014b. Establishment of regions of genomic activity during the drosophila maternal to zygotic transition. Elife 3. doi:10.7554/eLife.03737.

Li, X.Y., Thomas, S., Sabo, P.J., Eisen, M.B., Stamatoyannopoulos, J.A., Biggin, M.D., 2011. The role of chromatin accessibility in directing the widespread, overlapping patterns of Drosophila transcription factor binding. Genome Biology 12. doi:10.1186/gb-2011-12-4-r34.

Liang, H.L., Nien, C.Y., Liu, H.Y., Metzstein, M.M., Kirov, N., Rushlow, C., 2008. The zinc-finger protein zelda is a key activator of the early zygotic genome in drosophila 456, 400–403. URL: http://www.ncbi.nlm.nih.gov/pmc/articles/PMC2597674/, doi:10.1038/nature07388.

Little, S.C., Tikhonov, M., Gregor, T., 2013. Precise developmental gene expression arises from globally stochastic transcriptional activity. Cell 154, 789–800. doi:10.1016/j.cell.2013.07.025.

Liu, F., Morrison, A.H., Gregor, T., 2013. Dynamic interpretation of maternal inputs by the drosophila segmentation gene network. Proc Natl Acad Sci U S A 110, 6724–9. doi:10.1073/pnas.1220912110.

Lucas, T., Ferraro, T., Roelens, B., De Las Heras Chanes, J., Walczak, A.M., Coppey, M., Dostatni, N., 2013. Live imaging of bicoid-dependent transcription in drosophila embryos. Curr Biol 23, 2135–9. doi:10.1016/j.cub.2013.08.053.

Margolis, J.S., Borowsky, M.L., Steingrimsson, E., Shim, C.W., Lengyel, J.A., Posakony, J.W., 1995. Posterior stripe expression of hunchback is driven from two promoters by a common enhancer element. Development 121, 3067–77.

Martins, B.M., Swain, P.S., 2011. Trade-offs and constraints in allosteric sensing. PLoS Comput Biol 7, e1002261. doi:10.1371/journal.pcbi.1002261.

Marzen, S., Garcia, H.G., Phillips, R., 2013. Statistical mechanics of Monod-Wyman-Changeux (MWC) models. Journal of Molecular Biology 425, 1433–1460. URL: http://dx.doi.org/10.1016/j.jmb.2013.03.013, doi:10.1016/j.jmb.2013.03.013.

Miller, J.A., Widom, J., 2003. Collaborative competition mechanism for gene activation in vivo. Mol Cell Biol 23, 1623–32.

Mir, M., Stadler, M.R., Ortiz, S.A., Hannon, C.E., Harrison, M.M., Darzacq, X., Eisen, M.B., 2018. Dynamic multifactor hubs interact transiently with sites of active transcription in drosophila embryos. Elife 7. doi:10.7554/eLife.40497.

Mirny, L.A., 2010. Nucleosome-mediated cooperativity between transcription factors. Proc Natl Acad Sci U S A 107, 22534–9. doi:0913805107[pii]10.1073/pnas.0913805107.

Monod, J., Wyman, J., Changeux, J.P., 1965. On the nature of allosteric transitions: A plausible model. J Mol Biol 12, 88–118.

Narula, J., Igoshin, O.A., 2010. Thermodynamic models of combinatorial gene regulation by distant enhancers. IET Syst Biol 4, 393. doi:10.1049/iet-syb.2010.0010.

Nien, C.Y., Liang, H.L., Butcher, S., Sun, Y., Fu, S., Gocha, T., Kirov, N., Manak, J.R., Rushlow, C., 2011. Temporal coordination of gene networks by Zelda in the early Drosophila embryo. PLoS Genetics 7. doi:10.1371/journal.pgen.1002339.

Park, J., Estrada, J., Johnson, G., Vincent, B.J., Ricci-Tam, C., Bragdon, M.D., Shulgina, Y., Cha, A., Wunderlich, Z., Gunawardena, J., DePace, A.H., 2019. Dissecting the sharp response of a canonical developmental enhancer reveals multiple sources of cooperativity. Elife 8. doi:10.7554/eLife.41266.

Parker, D.S., White, M.A., Ramos, A.I., Cohen, B.A., Barolo, S., 2011. The cis-regulatory logic of hedgehog gradient responses: key roles for gli binding affinity, competition, and cooperativity. Sci Signal 4, ra38. doi:10.1126/scisignal.2002077.

Parsons, G.G., Tg, A.C., 1997. Mitotic Repression of RNA Polymerase II Transcription Is Accompanied by Release of Transcription Elongation Complexes 17, 5791–5802.

Perry, M.W., Bothma, J.P., Luu, R.D., Levine, M., 2012. Precision of hunchback expression in the drosophila embryo. Curr Biol 22, 2247–52. doi:10.1016/j.cub.2012.09.051.

Phillips, R., Belliveau, N.M., Chure, G., Garcia, H.G., Razo-Mejia, M., Scholes, C., 2019. Figure 1 theory meets figure 2 experiments in the study of gene expression. Annu Rev Biophys 48, 121–163. doi:10.1146/annurev-biophys-052118-115525.

Phillips, R., Kondev, J., Theriot, J., Garcia, H.G., 2013. Physical Biology of the Cell, 2nd Edition. Garland Science, New York.

Polach, K.J., Widom, J., 1995. Mechanism of protein access to specific DNA sequences in chromatin: A dynamic equilibrium model for gene regulation. J Mol Biol 254, 130–49.

Rapp, O., Yifrach, O., 2017. Using the mwc model to describe heterotropic interactions in hemoglobin. PLoS One 12, e0182871. doi:10.1371/journal.pone.0182871.

Rapp, O., Yifrach, O., 2019. Evolutionary and functional insights into the mechanism underlying body-size-related adaptation of mammalian hemoglobin. Elife 8. doi:10.7554/eLife.47640.

Raveh-Sadka, T., Levo, M., Segal, E., 2009. Incorporating nucleosomes into thermodynamic models of transcription regulation. Genome Res 19, 1480–1496.

Razo-Mejia, M., Barnes, S.L., Belliveau, N.M., Chure, G., Einav, T., Lewis, M., Phillips, R., 2018. Tuning transcriptional regulation through signaling: A predictive theory of allosteric induction. Cell Syst 6, 456–469 e10. doi:10.1016/j.cels.2018.02.004.

Samee, M.A., Lim, B., Samper, N., Lu, H., Rushlow, C.A., Jimenez, G., Shvartsman, S.Y., Sinha, S., 2015. A systematic ensemble approach to thermodynamic modeling of gene expression from sequence data. Cell Syst 1, 396–407. doi:10.1016/j.cels.2015.12.002.

Sayal, R., Dresch, J.M., Pushel, I., Taylor, B.R., Arnosti, D.N., 2016. Quantitative perturbation-based analysis of gene expression predicts enhancer activity in early drosophila embryo. Elife 5. doi:10.7554/eLife.08445.

Scholes, C., DePace, A.H., Sanchez, A., 2017. Combinatorial gene regulation through kinetic control of the transcription cycle. Cell Syst 4, 97–108 e9. doi:10.1016/j.cels.2016.11.012.

Schulze, S.R., Wallrath, L.L., 2007. Gene Regulation by Chromatin Structure: Paradigms Established in Drosophila melanogaster. Annual Review of Entomology 52, 171–192. doi:10.1146/annurev.ento.51.110104.151007.

Segal, E., Fondufe-Mittendorf, Y., Chen, L., Thastrom, A., Field, Y., Moore, I.K., Wang, J.P., Widom, J., 2006. A genomic code for nucleosome positioning. Nature 442, 772–8.

Segal, E., Raveh-Sadka, T., Schroeder, M., Unnerstall, U., Gaul, U., 2008. Predicting expression patterns from regulatory sequence in drosophila segmentation. Nature 451, 535–40. doi:nature06496[pii]10.1038/nature06496.

Segel, L.A., Slemrod, M., 1989. The quasi-steady-state assumption: a case study in perturbation. SIAM Review 31, 446–477.

Sepulveda, L.A., Xu, H., Zhang, J., Wang, M., Golding, I., 2016. Measurement of gene regulation in individual cells reveals rapid switching between promoter states. Science 351, 1218–22. doi:10.1126/science.aad0635.

Sharon, E., Kalma, Y., Sharp, A., Raveh-Sadka, T., Levo, M., Zeevi, D., Keren, L., Yakhini, Z., Weinberger, A., Segal, E., 2012. Inferring gene regulatory logic from high-throughput measurements of thousands of systematically designed promoters. Nat Biotechnol 30, 521–30. doi:10.1038/nbt.2205.

Sherman, M.S., Cohen, B.A., 2012. Thermodynamic State Ensemble Models of cis-Regulation. PLOS Computational Biology 8, 1–10. URL: https://doi.org/10.1371/journal.pcbi.1002407, doi:10.1371/journal.pcbi.1002407.

Shermoen, A.W., O’Farrell, P.H., 1991. Progression of the cell cycle through mitosis leads to abortion of nascent transcripts. Cell 67, 303–10.

Staudt, N., Fellert, S., Chung, H.R., Jäckle, H., Vorbrüggen, G., 2006. Mutations of the Drosophila zinc finger-encoding gene vielfältig impair mitotic cell divisions and cause improper chromosome segregation. Molecular biology of the cell 17, 2356–65. URL: http://www.ncbi.nlm.nih.gov/pubmed/16525017 http://www.pubmedcentral.nih.gov/articlerender.fcgi?artid=PMC1446075, doi:10.1091/mbc.e05-11-1056.

Struhl, G., Struhl, K., Macdonald, P.M., 1989. The gradient morphogen bicoid is a concentration-dependent transcriptional activator. Cell 57, 1259–73. doi:0092-8674(89)90062-7[pii].

Swem, L.R., Swem, D.L., Wingreen, N.S., Bassler, B.L., 2008. Deducing receptor signaling parameters from in vivo analysis: Luxn/ai-1 quorum sensing in vibrio harveyi. Cell 134, 461–73. doi:10.1016/j.cell.2008.06.023.

Tu, Y., 2008. The nonequilibrium mechanism for ultrasensitivity in a biological switch: sensing by maxwell’s demons. Proc Natl Acad Sci U S A 105, 11737–41. doi:10.1073/pnas.0804641105.

Vilar, J.M., Leibler, S., 2003. Dna looping and physical constraints on transcription regulation. J Mol Biol 331, 981–9.

Wang, F., Shi, H., He, R., Wang, R., Zhang, R., Yuan, J., 2017. Non-equilibrium effect in the allosteric regulation of the bacterial flagellar switch. Nature Physics 13, 710–714. doi:10.1038/nphys4081.

White, M.A., Parker, D.S., Barolo, S., Cohen, B.A., 2012. A model of spatially restricted transcription in opposing gradients of activators and repressors. Mol Syst Biol 8, 614. doi:10.1038/msb.2012.48.

Xu, H., Sepulveda, L.A., Figard, L., Sokac, A.M., Golding, I., 2015. Combining protein and mrna quantification to decipher transcriptional regulation. Nat Methods 12, 739–42. doi:10.1038/nmeth.3446.

Xu, Z., Chen, H., Ling, J., Yu, D., Struffi, P., Small, S., 2014. Impacts of the ubiquitous factor zelda on bicoid-dependent dna binding and transcription in drosophila. Genes Dev 28, 608–21. doi:10.1101/gad.234534.113.

Yamada, S., Whitney, P.H., Huang, S.K., Eck, E.C., Garcia, H.G., Rushlow, C.A., 2019. The drosophila pioneer factor zelda modulates the nuclear microenvironment of a dorsal target enhancer to potentiate transcriptional output. Curr Biol 29, 1387–1393 e5. doi:10.1016/j.cub.2019.03.019.

Zeigler, R.D., Cohen, B.A., 2014. Discrimination between thermodynamic models of cis-regulation using transcription factor occupancy data. Nucleic Acids Res 42, 2224–34. doi:10.1093/nar/gkt1230.

Zeng, L., Skinner, S.O., Zong, C., Sippy, J., Feiss, M., Golding, I., 2010. Decision making at a subcellular level determines the outcome of bacteriophage infection. Cell 141, 682–91. doi:10.1016/j.cell.2010.03.034.

Zinzen, R.P., Senger, K., Levine, M., Papatsenko, D., 2006. Computational models for neurogenic gene expression in the drosophila embryo. Curr Biol 16, 1358–65. doi:10.1016/j.cub.2006.05.044.

Zoller, B., Little, S.C., Gregor, T., 2018. Diverse spatial expression patterns emerge from unified kinetics of transcriptional bursting. Cell 175, 835–847 e25. doi:10.1016/j.cell.2018.09.056.

